# Novel pipeline for large-scale comparative population genetics Pipeline for population genetics

**DOI:** 10.1101/2023.01.23.524574

**Authors:** Samantha E. Majoros, Karl Cottenie, Sarah J. Adamowicz

## Abstract

As scientists continue to ask complex questions about biodiversity and deal with increasingly large amounts of data, there is a demand for new methods and computational developments to perform scientific analyses. Analytical pipelines and modules can provide a way to meet these demands and ensure reproducibility in scientific methods and analyses. The goal of this study was to create efficient, reproducible, reusable programming modules that are publicly available for future research. These modules were used to determine population genetic structure measures and compare these measures across species with different biological traits. The functionality of the modules is shown through a case study on Diptera (true fly) species from Canada and Greenland. We leveraged high-throughput DNA sequencing data from Northern areas, as it is a valuable resource and provides new opportunities to study the Arctic. Data were pulled from public databases (Barcode of Life Data System and Global Biodiversity Information Facility), as well as taxon-specific literature. The pipeline we developed in R includes fifteen modules, including modules to prepare and filter the data, calculate population genetic structure measures (e.g., F_ST_), and run a multiple regression. These modules can be easily adapted and applied to a diverse set of animal groups, geographic regions, and biological traits. Best practices were followed for pipeline development, and the modules were designed and tested to work for datasets of different sizes by providing multiple different analyses and filtering options. Biological results were also obtained for Diptera species. Habitat and larval diet were both significantly related to population genetic structure. Evidence of isolation by distance and a relationship between population genetic structure and both latitude and longitude were also found. Overall, this study has created efficient, reusable bioinformatics modules, and provided insight into the factors affecting population genetic structure in Northern fly communities.

**Author Summary:** The goal of our study is to provide researchers with a series of R programming modules that can be used to determine how genetically different populations of the same species are and see how these differences are influenced by biological traits of the species and other variables. The scripts provided are flexible, accessible, and provide researchers with a useful tool for answering questions about population genetics and dealing with large amounts of DNA sequencing data. To show the functionality of the modules, we performed a case study using fly species from Canada and Greenland. We chose to focus on Northern regions as these areas are undergoing significant changes, and as the climate shifts, so will Northern species communities and compositions. Our study revealed that habitat and larval diet were both significantly related to population genetic structure. We also found evidence of isolation by distance, suggesting that populations that are further apart are less genetically similar, as well as relationships with latitude and longitude. Through this study we not only provided some insight into the factors influencing population genetic structure in Northern fly communities but also provided efficient and reusable R programming modules that can be used by other researchers.

## 1 Introduction

As scientists continue to ask complex and urgent questions about our world and environments, and deal with increasingly large amounts of data, there is a demand for new methods and computational developments to perform scientific analyses [1–3].

Bioinformatics and computational science are becoming increasingly important to scientific research [3]. There is also a need to ensure reproducibility, which becomes more difficult as scientific analyses increase in complexity [1]. Scientific workflows and analytical pipelines can make this more manageable [4]. Pipelines allow for the process to be understood and repeated by others, improve the efficiency of research, allow for repeatability of methods, and provide a foundation upon which to build future research. With more scientists needing to use some element of bioinformatics and programming, there is a need for user-friendly and accessible pipelines [3].

A programming pipeline includes all steps of the analysis, from data filtering and formatting to specific analyses required by the research, to the export of statistical results and data visualizations. There are best practices to consider when creating a programming pipeline, such as those outlined in Reiter et al. [1], Shade & Teal [2], and Akalin [5] (Table 1). To make the pipeline reusable and analyses reproducible, all relevant information needs to be made available to future users [5], including the files and data used, the choices made at each step of the analysis, dependencies and version information, and the fully commented code [5]. Pipelines, with associated data and documentation, need to be accessible and findable and can be shared using GitHub or similar websites [5, 6].

**Table 1:**
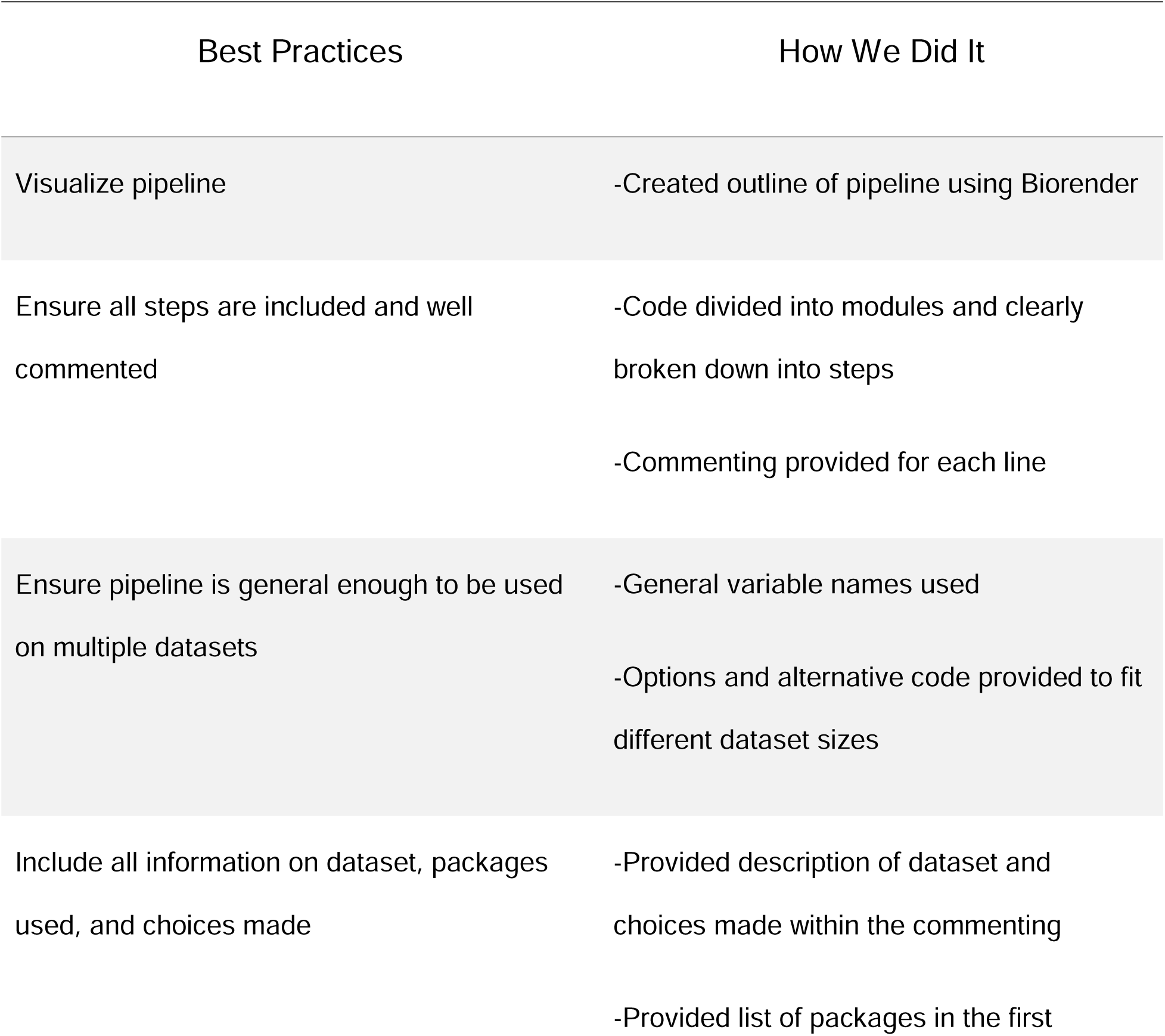
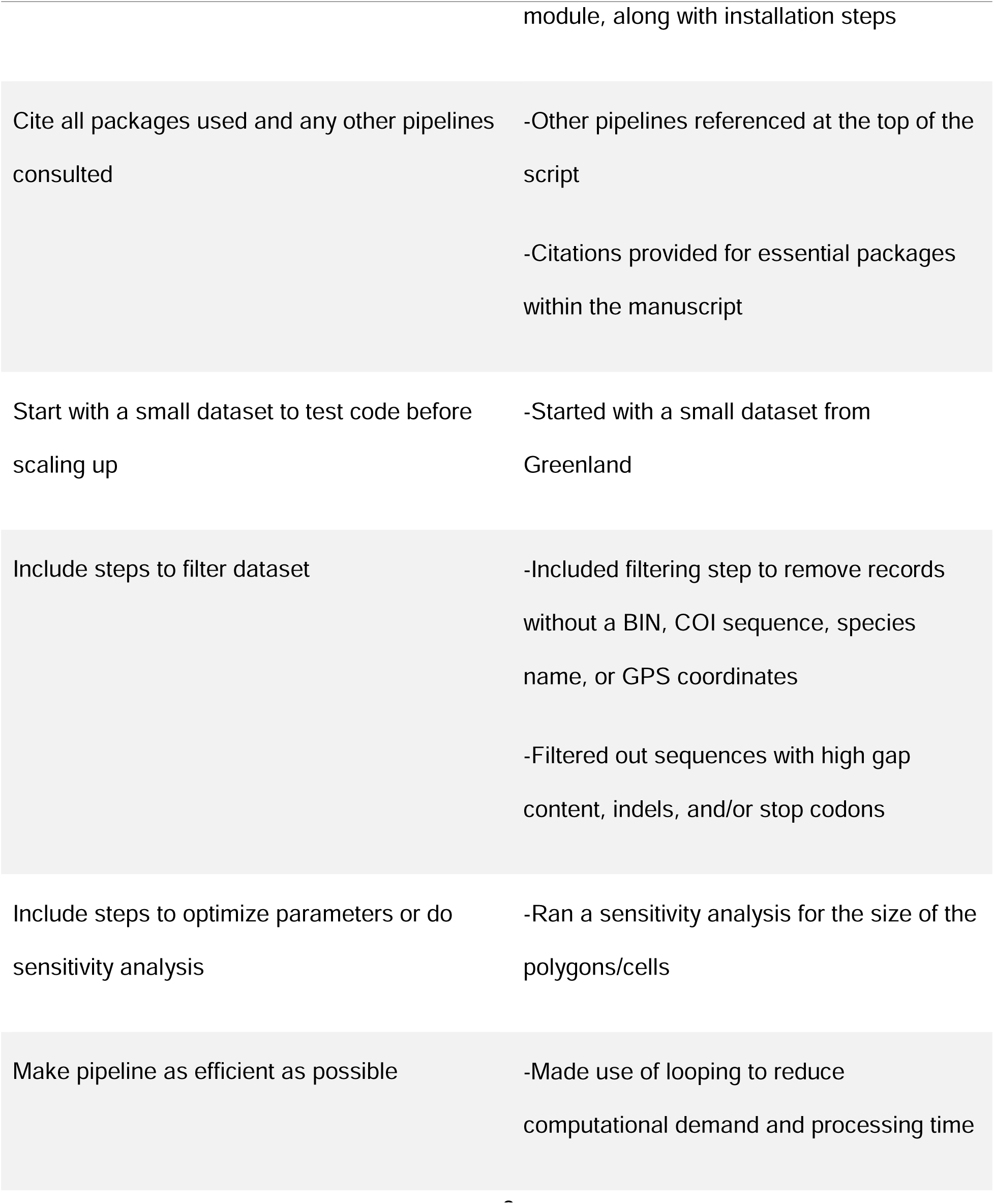
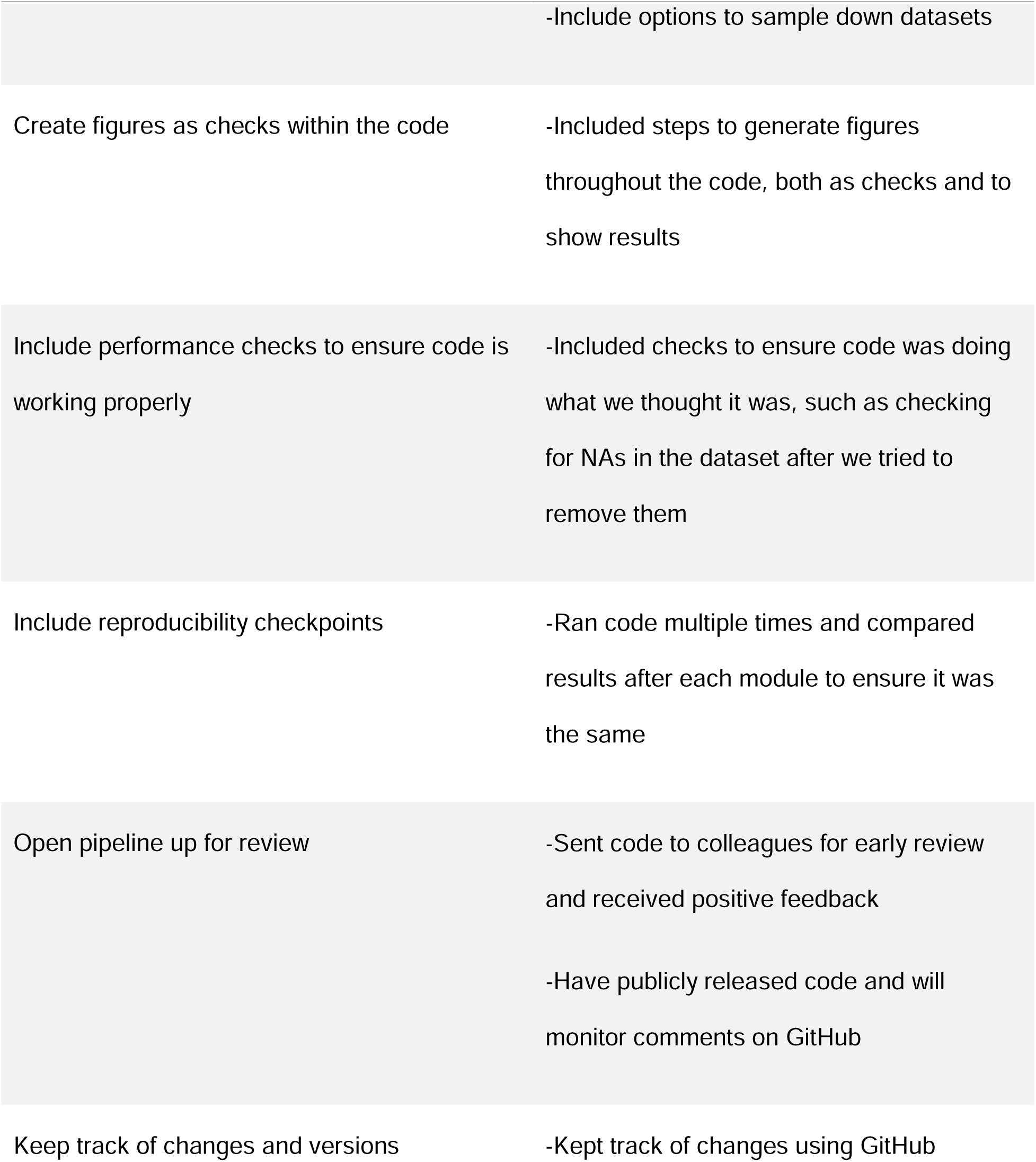
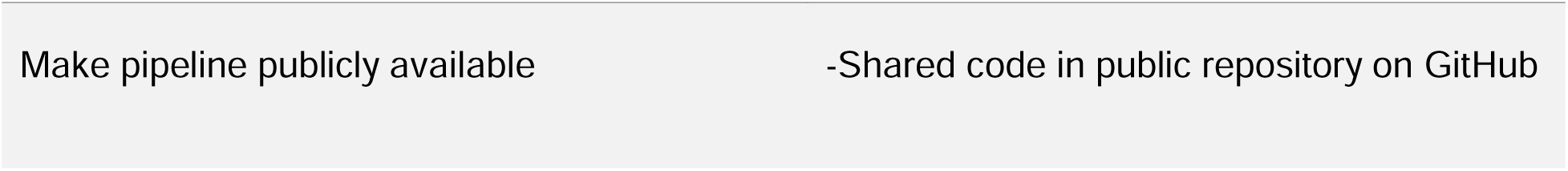
How each best practice, as outlined by Reiter et al. [1], Shade & Teal [2] and Akalin [5], was met during the creation of our pipeline.

Programming modules have been created for a variety of topics, including in the biological sciences, and many are publicly available on GitHub, such as the Landscape Genomics Pipeline [7], which investigates genetic structure and diversity. However, there are still many important questions that are unanswered, and more workflows and pipelines are needed. With increasing data availability and the continuous addition of new data to databases, there is value in making the analytical process easier and more automated. In this study, we selected, created, combined a series of modules and functions into a modular pipeline in the R programming language that can be used to answer biological questions regarding population genetic structure and realized dispersal. Understanding dispersal patterns can help us understand how communities and ecosystems function and how these communities will react to disturbance events, such as climate change or habitat fragmentation [8–9].

To develop programming modules that address biological questions, a thorough understanding of the biological background is needed. When developing a pipeline that focuses on realized dispersal and population genetic structure, it is important to understand the processes and the factors that influence them. Dispersal is the movement of one organism from one location to another and is important for many aspects of community ecology and evolution [8–9]. Dispersal rates, however, are difficult to calculate. Various methods are used to study dispersal, but the differences between these methods often lead to inconsistency in the data and results between methods [9]. Dispersal can be determined by direct methods, such as mark and recapture studies and in lab experiments [9–10]. Indirect methods include using the distribution of genetic diversity among populations to infer inter-population genetic differences and gene flow [9]. These studies can be time consuming and technologically difficult, and factors that contribute to dispersal rates (such as physiological and behavioural tradeoffs, life history traits, environmental conditions, and other factors influencing gene flow) are often unaccounted for using one method [10–11].

Dispersal as a general concept is the movement of organisms, but realized dispersal occurs when organisms are able to disperse and then reproduce and establish a population in a new location or leave descendants in an already established population. Indicators of relative realized dispersal rates, while still likely influenced by many factors, are more easily determined and may be important for determining the potential of species for successful colonization. Population genetic structure measures can be used as indicators of realized dispersal. Populations experience both genetic drift, which contributes to the genetic differentiation between populations, and gene flow, which contributes to the population connectivity and allows the exchange of genetic information. By calculating population genetic structure measures, we can investigate the balance between these two factors.

One potential set of factors that may influence realized dispersal is biological traits. Relationships between dispersal and traits have been found in prior studies and diverse taxa. For example, Fobert et al. [8] found that the larval phenotypes of marine fish species in Australia influenced their vertical placement in the water body and their dispersal ability. Steyn et al. [10] found similar results in different phenotypes of a fruit fly species. The relationship between dispersal and biological traits is also shown in salamander species with differing traits, where sympatric sexual species were better dispersers than unisexual species, likely due to higher locomotive endurance [12]. As a final example, successful invasion was found to be related to traits, such as dispersal ability, diet, reproductive rate, and environmental tolerance, in aquatic insect species by Carbonell et al. [13].

Other studies suggest that traits are not the most important predictor of successful dispersal. Investigating primary succession and colonization in plant species in Germany, Kirmer et al. [14] determined that the abundance of nearby species was more important than the traits of the colonizing species. Similarly, Downes & Lancaster [11] found that the traits of Australian insects associated with dispersal ability were not necessarily good predictors of colonization and that environmental conditions were most important. Öster et al. [15] also found that colonization was not related to functional traits of plant species in Sweden.

While many studies have investigated dispersal, most only considered differences between realized dispersal and colonization within a species or among a few closely related species or at a single location [8, 10–13,15]. There are still many taxa and locations that are understudied. However, the increased availability of georeferenced DNA sequence data provide an opportunity to advance knowledge across multiple taxa and geographic regions by semi-automating large-scale analyses. This study developed a novel way to study indicators of realized dispersal and population genetic structure across taxa and geographic regions. It thus provides the opportunity to discover patterns for understudied taxa and geographic regions, and lays the groundwork upon which to build further population genetic structure studies. This pipeline and set of modules infer relative realized dispersal rates, using population genetic measures calculated from DNA sequences as an indicator of realized dispersal, using molecular data. The case study used to demonstrate and test the pipeline makes use of publicly available DNA sequence data and uses cytochrome c oxidase subunit 1 (COI) sequences.

COI is a mitochondrial gene commonly used for DNA barcoding in animals and is good for identification of specimens to the species level and for species delineation and estimation of species richness using molecular methods [16–17]. DNA barcoding involves using one or more standardized gene regions to identify specimens to species and to discover new species [16, 18]. COI is expected to be more conservative than some other markers, such as microsatellites, which are commonly used for analysis below the species level in detecting differences among individuals within species, and has some disadvantages when compared to nuclear genes when determining phylogenetic relationships at deep nodes and investigating older demographic events [17, 19–20]. However, there is evidence that there is sufficient variability in COI to detect major population divisions and phylogeographic history, and COI has been shown to be comparable to nuclear genes for determining relationships between genera [17, 21–24]. Some other benefits of COI and mitochondrial genes include a haploid mode of inheritance, lack of introns in most taxa, and reduced exposure to recombination [16].

COI has been used to answer questions at a variety of temporal scales, including drawing conclusions about demographic history, divergence times, and events occurring millions of years ago [25–31]. Various studies have had success using COI to determine population genetic structure [19–20, 25–38]. This success was primarily due to COI’s high rate of evolution and mutation, as well as faster coalescence rate compared to nuclear markers, which makes it useful for determining genetic differences between isolated populations [19, 27, 30, 33, 38].

We wrote efficient, scalable, reproducible, and reusable modules, using the R language [39]. The R programming language provides a large variety of functions and packages for bioinformatics, ecological analysis, statistics, and visualization that allow analyses to be performed accurately and efficiently, with the source code available with open access. Due to R’s functionality, accessibility, the number of options it provides, and large community of contributors and users, it is a good environment and language in which to create analytical modules. These modules allow for accurate, reproducible, and efficient calculation of relative realized dispersal rates using population genetic structure (i.e., degree of genetic isolation between subpopulations) and geographic range as indicators of successful dispersal. The pipeline will also determine how the biological traits of species influence the indicators of population genetic structure.

Through this study, we produced a publicly available series of modules and commented code that aligns with best practices (as outlined in Akalin [5], Reiter et al. [1], and Shade & Teal [2]) that is available for other researchers to use.

To test the efficiency and accuracy of the modules, we ran an analysis on Diptera species from two relatively well-sampled Northern regions, Greenland and Canada [18, 40]. These regions were chosen due to the Northern and Arctic environments they contain. The climate is shifting, and Arctic areas, in particular, are undergoing a variety of changes at a rapid rate, including disappearing sea ice, melting permafrost, and increasing temperatures [41–45]. These changes affect the organisms present in the Arctic and can impact their survival and reproduction. By understanding the current population genetic structure patterns present in Arctic populations, we can potentially better understand how they will be impacted by the changing climate. Greenland provides an example of the minimal data needed to run the modules and as a check of module functionality. The Canada dataset acts as a case study and shows the module performance on a larger dataset. Diptera was chosen due to the abundance of data available on databases such as the Barcode of Life Data Systems (BOLD) [46] and Global Biodiversity Information Facility (GBIF) [47] and large barcoding efforts by programs such as the Global Malaise Trap Program [48–50]. Diptera is also a large, diverse order, possessing a variety of biological traits, including different diets, feeding modes, and habitat requirements [51], and information about selected ecological traits has been organized into a publicly available dataset [52].

The modules developed in this study allowed us to discover population genetic structure patterns for Diptera and will allow for future research to determine realized dispersal patterns for a wide range of taxa and environments. We determined relative realized dispersal rates for Diptera, with population genetic structure measures as an indicator, and compared these rates across species with differing traits. We hypothesized that indicators of realized dispersal rates will be influenced by the traits of organisms due to the importance of traits to the dispersal and colonization process [8, 10, 12–13]. We predicted that indicators of realized dispersal rates will be similar in species that share traits. While there is a large variety of traits that can be investigated, a select few were chosen for this example. We investigated habitat, adult diet, and larval diet [52]. We expected a generalist diet and terrestrial habitat to be associated with higher successful dispersal rates. Having flexibility of food choice may allow individuals of a given species to survive in a different environment more readily, and increased connectivity between terrestrial environments compared to aquatic may result in more dispersal and gene flow. In these cases, we would expect these populations to have lower genetic differentiation, due to high dispersal and successful colonization.

## 2 Methods

The pipeline and modules can be found in the repository Population_Genetic_Structure_Pipeline on Github (https://github.com/S-Majoros/Population_Genetic_Structure). This is a public repository containing the R script and necessary data files. The modules included in the pipeline and analysis are outlined below. A flowchart showing the steps of the pipeline can be found in Figure 1, and a brief description of each module is provided in Table 2.

**Figure 1:**
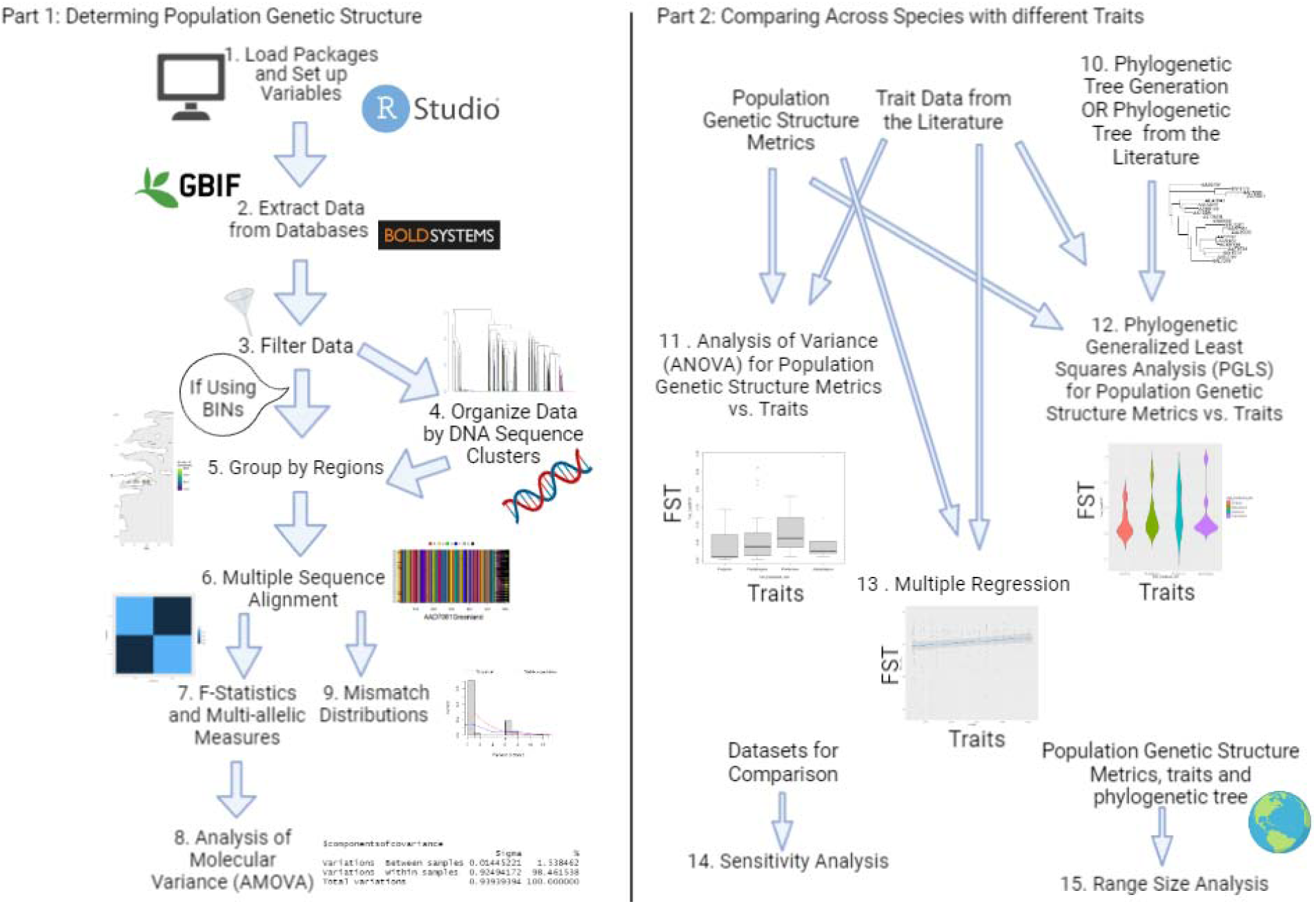
Steps in the population genetic strutcure pipeline. The numbered steps correspond to sections of code.

**Table 2:**
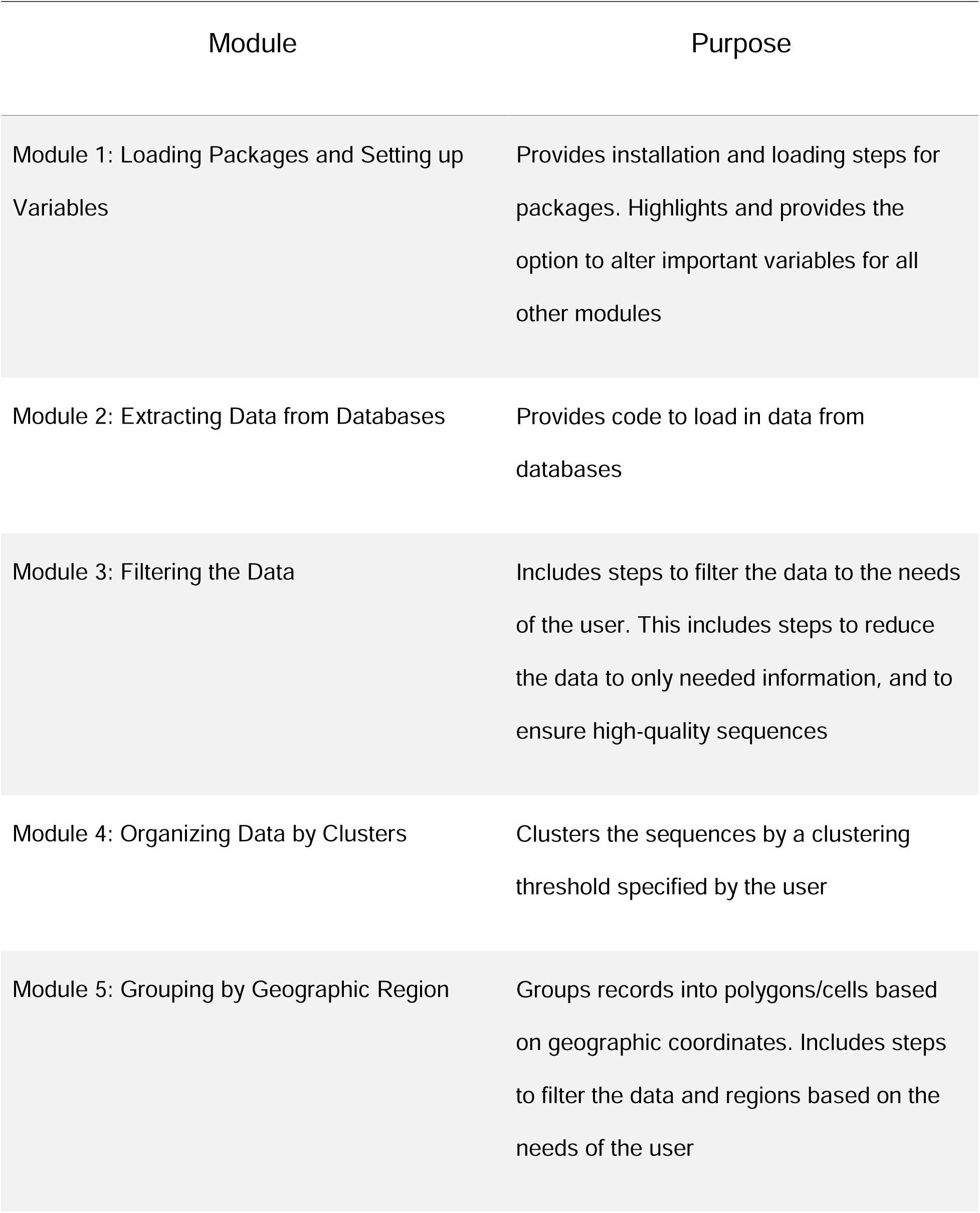
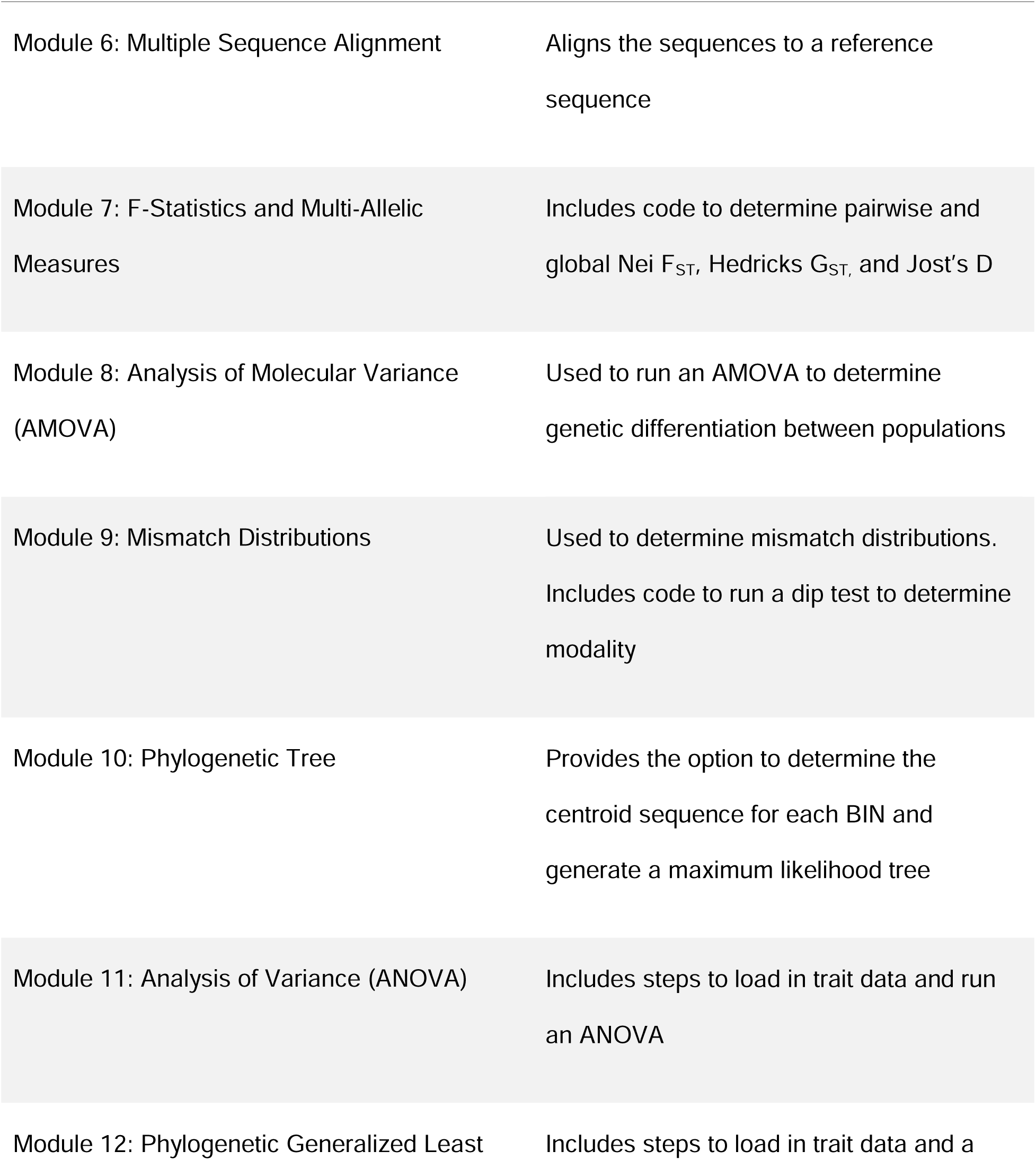
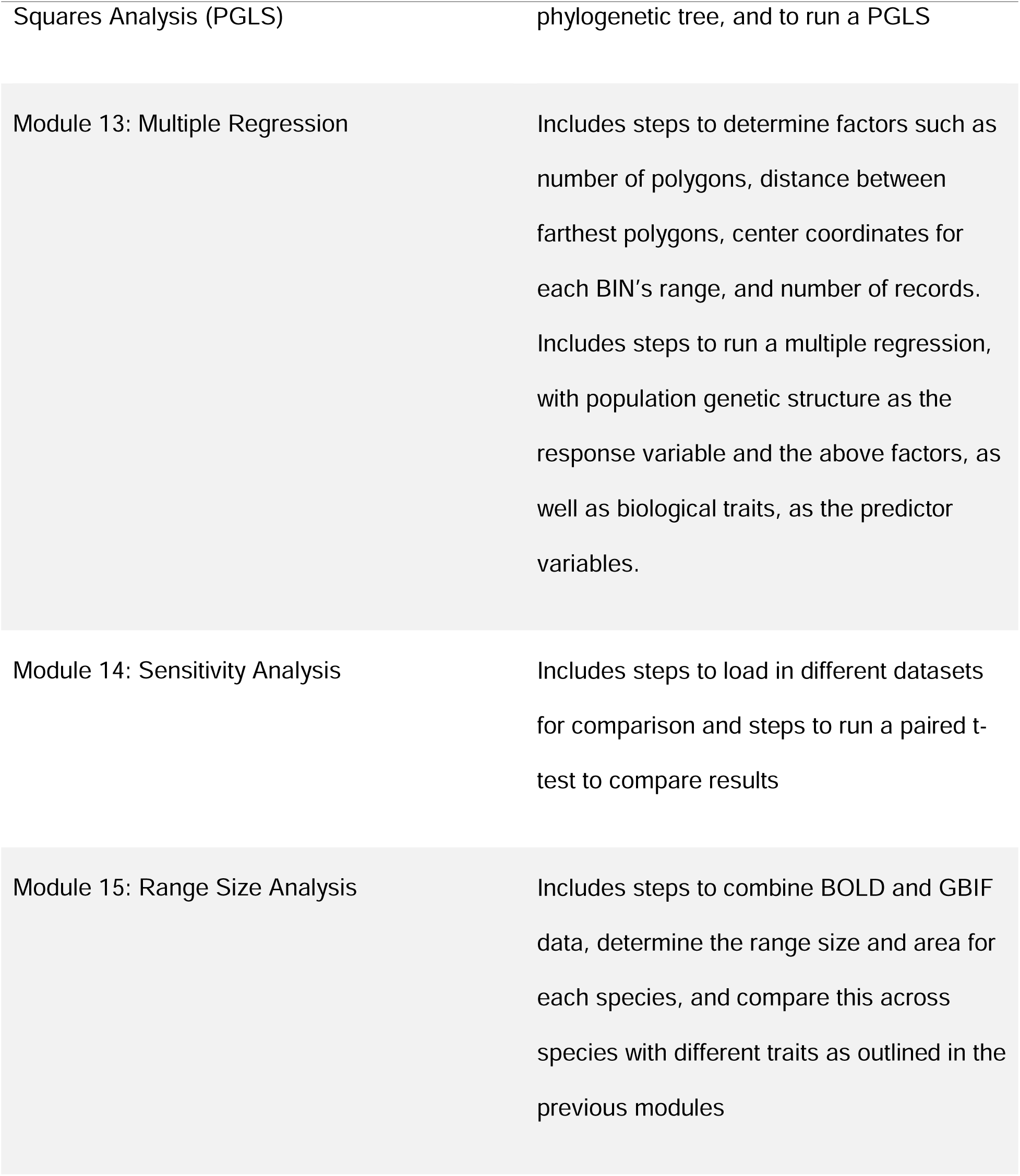
Brief description of each module and its purpose.

### 2.1 Loading Packages and Setting up Variables (Module 1)

Module 1 acts as a precursor for all the other modules and contains all the needed packages and parameters to run each module. All packages and parameters are separated by module and can be run together or individually. Important parameters are outlined in this module and made alterable by the user. When a variable or parameter appears in italics throughout this method section, it can be adjusted by the user in this module. This is to draw attention to these important parameters and allow easy adjustment, but other parameters and aspects of the script can also be edited as needed within each module.

### 2.2 Extracting Data from Databases (Module 2)

The main dataset needed to run the full population genetic structure analysis has to include nucleotide sequences and geographic coordinates, as well as species identification if performing the trait-based analysis. These data were all provided by the Barcode of Life Data Systems (BOLD) [46] database. Taxon-specific trait data are also required for Modules 11-13, and 15, and were obtained from the literature and loaded in during a later module [52]. Additional datasets, such as the Global Biodiversity Information Facility (GBIF) [47] dataset, can be used to provide additional geographic distribution information for the species. Geographic regions where there has been an effort to gather DNA barcode sequence data were emphasized for the case study, as such data are often associated with GPS information and other metadata [46].

Data were retrieved from BOLD. Data for Diptera from Canada and Greenland were downloaded on June 24th, 2021, using geographic keywords. As of Dec. 15th, 2021, BOLD contained 3,118,612 records of Diptera representing 27,440 species and 132,855 Barcode Index Numbers (BINs) [53]. BINs are operational taxonomic units (OTU) used by BOLD that are clusters of barcode sequences similar to species [53]. Data were pulled directly into the R environment using BOLD’s application programming interface (API). Data were also retrieved for Diptera from Greenland from GBIF [54].

These data included Diptera occurrences from Canada and Greenland, and only occurrences with geographic coordinates were retained.

### 2.3 Filtering the Data (Module 3)

The BOLD data include a variety of attributes and information, including the process and record IDs specific to BOLD, BIN numbers, nucleotide sequences, catalog, storing and collection information, taxonomic information, and geographic coordinates. Some records may contain further information about each specimen, including sex, lifestyle, reproduction, and habitat, but these are not always provided. For this analysis, the dataset was reduced to only the information needed for the modules, which included the process ID, BIN, family, genus, and species name, nucleotide sequence, and geographic coordinates. The BOLD records were filtered to exclude those without a BIN, COI nucleotide sequence of at least 500 base pairs (bp), species identification, or coordinates. BINs were used to represent species. Sequences with a high gap/N content (areas where there are deleted bases or missing information) were also removed. A threshold of 1% gap/N content was chosen because species often differ by more than 2% COI divergence [53]. Filtering out records with > 1% N and gap content is likely to yield a high-quality data set, given typical patterns of variability in COI in animals. The sequences were also checked for stop codons and non-biological insertions or deletions (indels) using the R package coil version 1.2.3 [55] and removed if these features were detected. The GBIF data were also filtered to remove occurrences without a species-level identification.

### 2.4 Organizing Data by Clusters (Module 4)

As mentioned above, BINs can be used to represent species in this study, but the module will also provide the option to use a different *clustering method* or *genetic distance threshold*. In our example, we used a 4% single linkage clustering threshold to group closely related sequences together into clusters. Lin et al. [56] determined that a threshold of 4-5% was suitable for clustering within the Diptera family Chironomidae.

Similarly, Baloğlu et al. [57] found stable results in Chironomidae across 3-5% clustering thresholds. Based on these results, 4% was deemed suitable for this study, but the threshold can be altered depending on the taxa being investigated. This option is important because, while BINs often correspond to species, the degree of concordance varies across taxa. By providing an alternative, users can more accurately assign specimens to species-like groupings across a variety of taxa or could use species names in the case of taxa with well-developed taxonomy and extensive taxonomic annotations of sequences. These results were then compared to those generated using BINs to understand how the two clustering approaches influence the results. To cluster the sequences, they were first aligned using the R package Decipher version 2.26.0 [58], using the default parameters, and a *reference sequence*, to which the sequences were aligned and trimmed. The reference sequence was a Diptera (*Patelloa xanthura)* sequence retrieved from BOLD (Process ID: ACGAZ1590-12, BIN:AAA1222). Users can upload their own reference sequence or choose one of those provided for some commonly studied taxa. The BIN was chosen because it met the following selection criteria: from the order Diptera, contained at least 10 COI-5P sequences, had at least one specimen photograph that matched the species-level taxonomy, and did not have taxonomic conflicts at the family level or above. The reference sequence within the selected BIN was chosen because it is 658 base pairs long, has 2 high-quality trace file chromatograms, and no missing nucleotides or stop codons.

A distance matrix was created using the model p-distance. This model was chosen as p-distance is used for the BIN algorithm, and keeping the distance model consistent allows for easier comparison to the BIN-based analysis. The sequences are then clustered using any method using the Decipher package. We chose the single linkage clustering method, since refined single linkage is used in Ratnasingham & Hebert [53] in the formation of BINs. To deal with computer memory constraints and reduce computing time, an option is included to randomly sample down the dataset to a maximum size for each BIN. This option is important as it allows the modules and script to be run by different researchers and makes it accessible to those who may not be able to access high-performance computing resources. If downsampling, users may consider repeating the randomization multiple times and comparing the results. This downsampling step was only used for the Canada data. A size of 100 *sequences per BIN* was chosen as the default setting to reduce strain on computer memory while still having a large sample size for analysis. Throughout this step, the data are kept together if the dataset is small, but an option is provided to separate the data by genus until after the clustering step, as looping through the genera individually during the clustering step reduces computational demands for large datasets.

### 2.5 Grouping by Geographic Region (Module 5)

Populations were geographically defined by grouping the records into polygons using the package dggridR version 3.0.0 [59] and the geographic coordinates provided for each record. *Polygon length* is a variable that can be set at the beginning of the analysis. The grid and polygons that are used for this analysis are hexagonal in shape. Using hexagons has several benefits when compared to using a traditional rectangular grid [60]. Hexagons allow for all points on the edges to be closer to the center than they would be in a rectangle and are able to form a grid that covers a whole area [60]. They are also less distorted by the curvature of the earth and have benefits when it comes to visualizing the grid [60]. The data were then filtered further to include BINs only if they meet the minimum sample size requirement of at least 20 sequences total and to be present in at least two polygons. Each BIN within each polygon was also required to have at least 10 sequences in the polygon. These filtering steps were included in order to ensure that each BIN included a suitable amount of data for population genetic analysis. Polygons were filtered out if they contained fewer than 10 BINs that met the above sample size criterion. Accumulation curves and abundance and incidence-based Chao richness estimators [61] were calculated for each polygon. For our example, polygons that have at least 50% of their *expected BIN richness sampled* were included in the analysis. While having a higher richness is preferable, this may not be possible depending on the geographic regions studied. This filtering step is included in order to ensure the polygons being included in the analysis are reasonably well sampled, to yield a more general, rather than BIN-specific, analysis of population genetic structure in Diptera.

### 2.6 Multiple Sequence Alignment (Module 6)

Once the dataset was filtered, the sequences were aligned again within each BIN. The sequences were trimmed using the *reference sequence* and then aligned using the same parameters outlined in the clustering step. If BINs were used, this was the first multiple sequence alignment for the dataset. If clusters were used, this was the second, but this time only the clusters included in the final analysis were included.

### 2.7 Population Genetic Structure Measures (Modules 7 - 9)

Measures of relative realized dispersal (population connectivity), pairwise Nei’s F_ST_ [62], Hedrick’s G_ST_ [63], and Jost’s D [64], were calculated using the package mmod version 1.3.3 [65] (Module 7). These measures quantify how genetic diversity is distributed among sub-populations and determine levels of genetic differentiation (Fig.2). These measures differ slightly in the way they are calculated [66]. Values of F_ST_ can range between 0 and a value determined by the heterozygosity in the data, while G_ST_, and Jost’s D use a standardized range of 0 – 1 [66], with zero indicating identical allele frequencies among populations and 1 indicating no sharing of alleles. The original F_ST_ was for two-allele data, but Nei’s F_ST_ is adjusted for multi-allelic data. The measures are calculated for each polygon pair for each BIN. For larger datasets, an option to calculate the global F_ST_, G_ST,_ and Jost’s D is included. This provides one measure for each BIN. This can also be calculated for small datasets, though if fewer polygons and BINs are included, it is more feasible to examine the results between each polygon pair. An analysis of molecular variance (AMOVA) was then used (using the package poppr version 2.9.3 [67]) to take into account the amount of genetic difference among alleles and quantify how the genetic variability is partitioned among geographically hierarchical populations (Module 8). The default parameters of this function were used, and raw pairwise distances are calculated from the data. Similar to the population genetic structure measures, the AMOVAs give us an idea of the genetic differentiation between populations. Mismatch distributions were also measured as an additional measure of population expansion, using the package pegas version 1.2 [68], with our example study targeting northern regions (Module 9). Mismatch distributions are used to investigate patterns in the distribution of pairwise genetic distances between sequences (as in Shum & Palumbi [36]) [69]. In order to determine whether the distribution is uni or multimodal, a dip test was performed [70]. The null hypothesis is a unimodal distribution. If the distribution is unimodal, this suggests that the population is still expanding.

**Figure 2:**
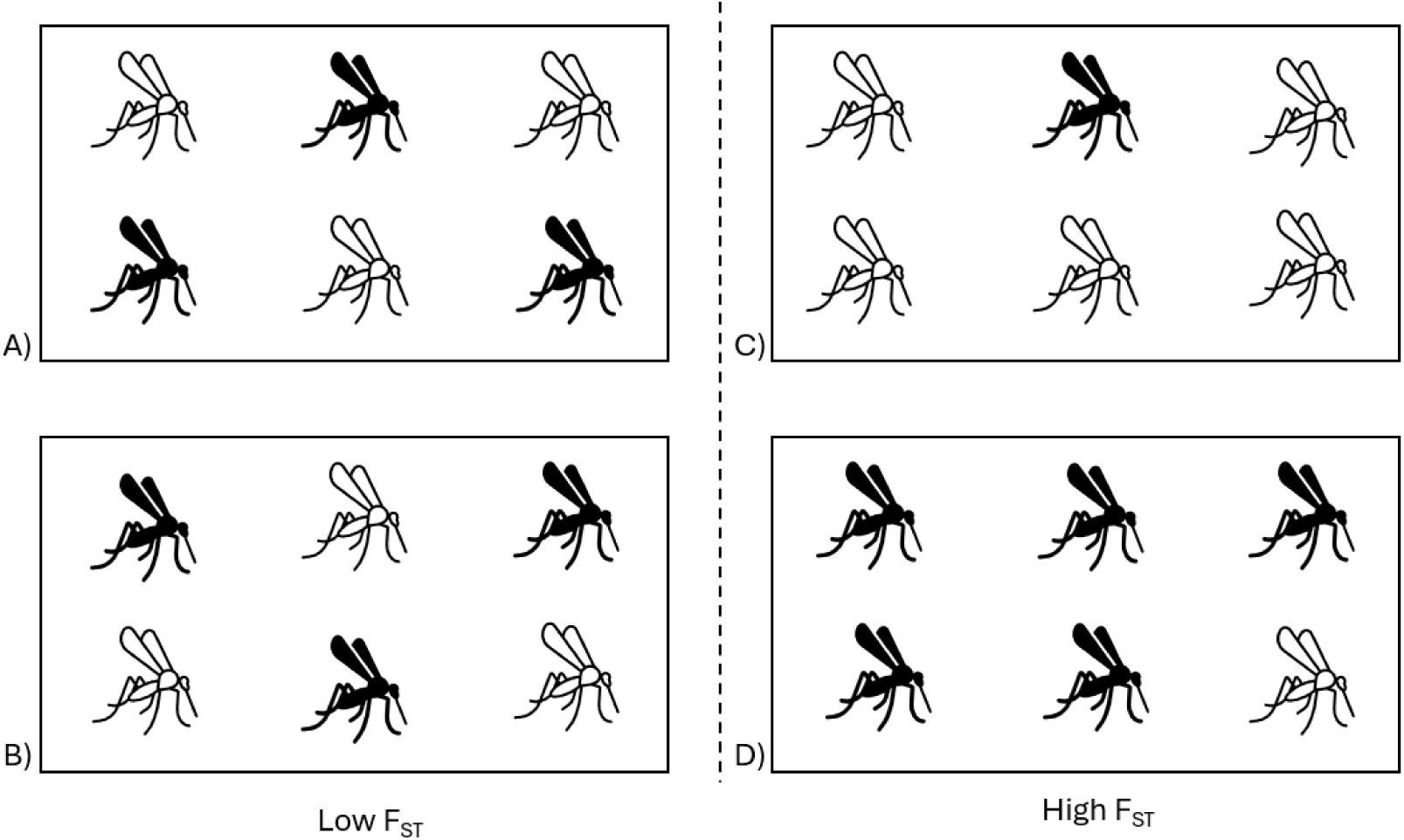
Conceptual illustration of low vs high F_ST_. The colours represent different alleles (haplotypes) of COI within the same species. Panels A & B show low F_ST_. The two populations share the same allele frequencies. Panels C & D show high F_ST_. While the global allele frequencies are the same as the other panels, in this case the two sub- populations have very different allele frequencies, indicating strong population genetic differentation, due to factors including the amount of gene flow, historical colonization patterns, and genetic drift.

### 2.8 Phylogenetic Tree (Module 10)

The phylogenetic tree module provides the option to generate a maximum likelihood tree from the user’s data, rather than uploading a tree from the literature. A well-supported, multi-gene tree may not be available from the literature for all study taxa, so the option to generate one de novo allows for the phylogenetic generalized least squares (PGLS) analysis to be performed for taxa for which the evolutionary relationships are not yet well known. The maximum likelihood tree was constructed using one representative sequence from each BIN or cluster, the centroid. A centroid sequence is defined as the sequence with the minimum average distance to all others in its BIN (or species cluster) [71]. Alignments for each BIN or cluster were performed using the same process as outlined above, and distances were found using the default parameters of the function dist.dna from the package ape version 5.7 [72]. The centroid sequences (one sequence for each BIN or cluster) were then aligned to the *reference sequence and the outgroups*. For this case study, three sequences from Trichoptera were chosen as outgroups. Trichoptera was chosen as it is closely related to Diptera, but definitively outside the ingroup [73]. Before creating the tree, the best-fit model of nucleotide evolution was estimated using the *Bayesian Information Criterion (BIC) score*. The tree was then generated using the function optim.pml from phangorn version 2.11.1 [74] and the following parameters: intervals of *discrete gamma distribution (the k-value)* was set to 4, and optNni, optGamma, and optInv were set to true. The proportion of invariant sites was determined based on the model, and a neighbour-joining (NJ) tree [75] (created using NJ from phangorn version 2.11.1 [74]) was used as the guide tree.

### 2.9 Relationship between Traits and Population Genetic Structure (Modules 11 – 13)

To determine the relationship between traits and population genetic structure, three commonly used methods were used. BINs or clusters were grouped by traits, and the population measures (F_ST_, G_ST_, Jost’s D) were compared across groups with different biological traits using an analysis of variance (ANOVA) (Module 11). A PGLS analysis was also performed to take into account phylogenetic relationships (Module 12). For larger datasets, such as our Canada dataset, an option to run a multiple regression was included (Module 13). If other analyses are preferred by the researcher, these modules can be replaced, and different methods can be used.

For our case study, the traits investigated were habitat, adult diet, and larval diet.

Other researchers can substitute in other traits. Traits were assigned to each species/BIN using the dataset outlined in Majoros et al. [52] and is available on the Dryad Data Repository (https://doi.org/10.5061/dryad.fqz612jwx). These traits were obtained through literature searches and were assigned at the lowest taxonomic level possible. For BINs that contained multiple species names, the most common species name within the BIN was used. For habitat, terrestrial is defined as “taxa that live primarily in land habitats”, aquatic is defined as “taxa that live primarily in water bodies or associated habitats”, and semi-aquatic is defined as “species that primarily live in wet habitats and require high levels of moisture and some elements of both terrestrial and aquatic habitats” [52]. For diet, predaceous taxa are defined as “those who prey on insects or other animals”, mycophagous taxa are defined as “those who feed on fungi”, saprophagous taxa are defined as “those who feed on decaying organic matter”, parasites are defined as “those who live and feed in or on an organism of another species”, parasitoids are defined as “those whose young develop within a host, eventually resulting in host death”, leaf/root/stem feeding taxa are defined as “those who feed on the leaves, roots and/or stems of plants”, detritus and algae feeding taxa are defined as “those who feed on small particles of algae and detritus”, kleptoparasites are defined as “those who take food and resources from another species“, and nectar/pollen/honeydew feeding taxa are defined as “those who primarily feed on nectar, pollen and/or honeydew” [52]. Taxa can also be assigned a combination of the diet categories. If a species’ diet consisted of more than three of the above categories, they were included in the polyphagous category. As some flies are non-feeding in the adult stage, this was also recorded as its own category. Many higher taxonomic levels are diverse and contain species with different traits. In this case, the most common category was used. When this was unclear and multiple categories were found in the literature with the same frequency, multiple categories were included separated by “or”. In some cases, there was insufficient information available, based upon the literature searches conducted by Majoros et al. [52], to assign a species to a trait category. These species are marked as “unclear” for that trait.

For the PGLS, the required tree can be specified by the user or created using the phylogenetic tree step. For the case study, the topology of the phylogenetic tree was created manually in the program Mesquite [76]. This tree was based on the family-level Diptera tree created by Wiegmann et al. [77] using 14 nuclear loci and complete mitochondrial genomes, as well as morphology. Haseyama et al.’s [78] phylogenetic hypothesis, based on 4 protein-coding genes, was used as an additional lower-level taxonomic tree for the relationships within the family Muscidae.

For the Canada case study, a multiple regression was performed using the lm function from the stats package version 4.2.2 [39]. The global F_ST_, G_ST,_ and Jost’s D values were the response variables, and the predictor variables used for each BIN were habitat, adult diet and larval diet, the number of polygons each BIN was found in, the centre coordinates of each BIN range (found by creating a polygon and finding the centre coordinates using the package sp version 1.6.0 [79]), the distance between the centre points of the two most distant polygons occupied by a given BIN or cluster, and the number of records. Steps to calculate the linearized version of the measures are also included so these can be compared to the distance, latitude, and longitude.

### 2.10 Sensitivity Analysis (Module 14)

While there are too many parameters to determine the effects of each parameter on the analysis within the scope of this study, a key variable, polygon size, was chosen to illustrate the concept of a sensitivity analysis using the Diptera case study data. The size of the polygon/region influences the number of BINs and records within each region, so it is important to determine the effect it will have on estimated population genetic structure and choose a polygon size that is suitable. A range of polygon lengths was included from 50 to 500 km to see the effects of polygon size. In addition to the original 500 km analysis, the analysis was performed again using lengths of 50 and 250 km, and the global population genetic structure measure results were compared using paired t-tests.

### 2.11 Testing Association between Traits and Species Ranges (Module 15)

The relationship between biological traits and geographic range size was studied, using range size as the response variable. Range size in m^2^ for each species included in the analysis was determined using the coordinates provided by both the BOLD and GBIF data, and the package sf version 1.0.9. [79]. Only species found in both BOLD and GBIF, were used and only the coordinates of the individual records, not entire BINs, were used. The range sizes were then compared across species with different biological traits using the same steps outlined in previous steps. This analysis was only performed for the Greenland dataset.

## 3 Results

### 3.1 Pipeline and Module Development

A novel pipeline consisting of 15 modules was developed, 14 of which are used to determine population genetic structure measures as proxies for realized dispersal, and compare these measures across species with differing biological traits. The 15^th^ module is an example of using range size. Best practices were met during the development of these modules, as outlined in Table 1. The modules ran without error and without unnecessary computational demands, and the checks included in the script showed that the R script was working as intended. Using reproducibility checkpoints after each module also showed that the modules produced the same results after each run.

### 3.2 Case Study

#### 3.2.1 Population Genetic Structure Measures

Twenty BINs, representing 18 species, across 2 geographic regions were included in the analysis for Greenland (Fig. 3a; Table 3a). All the F_ST_ values for the BINs were low, showing low levels of genetic differentiation. The results for Hedrick’s G_ST_ and Jost’s D were more variable, with 6 BINs having a higher value (above 0.5) and 14 having a lower value for both measures. The AMOVA analysis revealed low levels of variation between the regions for all BINs. From the figures produced by the mismatch distribution analysis, 15 BINs’ populations appeared to be unimodal, and five appeared to be bimodal. However, the dip test statistics indicated that there was no significant multimodality, suggesting that all populations are still expanding. An example of a mismatch distribution is provided in Figure 4. The full analysis was completed again using a 4% clustering threshold instead of BINs. Eighteen clusters were included, and the results were similar to those of the BIN analysis. The full clustering analyses results are outlined in S1 Appendix.

**Figure 3:**
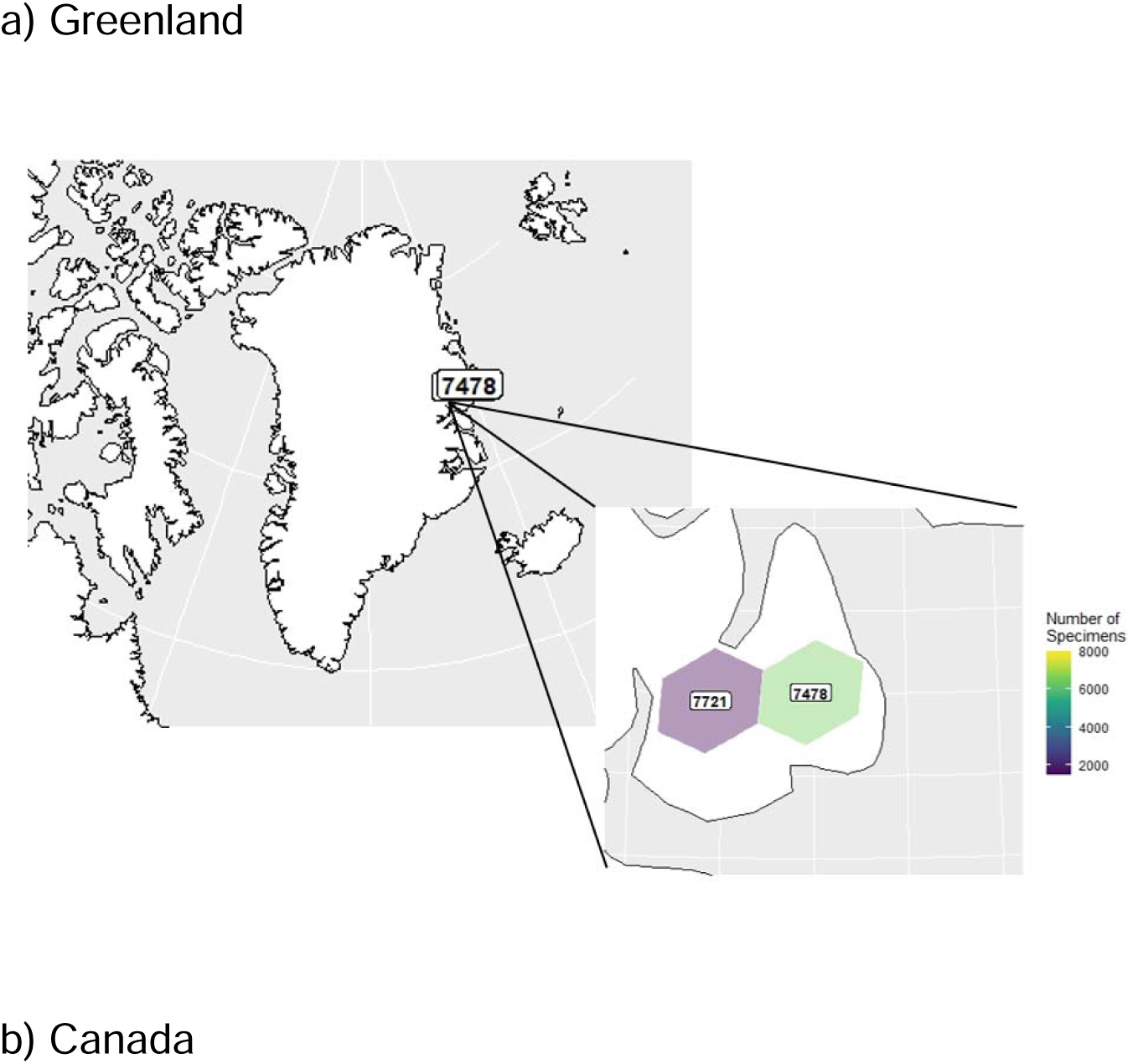
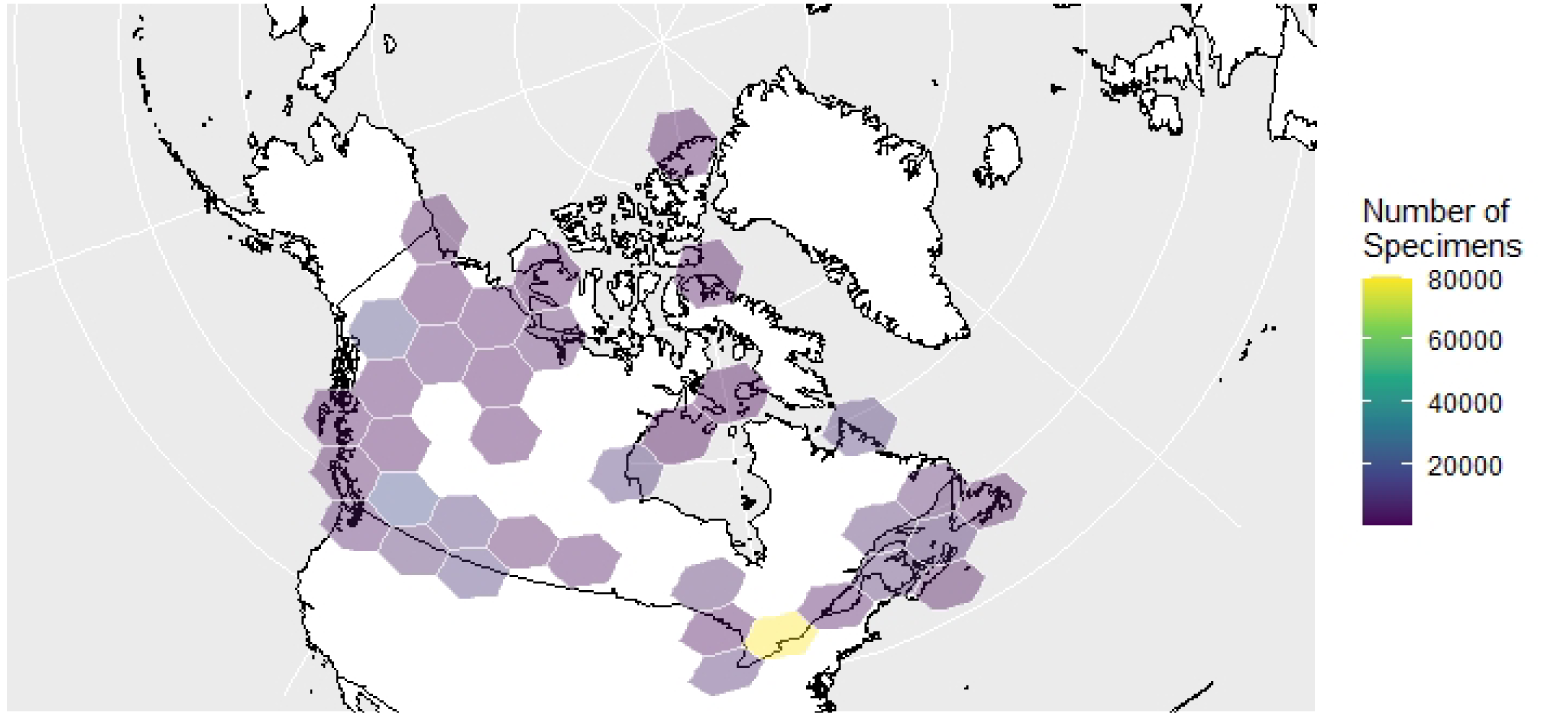
Regions and number of Diptera records for the case study BIN analysis in a) Greenland (polygon length = 50 km across) and b) Canada (polygon length = 500 km across). Regions labels were removed from the Canada map to avoid overcrowding.

**Figure 4:**
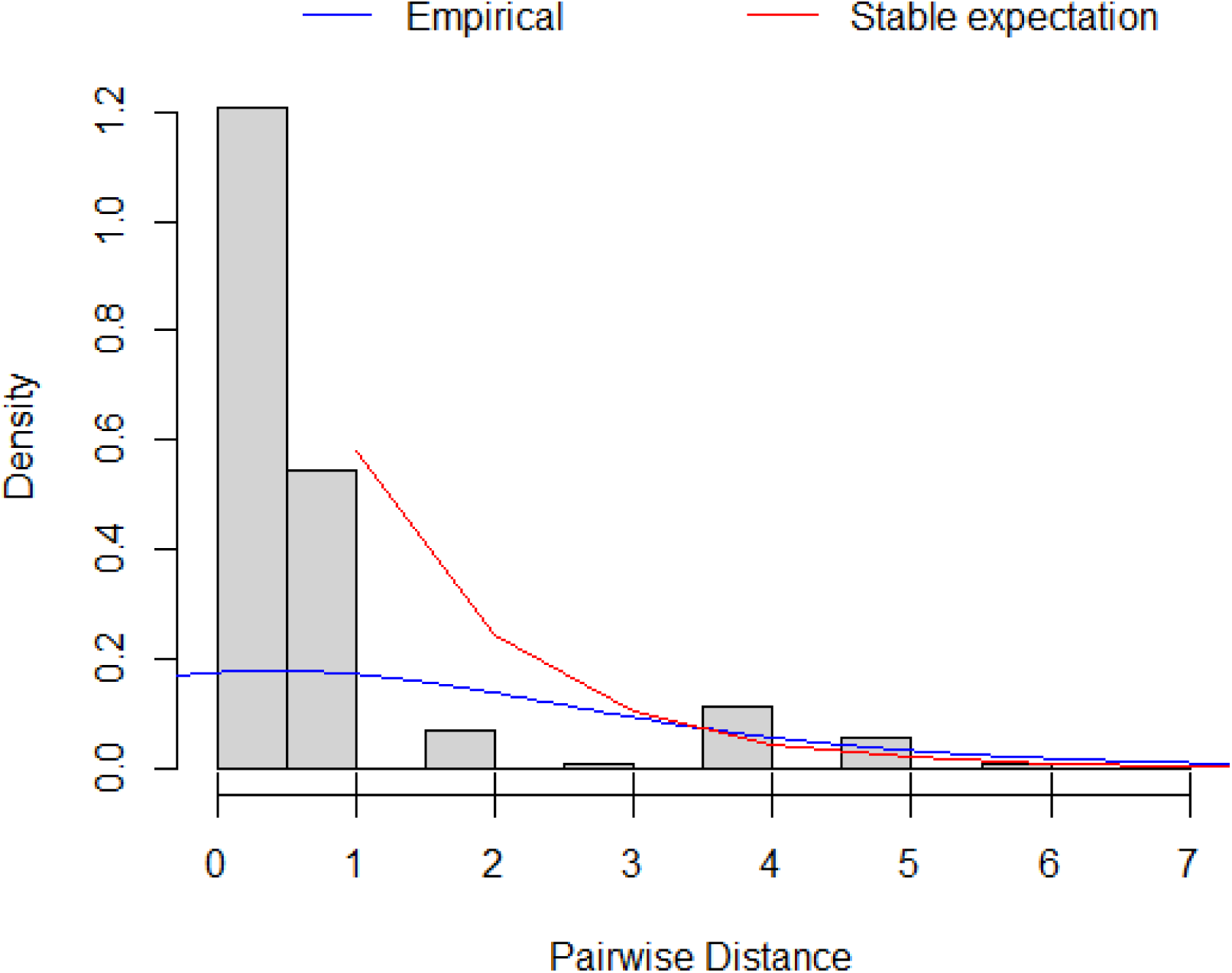
An example of a mismatch distribution for BIN BOLD:AAB0080. There were no significant signs of mulitmodality.

**Table 3:**
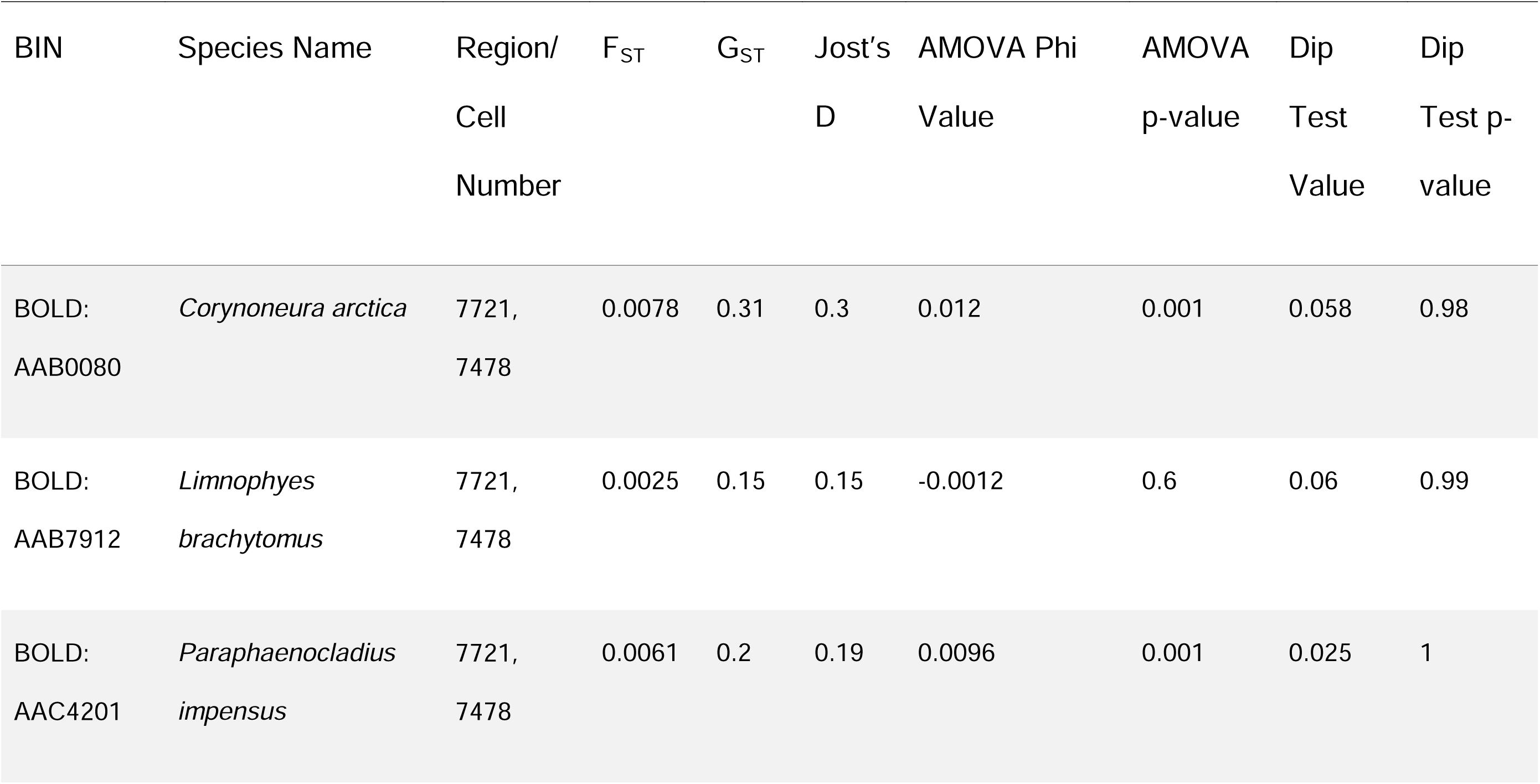
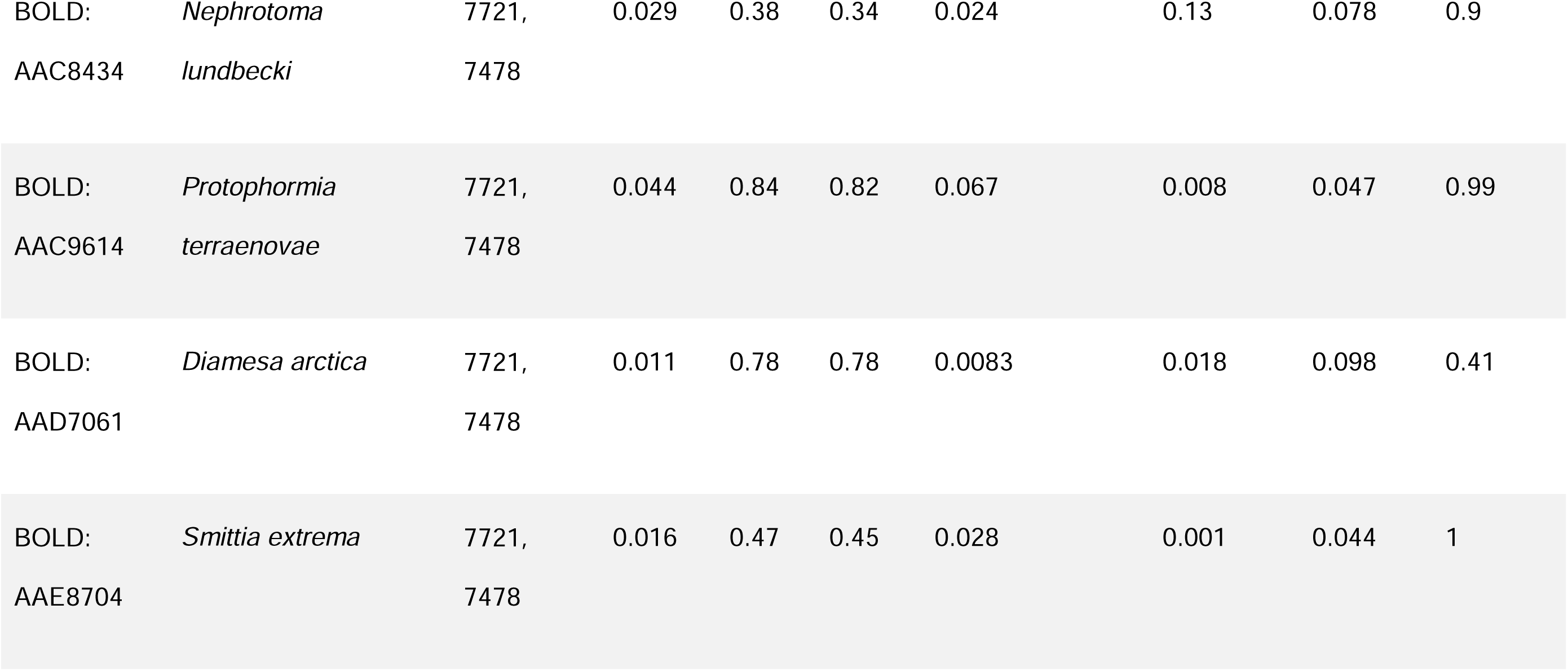
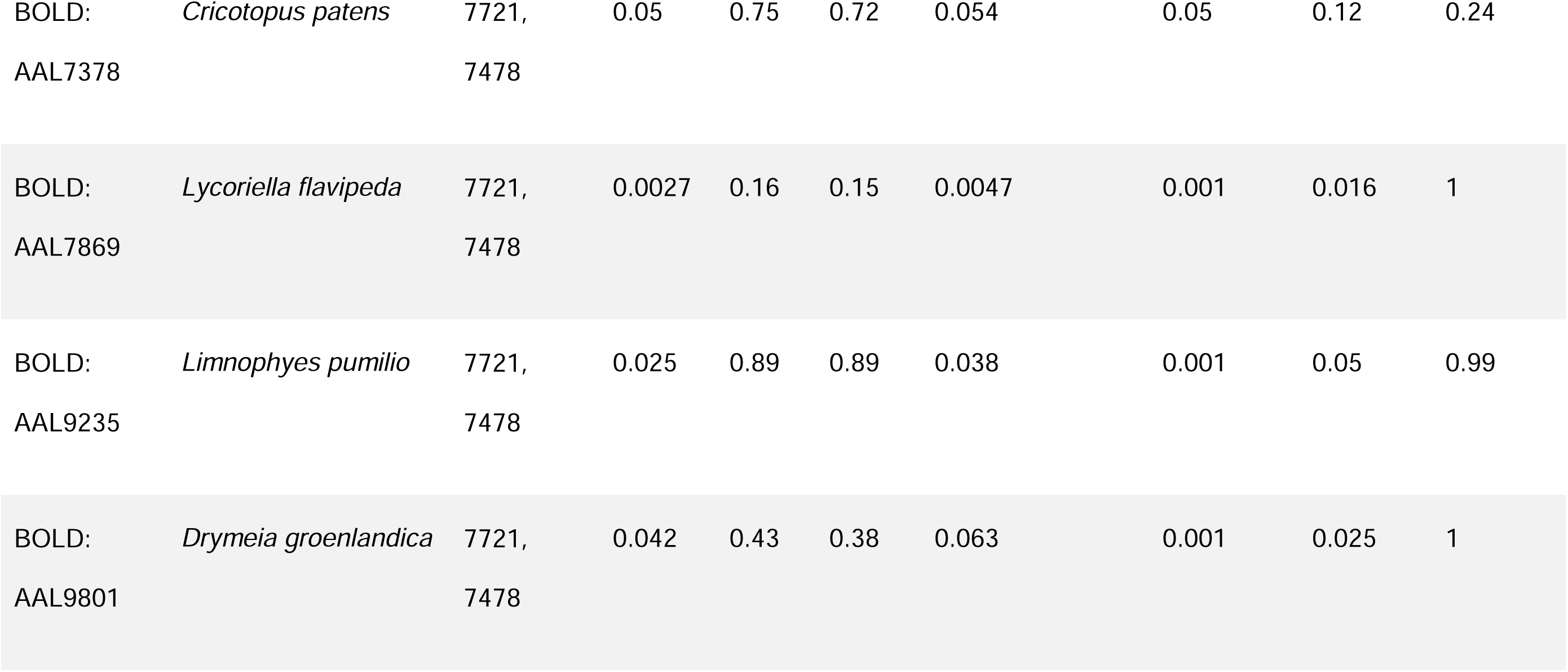
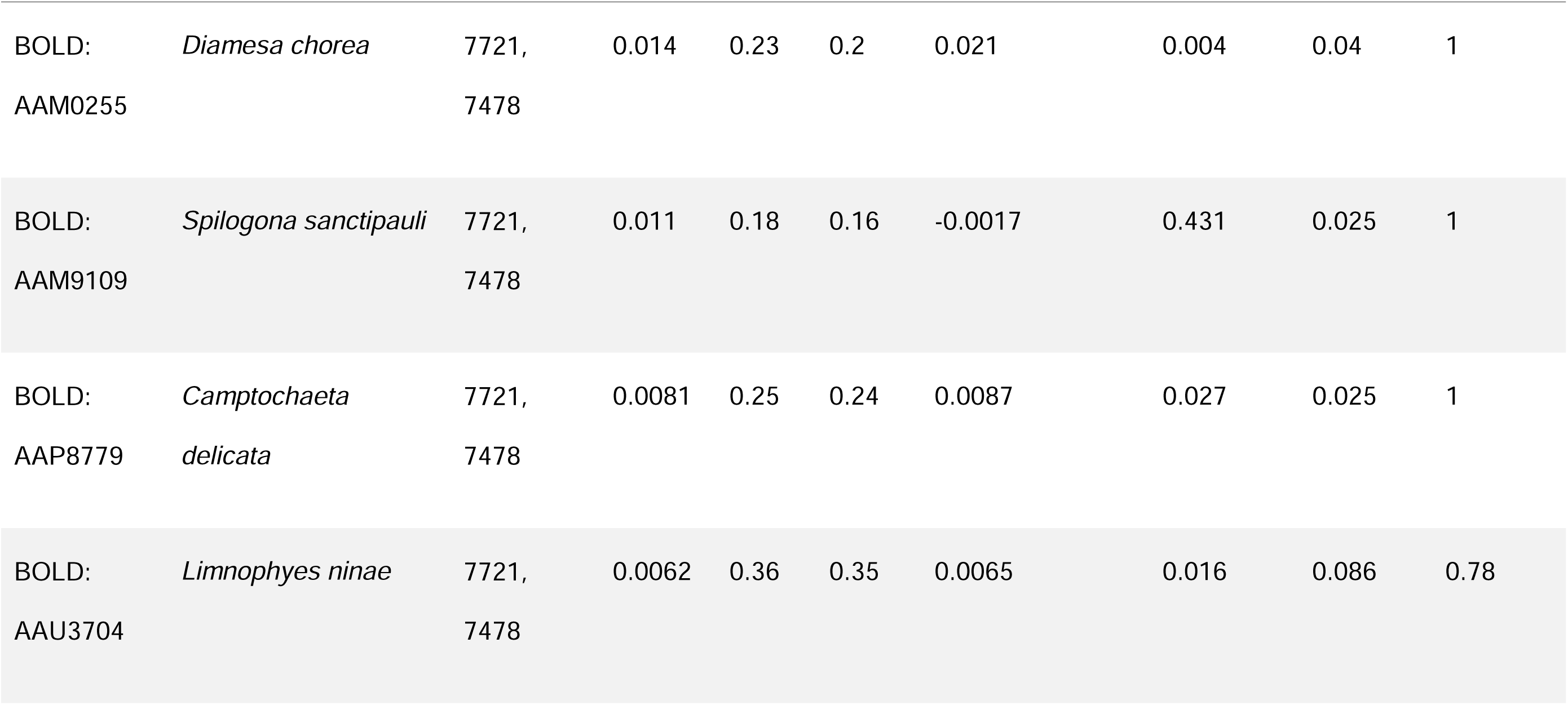
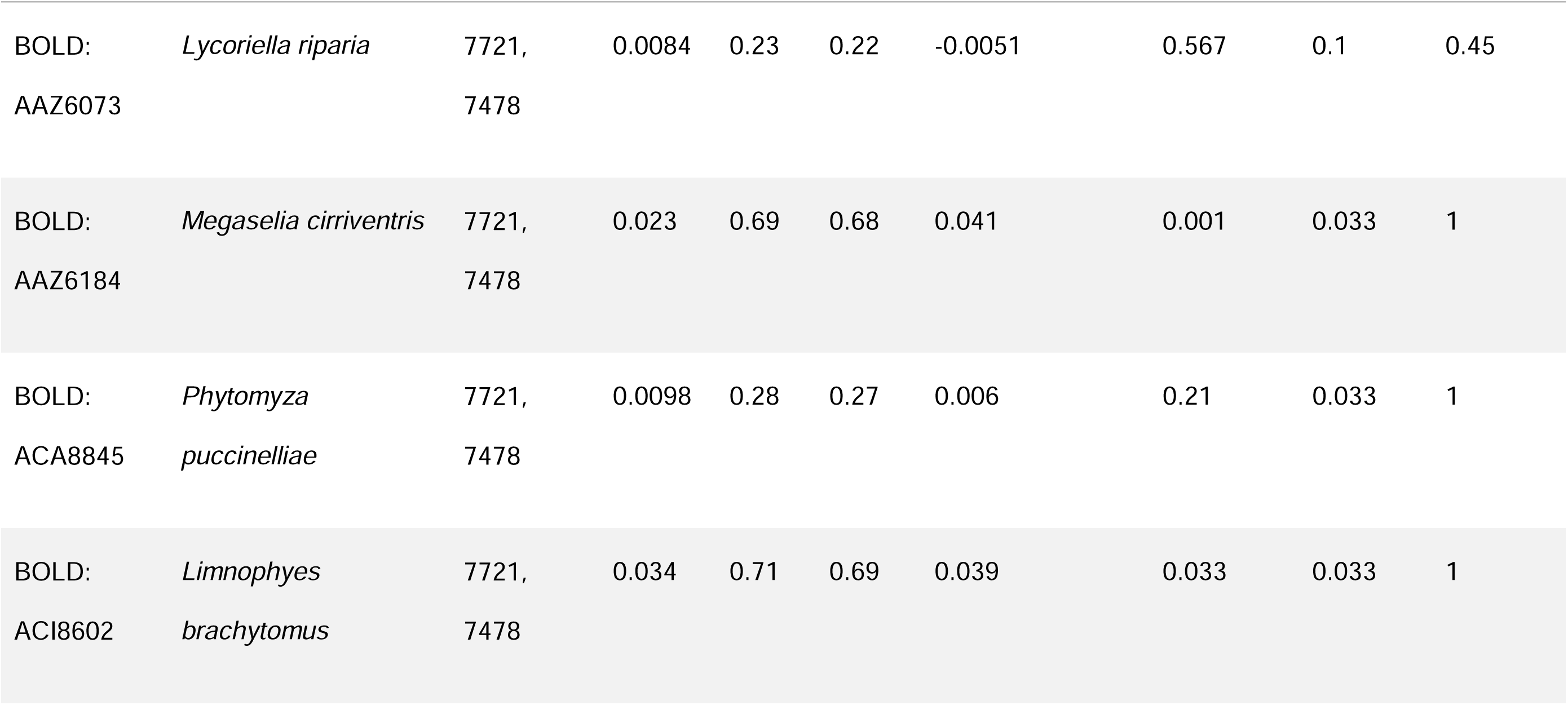
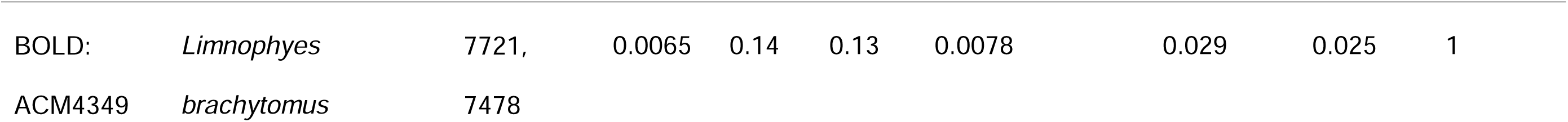
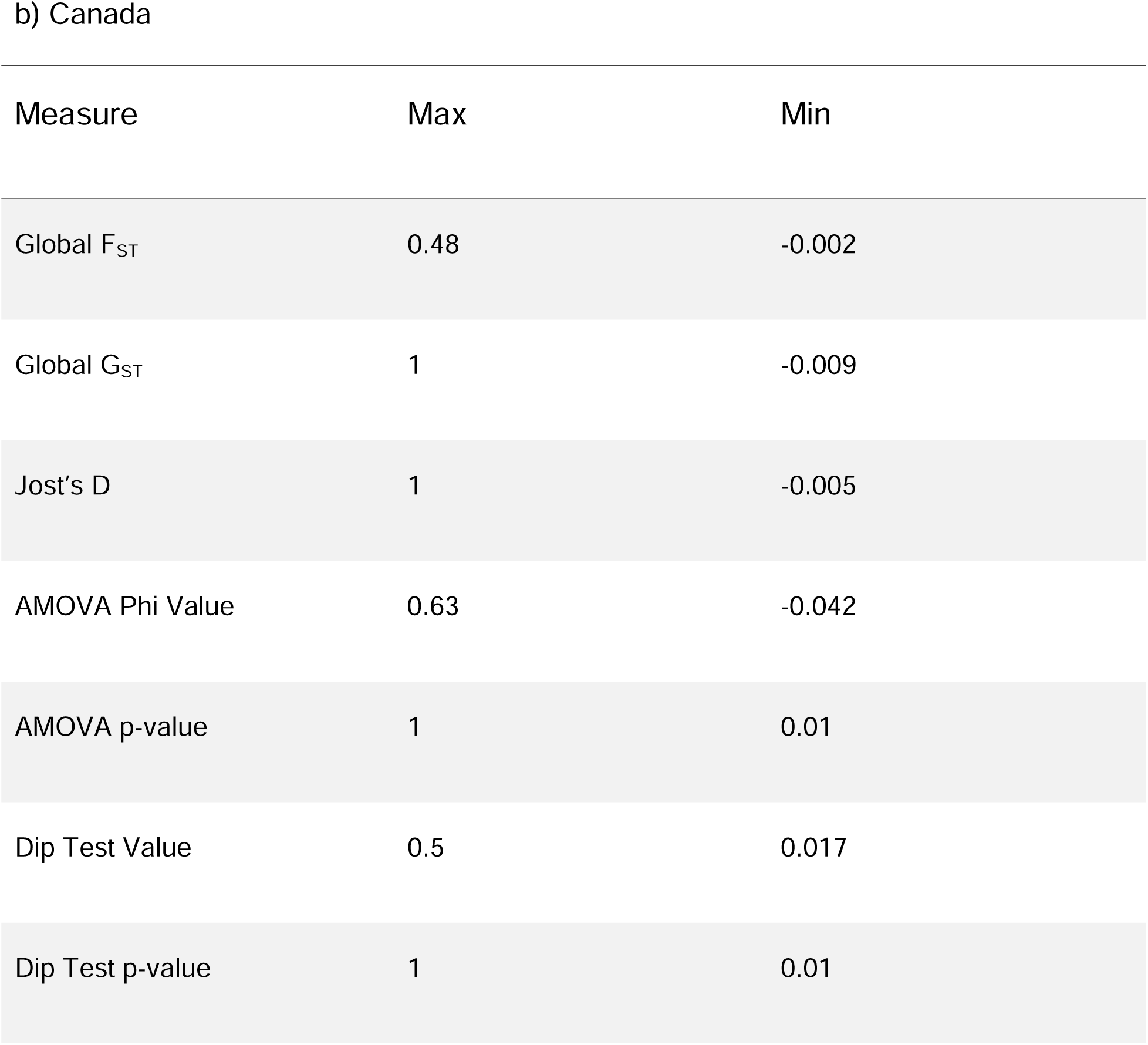
Population genetic structure measures for BINs from a) Greenland and b) Canada.

The Greenland populations showed few signs of genetic differentiation across all measures. There are 716 BINs in the Canada analysis, and only the max and min values for each measure are shown. Most FST values were below 0.1, indicating low levels of genetic differentiation, but most GST and Jost’s D values were above 0.8, indicating higher levels of genetic differentiation. Most AMOVA phi values were below 0.1, revealing low levels of variation between regions. The mismatch analysis revealed no significant signs of multimodality.

The analysis was also completed for Canada. In total, 716 BINs across 37 polygons/regions were included (Fig. 3b; Table 3b). F_ST_, G_ST_, and Jost’s D were determined for each region pair for each BIN (Fig. 5). Due to the dataset size, global F_ST_, G_ST,_ and Jost’s D were also determined for each BIN. Most F_ST_ values were below 0.1, indicating low levels of population genetic differentiation (Fig. 6a). However, most Jost’s D and G_ST_ values were over 0.8, indicating higher levels of genetic differentiation (Fig. 6b & 6c). The AMOVA for most BINs revealed low levels of variation between regions, with most phi values being below 0.1. The mismatch analysis revealed no significant signs of multimodality, suggesting that the populations are still expanding.

**Figure 5:**
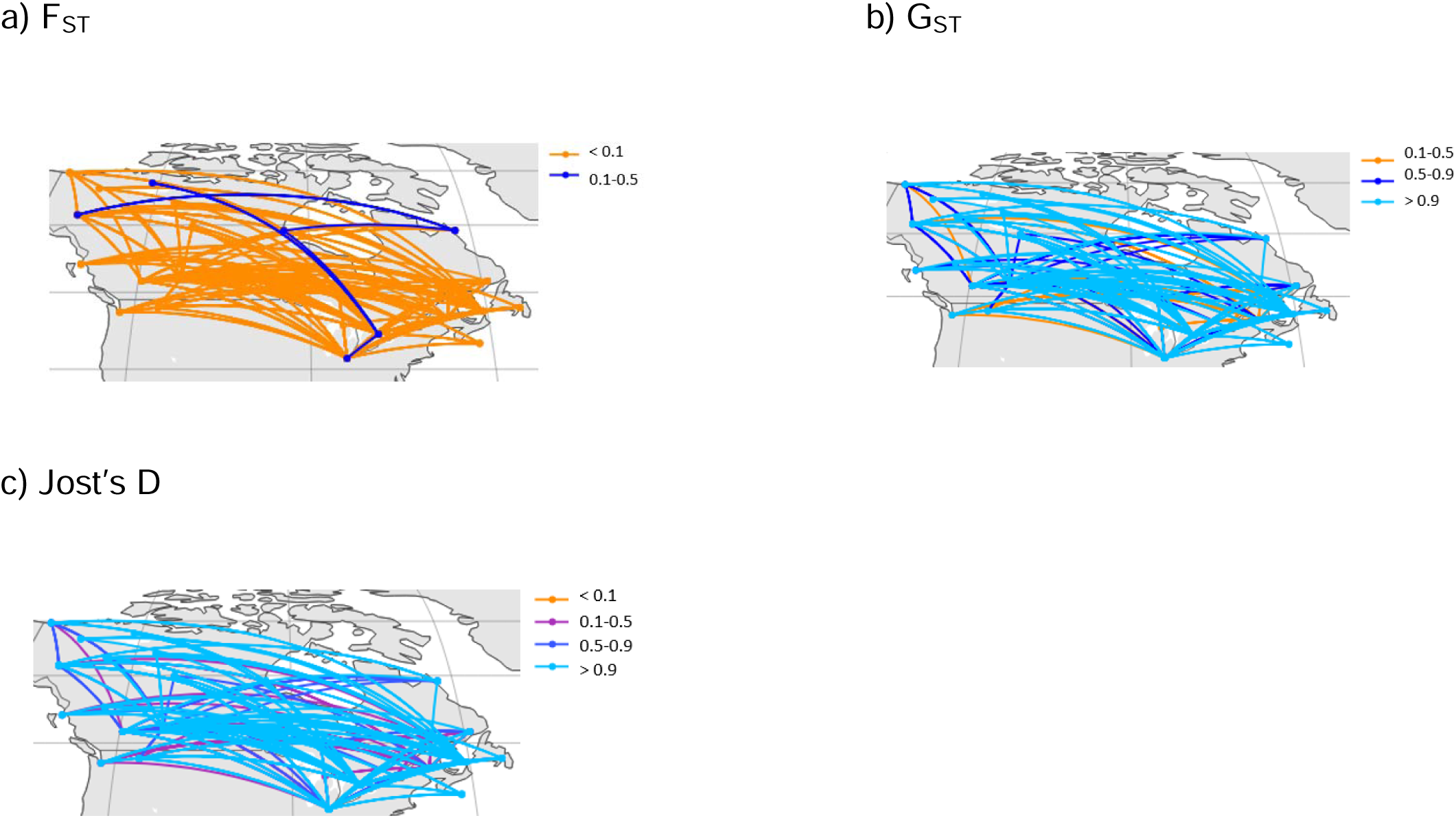
The pairwise a) F_ST_, b) G_ST_, and c) Jost’s D between geographic regions in Canada for the Diptera BIN analysis Only 10% of records from the 10 largest BINs are shown as an example. Most F_ST_ values were below 0.1, suggesting low levels of genetic differenitation, while most G_ST_ and Jost’s D values were above 0.9, suggesting higher levels of genetic differentiation.

**Figure 6:**
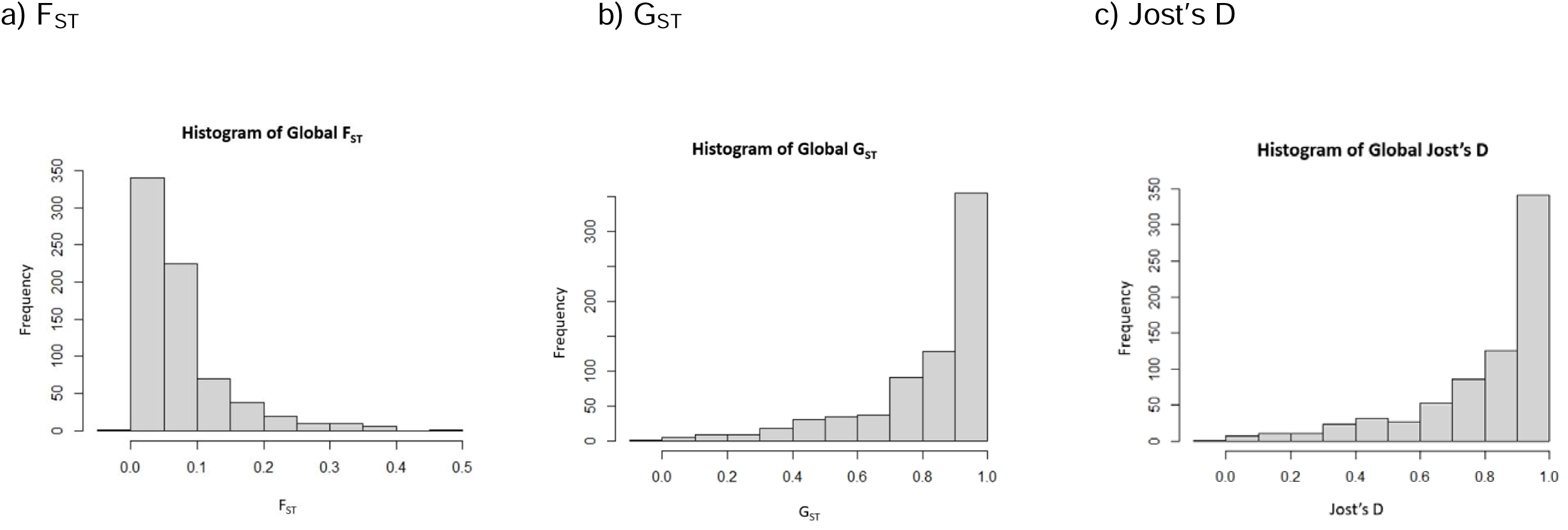
The distribution of the global (one value per BIN) a) Nei F_ST_ values, b) Hedrick’s G_ST_ values, and c) Jost’s D values for the Canada Diptera BIN analysis (716 BINs). Most F_ST_ values were below 0.1, suggesting low levels of genetic differenitation, while most G_ST_ and Jost’s D values were above 0.8, suggesting higher levels of genetic differentiation.

The measures for each individual species are included as a csv file in S2 Appendix.

#### 3.2.2 Comparison Across Traits

The population genetic structure measures were compared across species/BINs with different biological traits. The number of species from the Greenland BIN analysis with different biological trait categories is shown in Table 4a. Traits represented by only one BIN were dropped from the statistical analyses. For both the ANOVA and PGLS analyses, there was no significant relationship between the measures and biological traits in the Greenland dataset (Fig. 7, 8 & 9). As ANOVAs assume a normal distribution, a Shapiro-Wilk test was performed to test whether this assumption was met. This assumption was violated (p-value = 0.0085), so a Kruskal-Wallis test was performed. The results did not differ between the Kruskal-Wallis and ANOVA. ANOVAs also assume homogeneity of variances, so a Levene test was performed. This assumption was not violated (p-value = 0.68). The range size analysis was also done for Greenland, and there was no significant relationship between range size, the population genetic measures, and traits.

**Figure 7:**
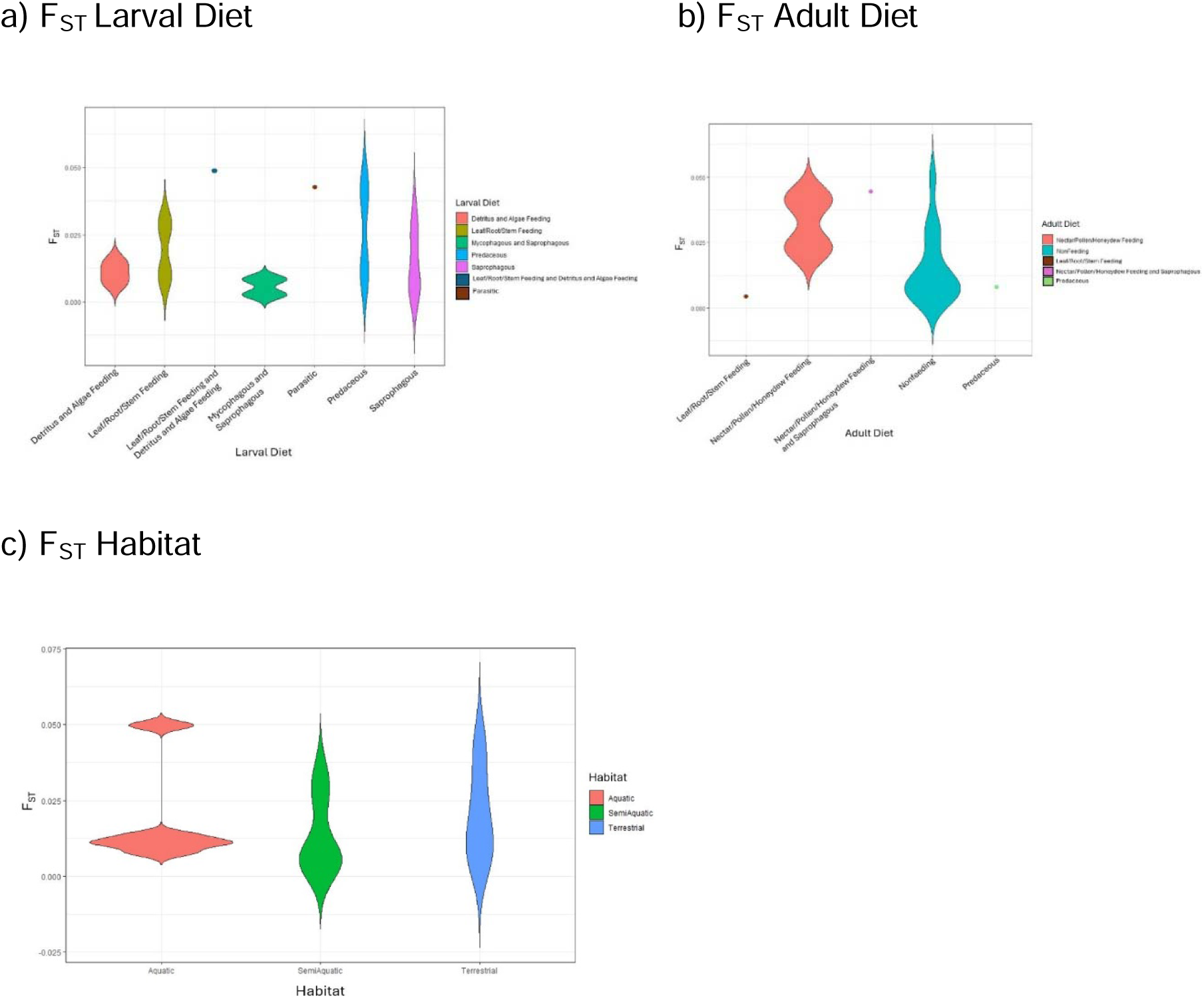
Violin plots showing the relationship between biological traits and population genetic structure measures for the BIN analysis for Greenland Diptera (20 BINs). Results are shown comparing Nei’s F_ST_ to a) larval diet, b) adult diet, and c) habitat. There was no significant relationship between the population genetic structure measures and traits. Traits with fewer than two datapoints were dropped from the analysis, but are shown by dots on the figures

**Figure 8:**
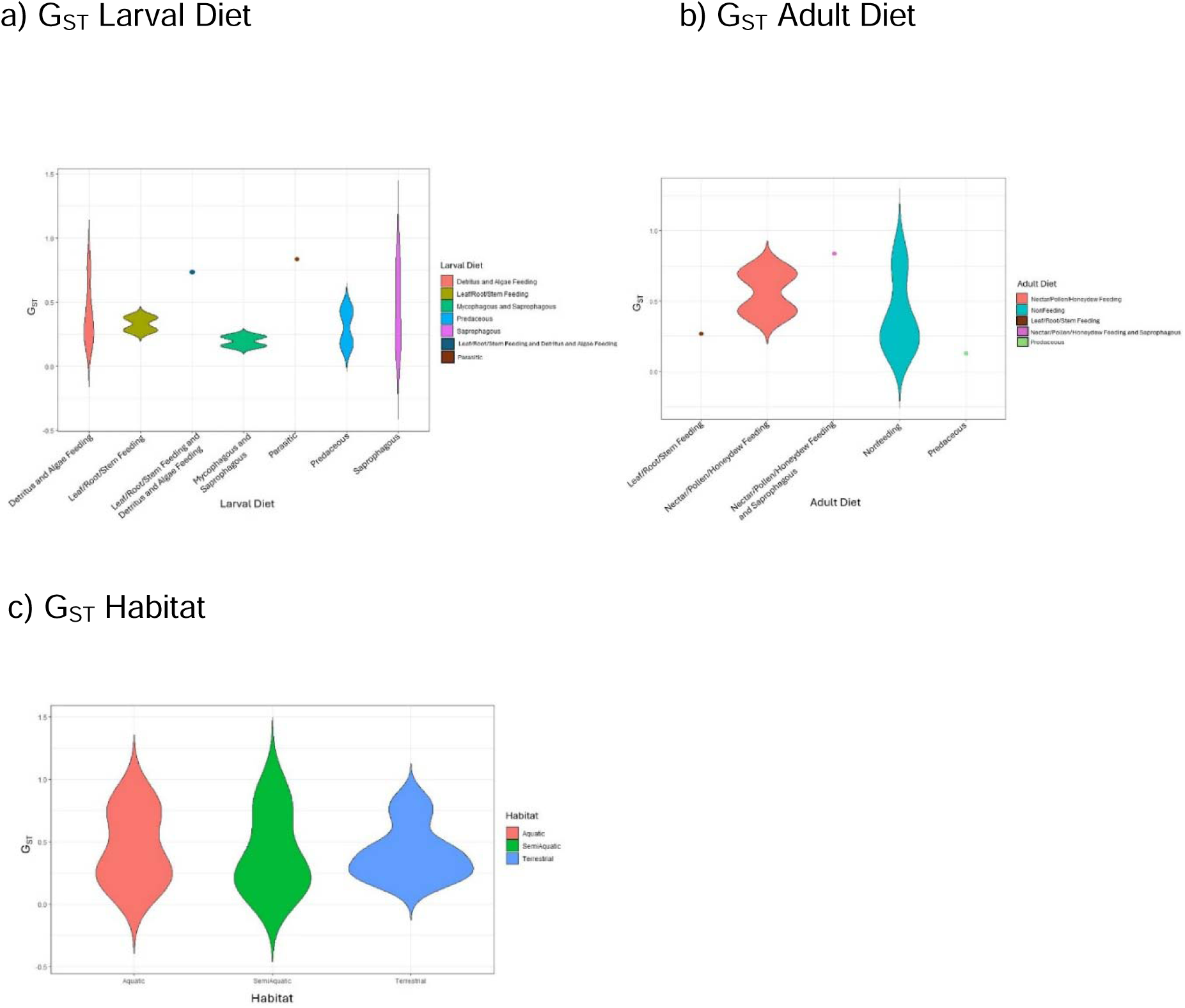
Violin plots showing the relationship between biological traits and population genetic structure measures for the BIN analysis for Greenland Diptera (20 BINs). Results are shown comparing Hedricks G_ST_ to a) larval diet, b) adult diet, and c) habitat. There was no significant relationship between the population genetic structure measures and traits. Traits with fewer than two datapoints were dropped from the analysis, but are shown by dots on the figures.

**Figure 9:**
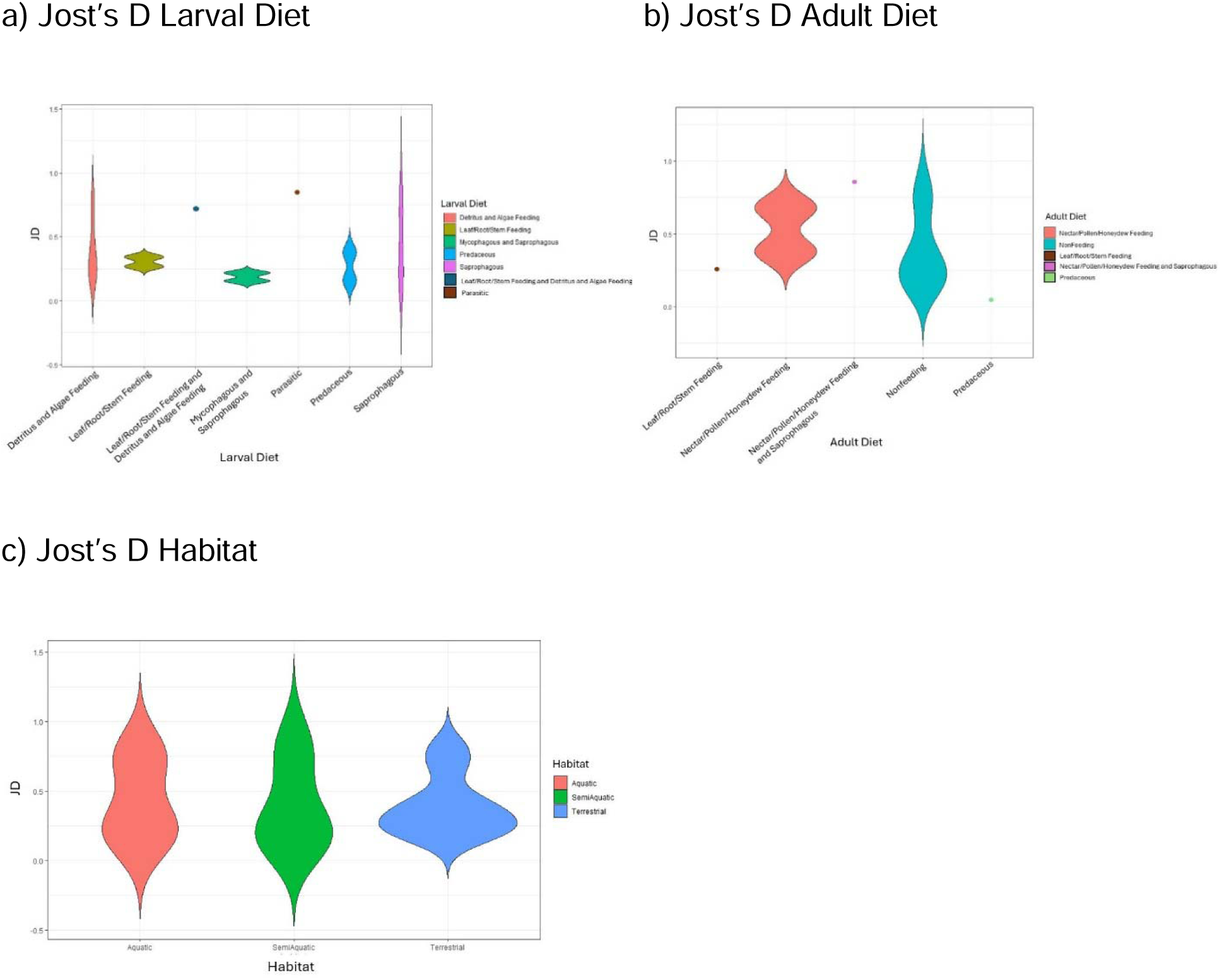
Violin plots showing the relationship between biological traits and population genetic structure measures for the BIN analysis for Greenland Diptera (20 BINs). Results are shown comparing Jost’s D to a) larval diet, b) adult diet, and c) habitat. There was no significant relationship between the population genetic structure measures and traits. Traits with fewer than two datapoints were dropped from the analysis, but are shown by dots on the figures.

**Table 4:**
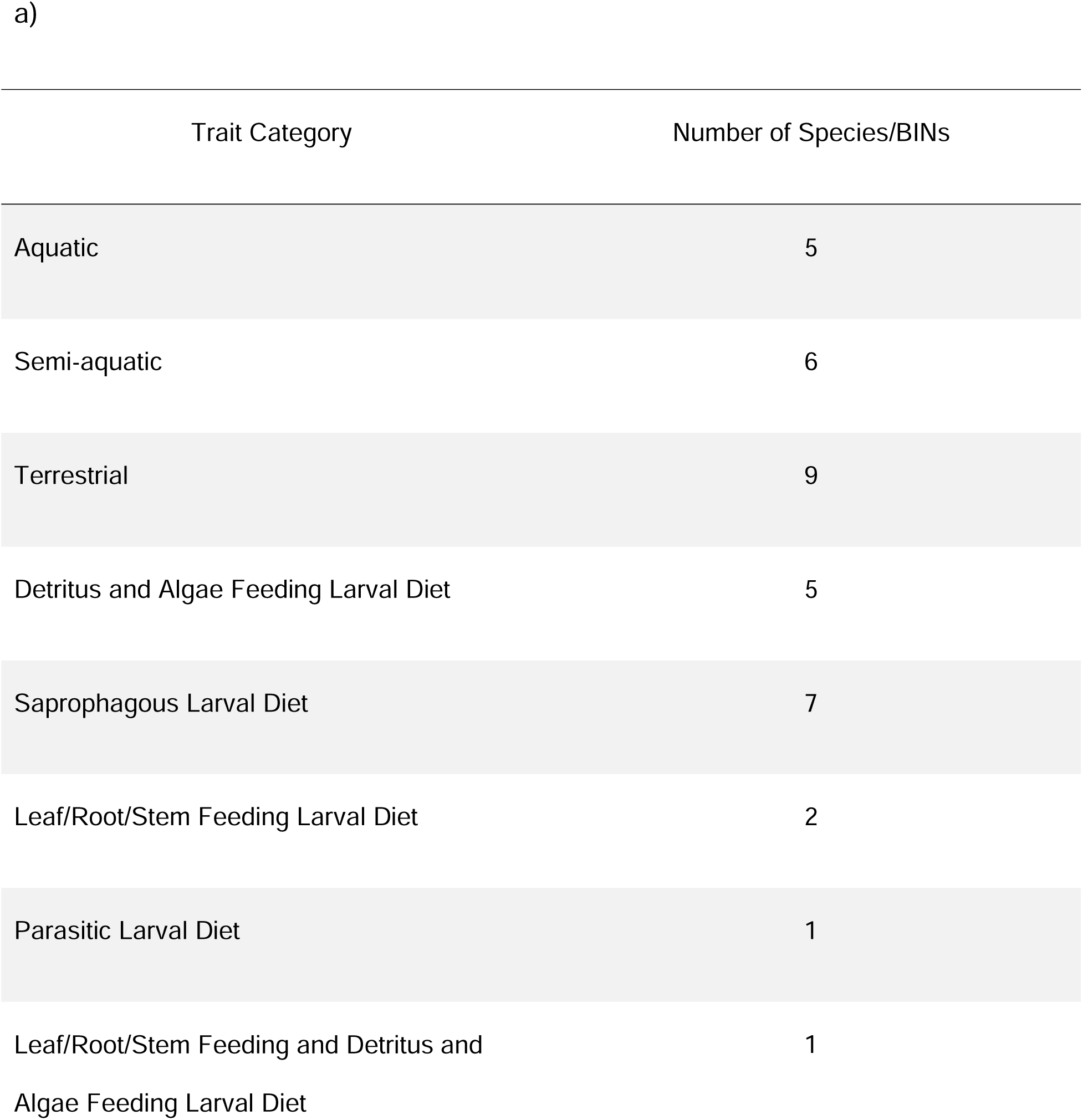
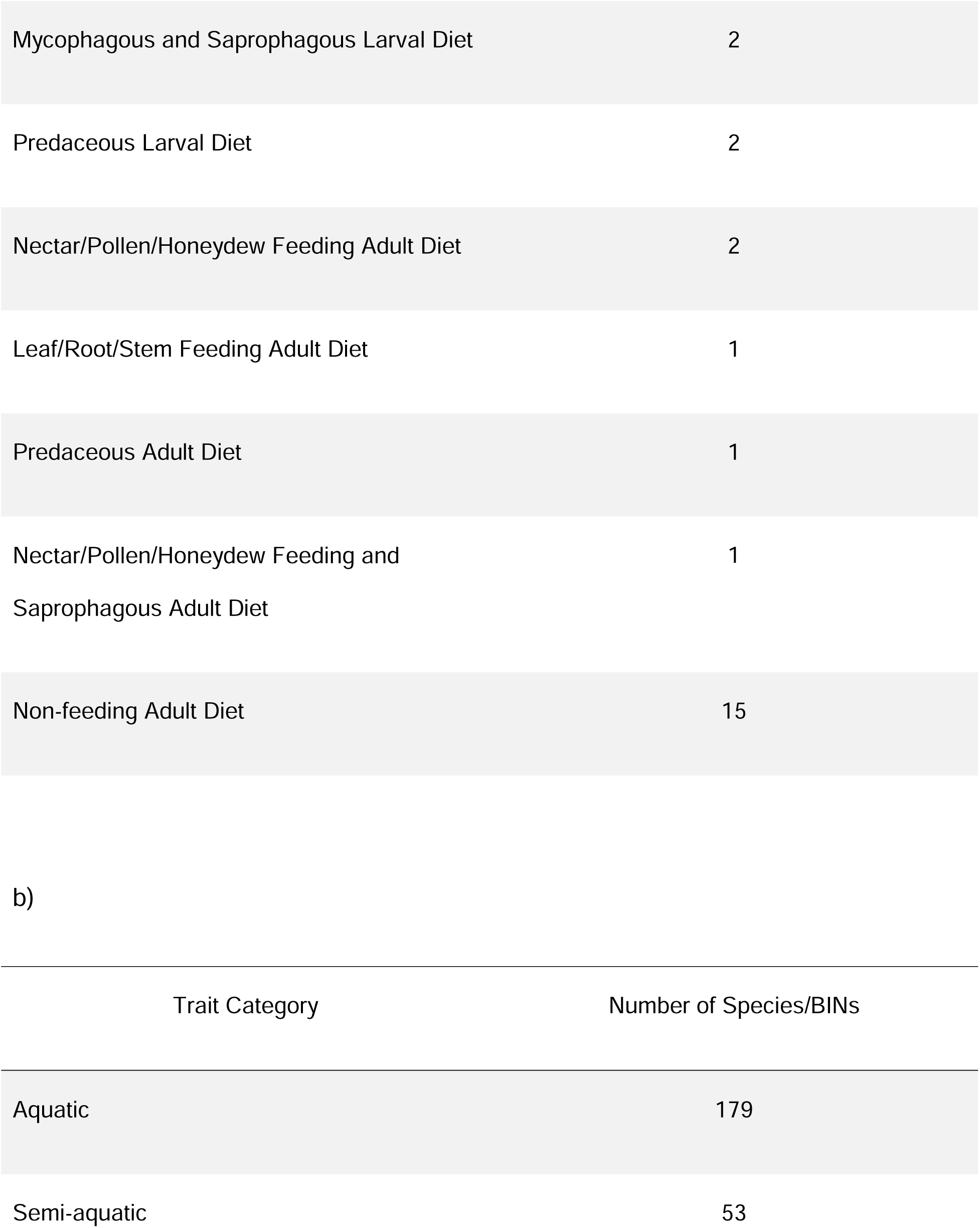
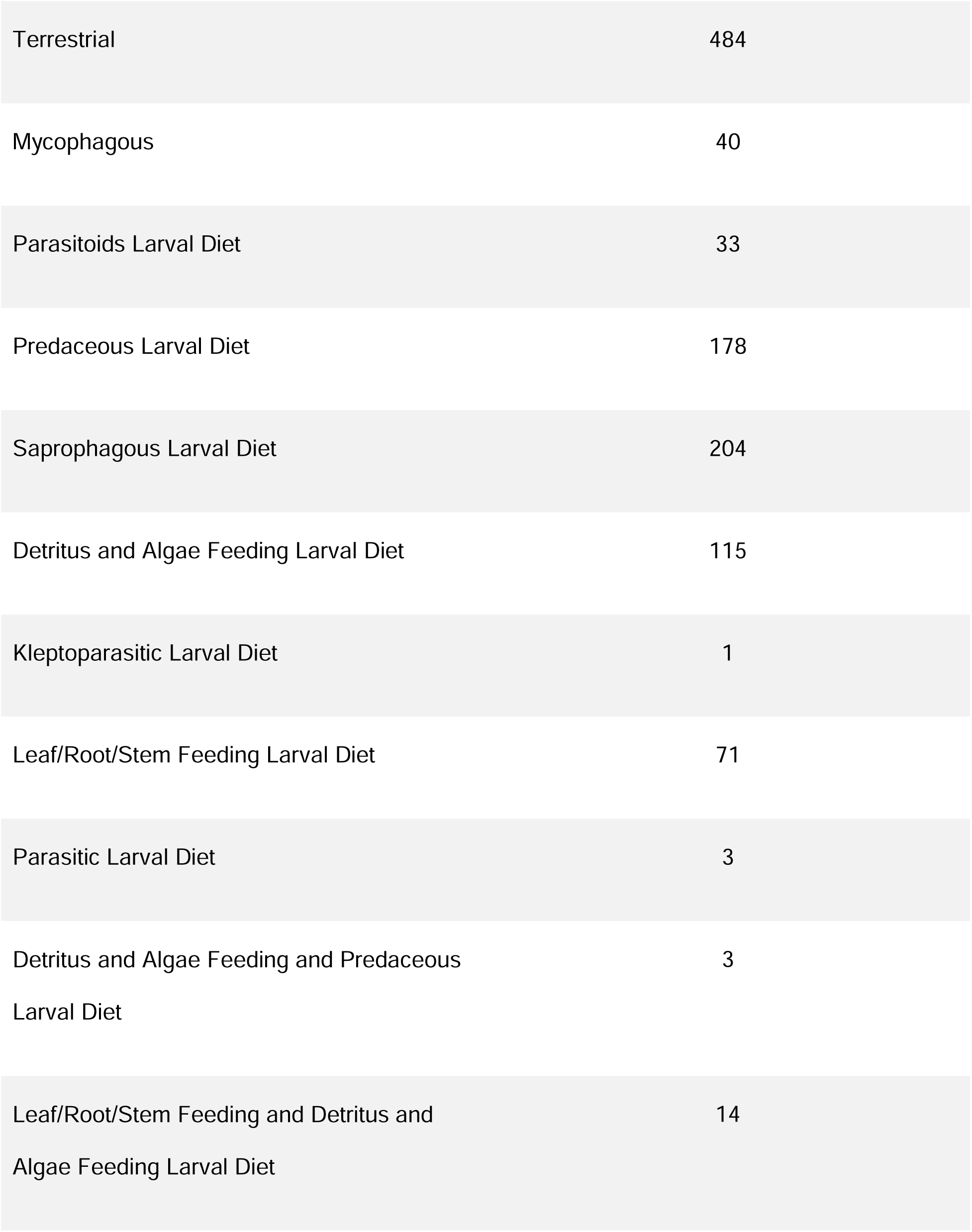
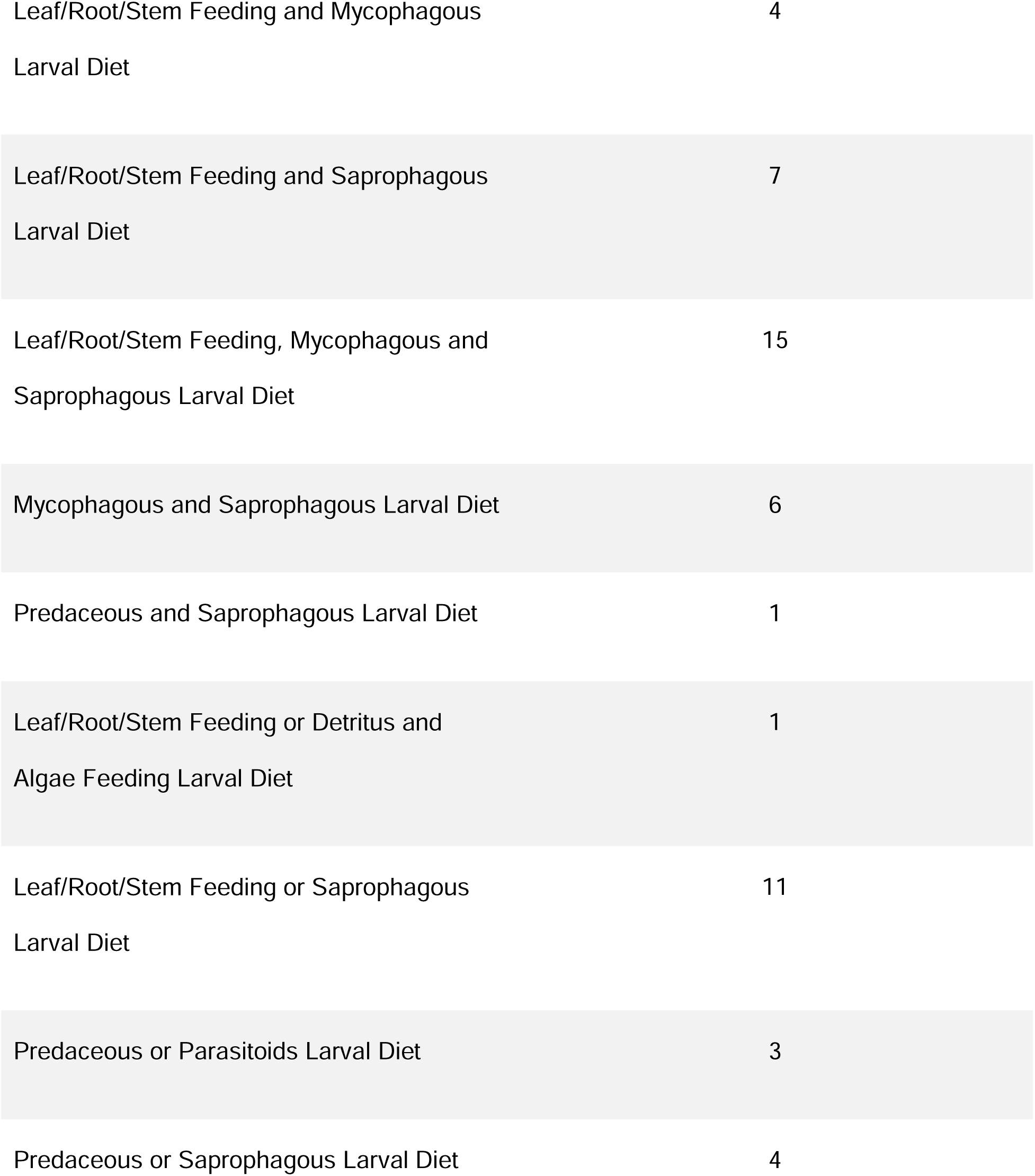
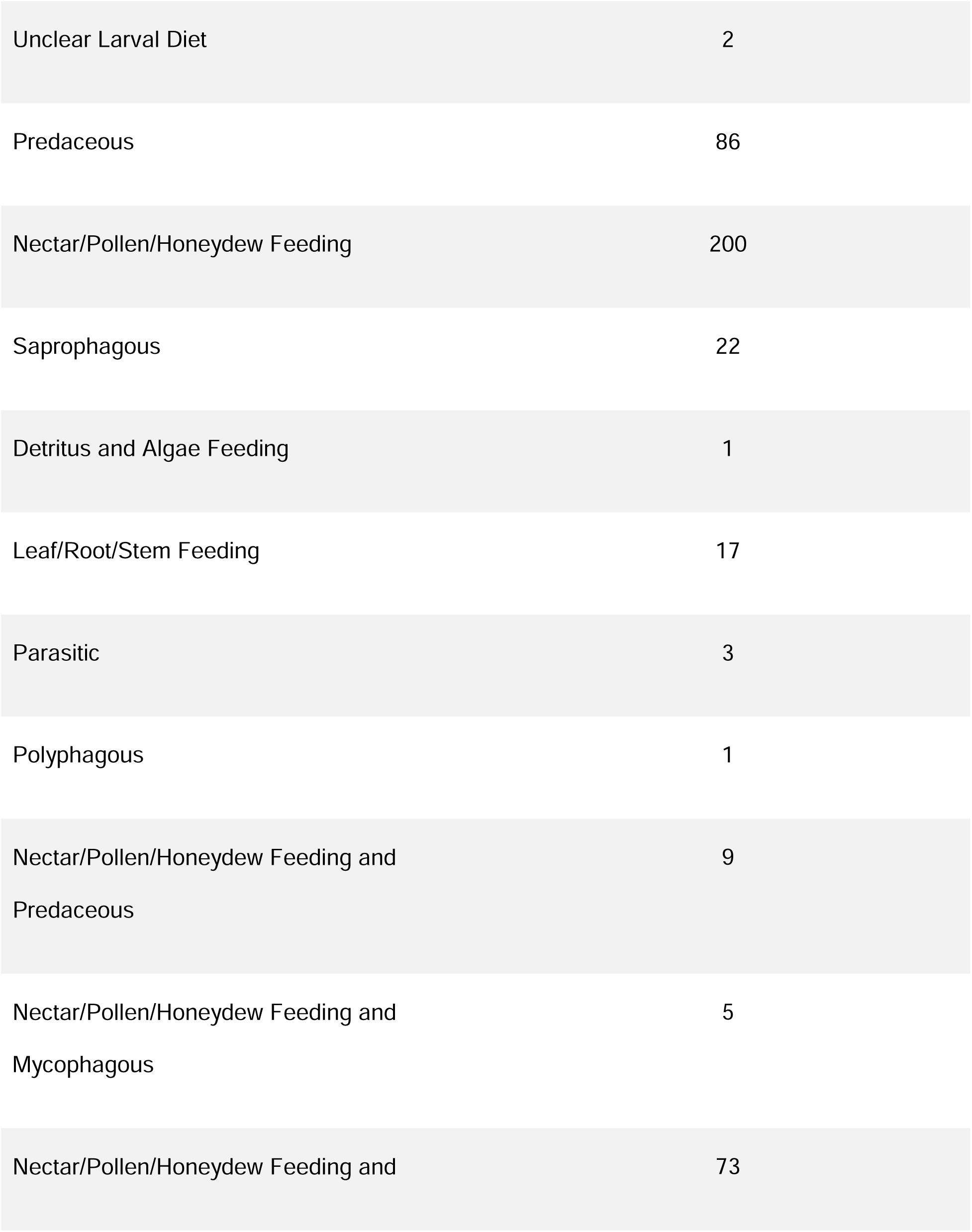
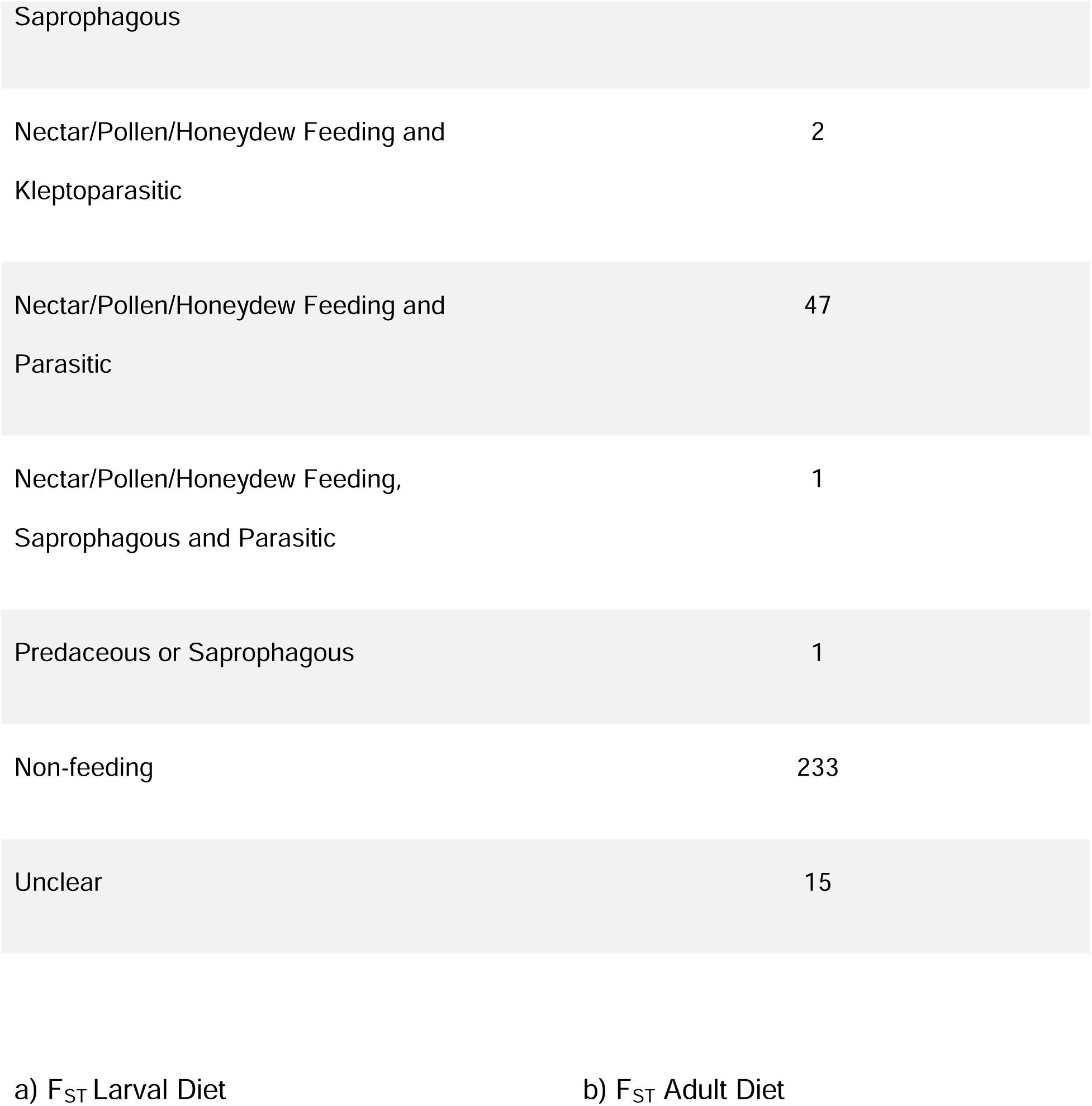
The number of species with each biological trait category for the a) Greenland BIN analysis and the b) Canada BIN analysis.

The number of species from the Canada BIN analysis with different biological trait categories is shown in Table 4b. The multiple regression revealed a significant relationship between the population genetic structure measures and habitat only in F_ST_ (Fig. 10; Table 5). Semi-aquatic populations were associated with significantly lower levels of genetic differentiation. Larval diet was also significantly related to population genetic structure measures (Fig. 11). Having a larval stage that is both leaf/root/stem feeding and mycophagous, or mycophagous and saprophagous, resulted in significantly lower levels of genetic differentiation for Jost’s D and G_ST_. The F_ST_ results suggested that higher levels of genetic differentiation were observed in BINs with a kleptoparasitic larval diet, a leaf/root/stem feeding diet, a diet that was a mix of leaf/root/stem feeding and saprophagous, and a diet that was either leaf/root/stem feeding or saprophagous.

**Figure 10:**
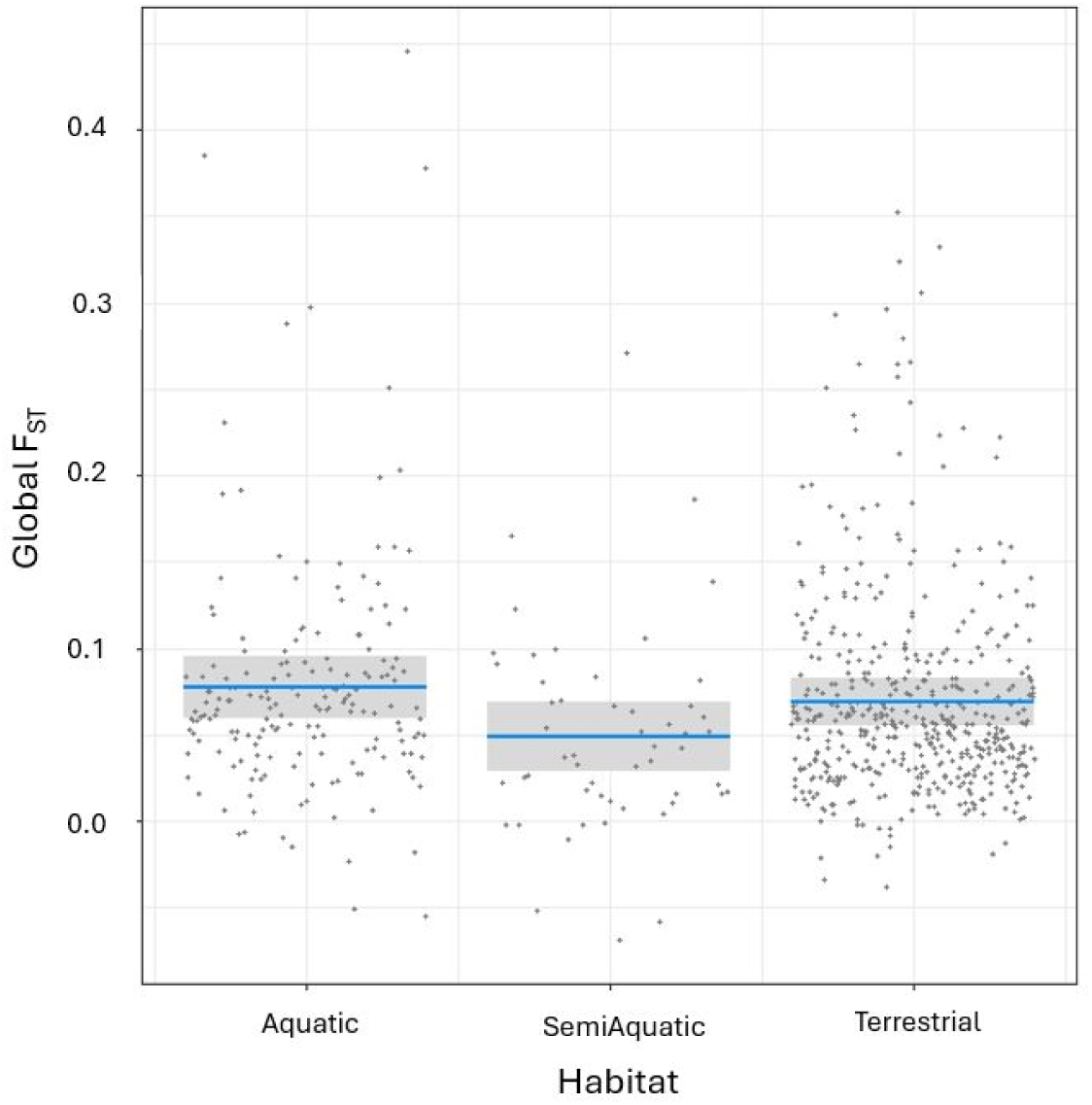
Plots showing the relationships between the global population genetic structure measures and habitat for the Canadian dataset of 716 Diptera BINs. Each dot represents a BIN. The relationship between habitat and F_ST_ is shown, as this is the only significant relationship with habitat (Table 3). F_ST_ suggests that semi-aquatic populations have significantly lower levels of population genetic differentiation.

**Figure 11:**
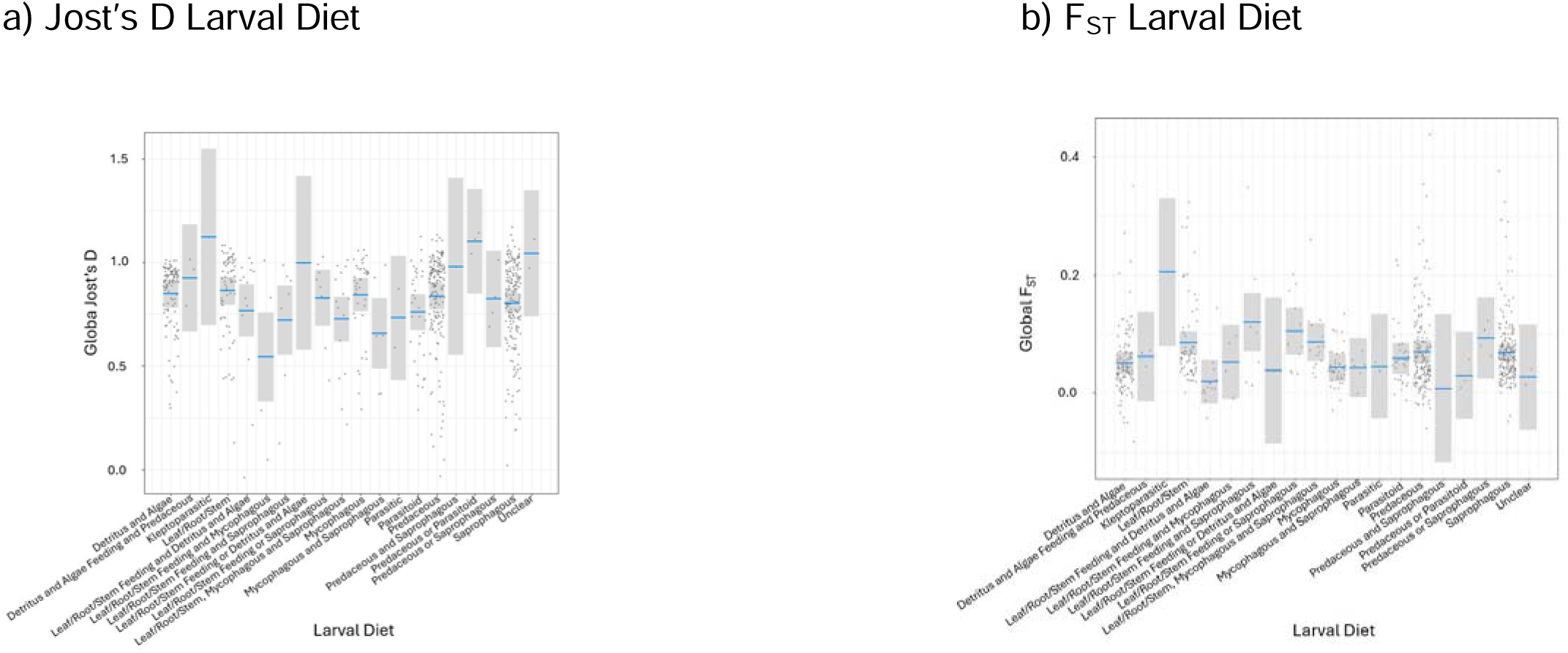
Plots showing the relationships between the global population genetic structure measures and larval diet for the Canadian dataset of 716 Diptera BINs. Each dot represents a BIN. Only the significant relationships from the multiple regression are shown (Table 3). The relationship between larval diet and i) Jost’s D and ii) F_ST_ are shown. Jost’s D and G_ST_ suggested significantly lower levels of genetic differentiation in BINs with a larval stage that is both leaf/root/stem feeding and mycophagous, or mycophagous and saprophagous. F_ST_ suggested significantly higher levels of genetic differentiation in BINs with a kleptoparasitic larval diet, a leaf/root/stem feeding diet, a diet that was a mix of leaf/root/stem feeding and saprophagous, and a diet that was either leaf/root/stem feeding or saprophagous.

**Table 5:**
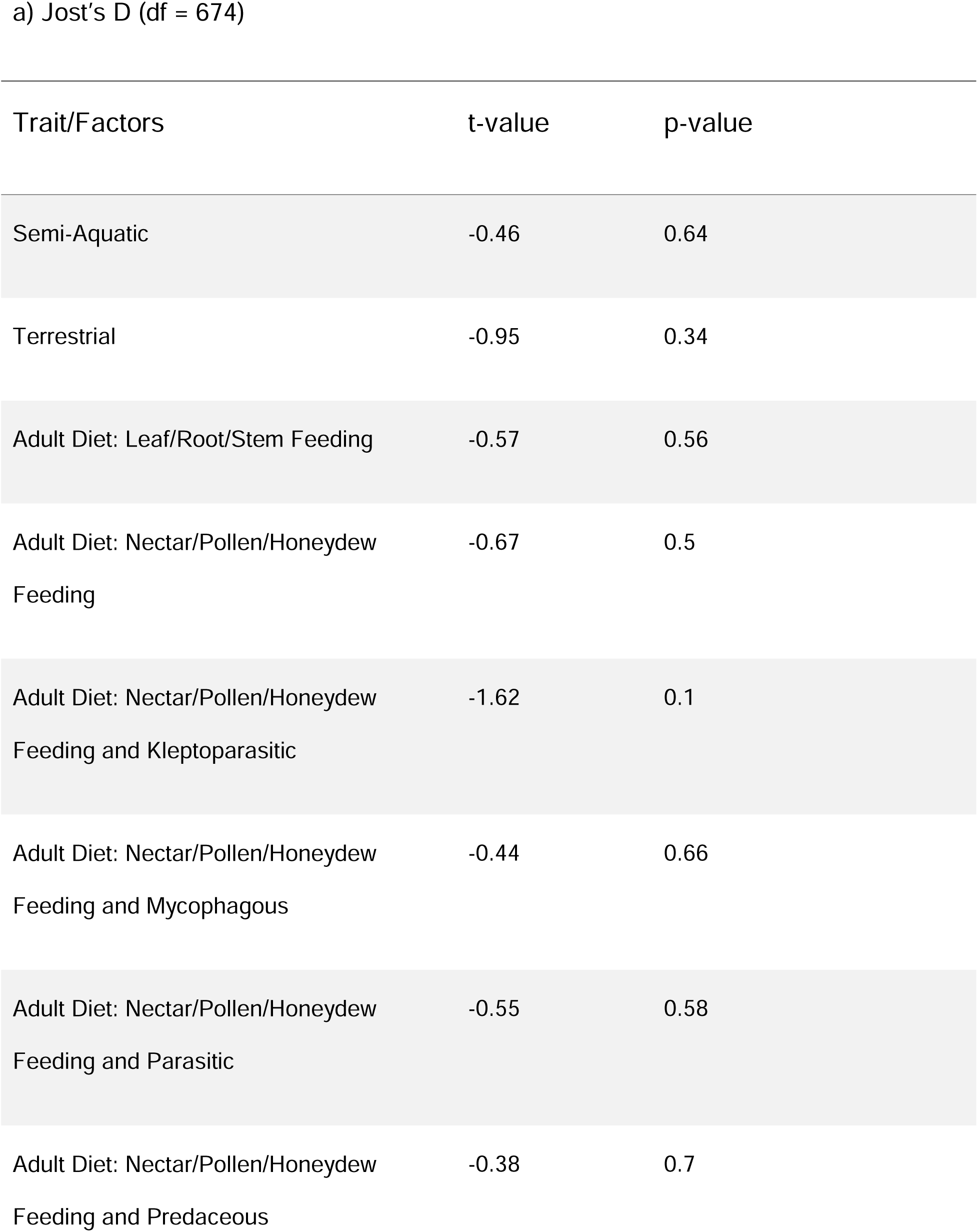
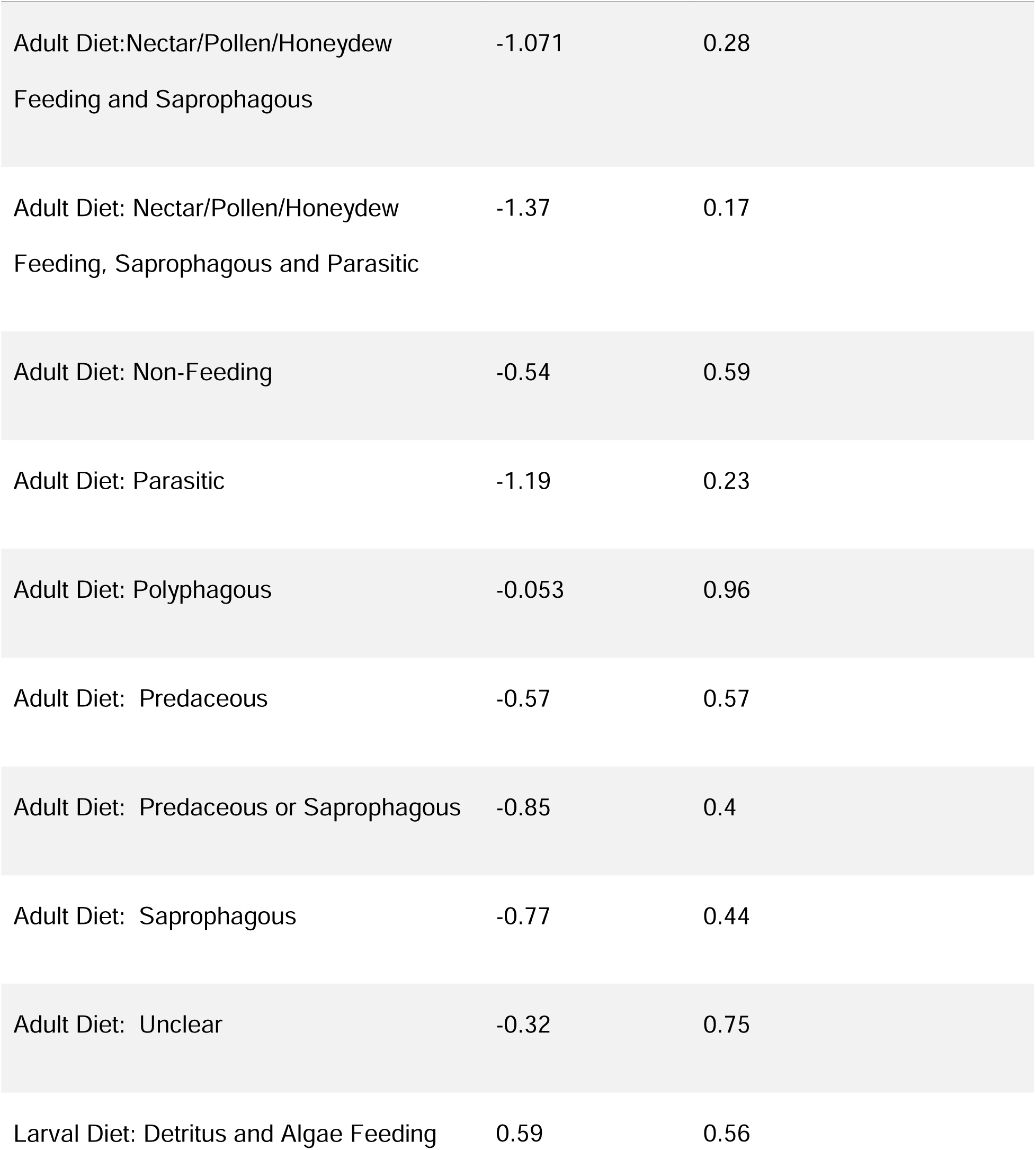
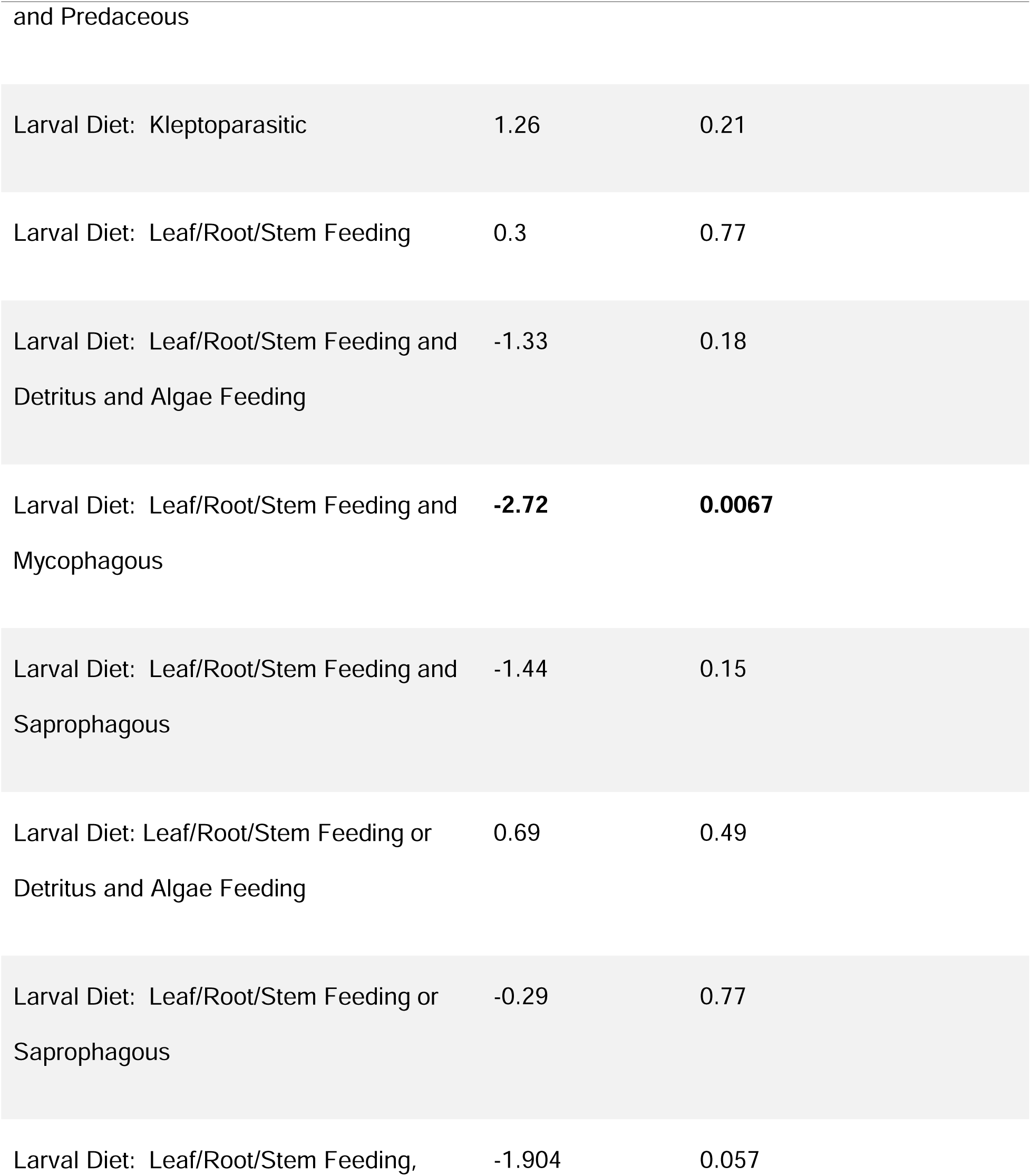
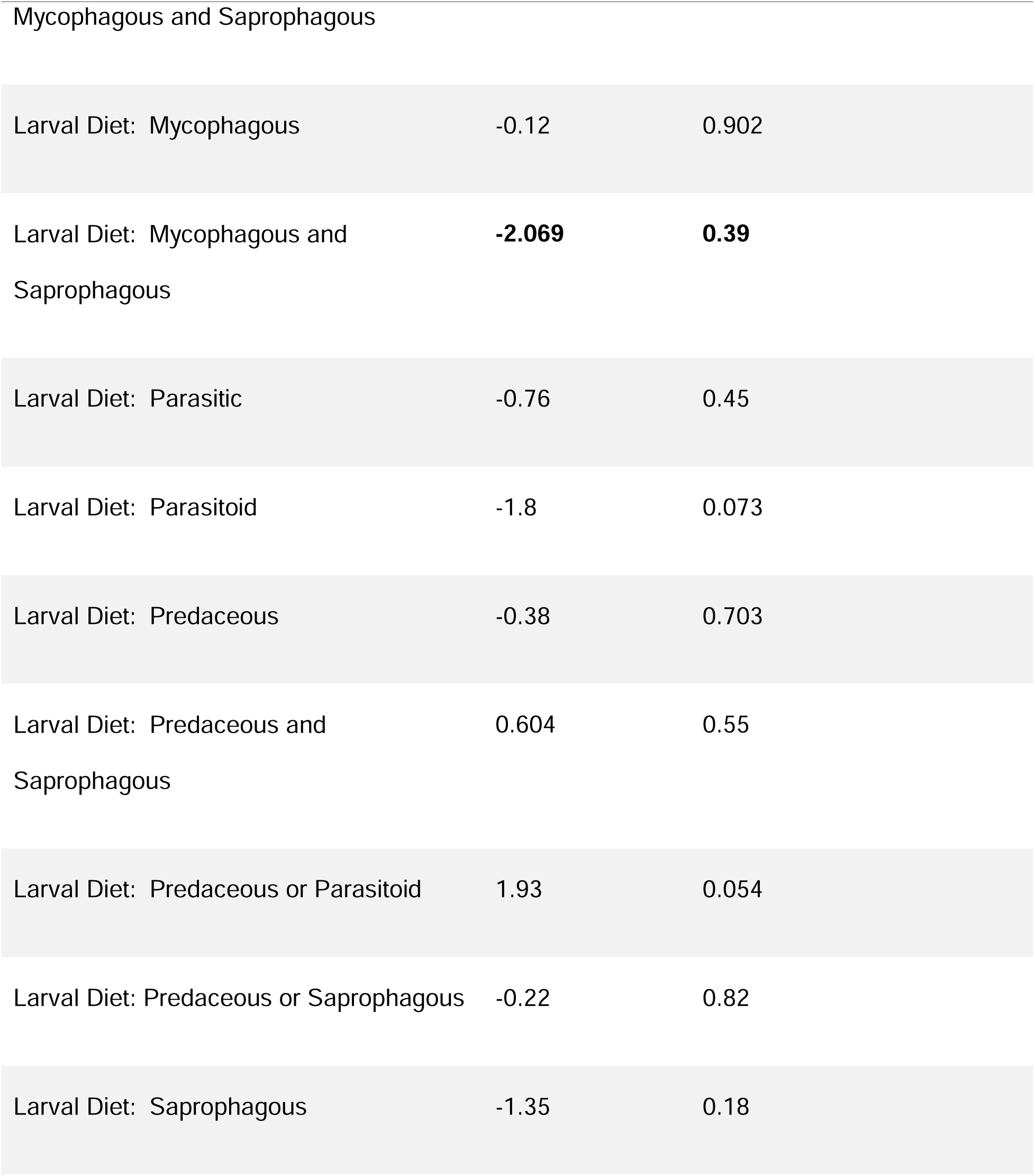
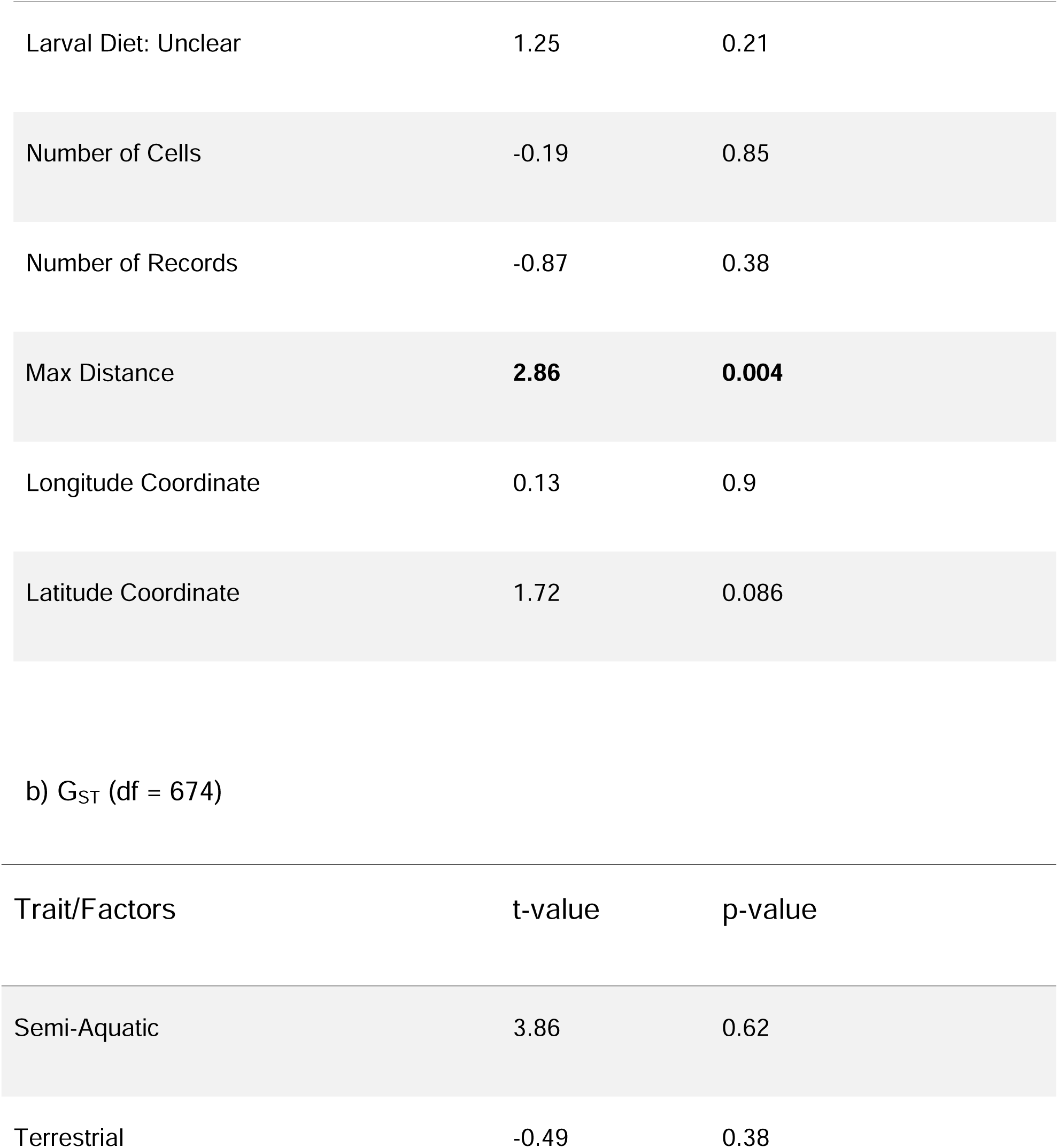
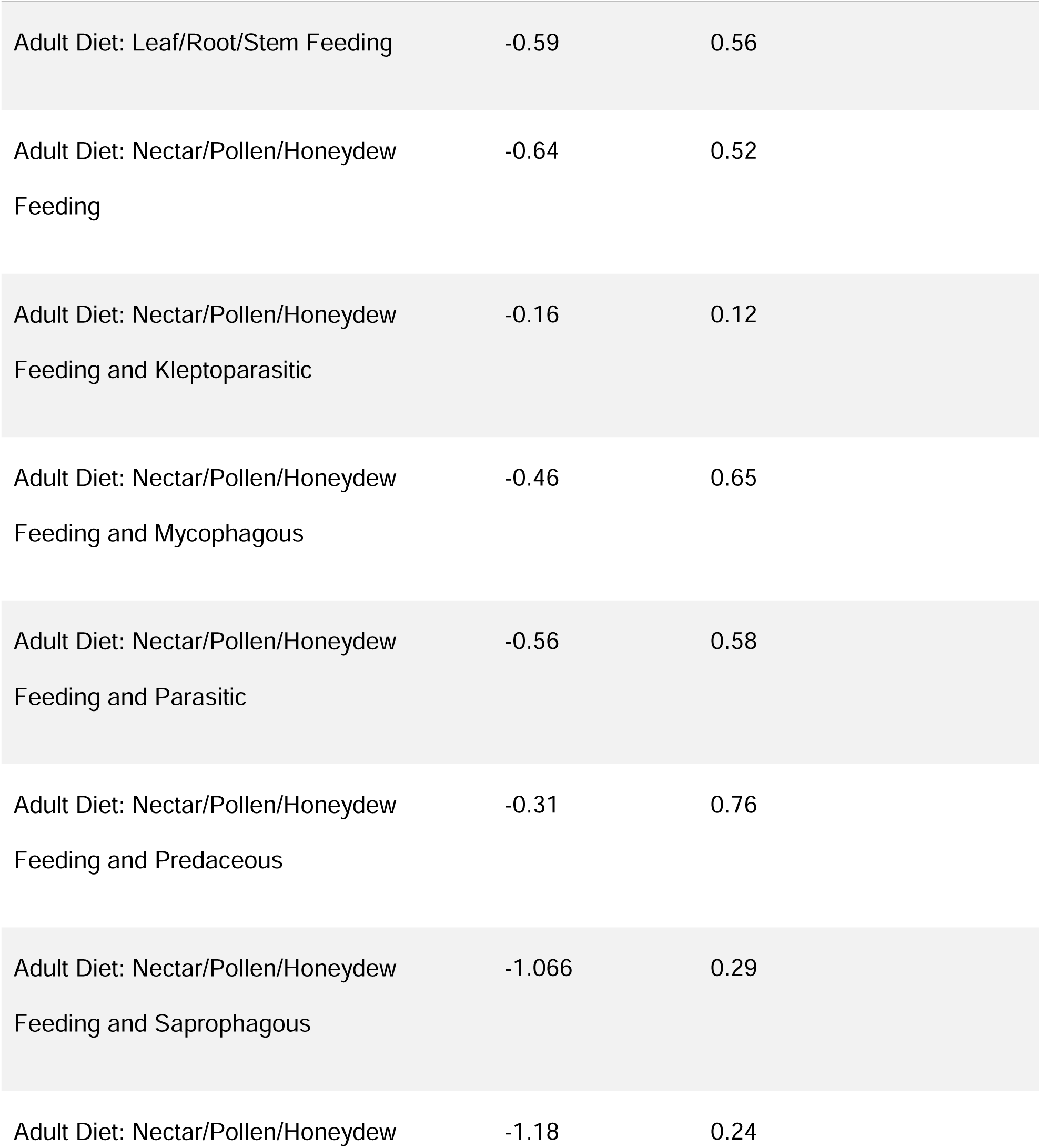
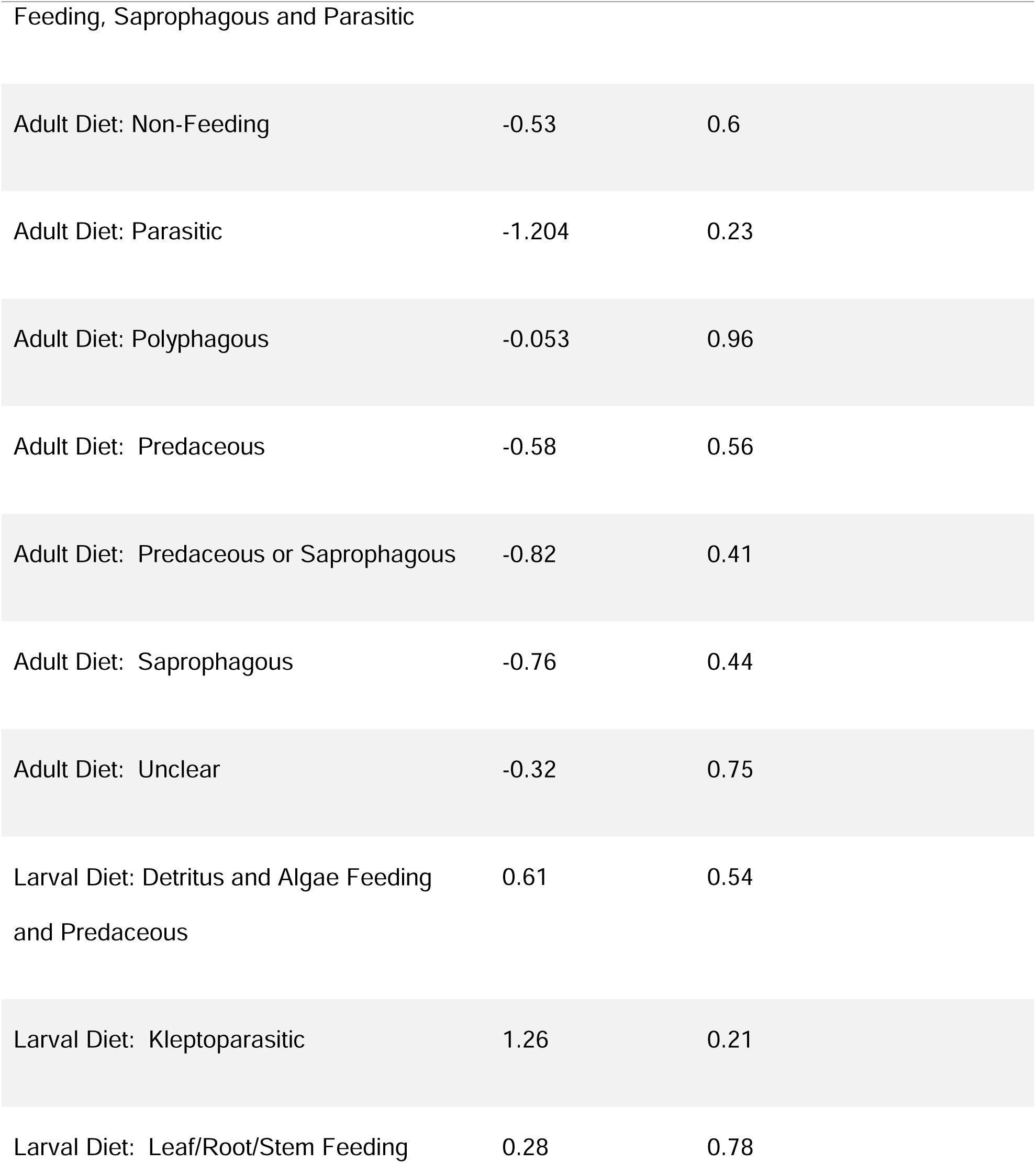
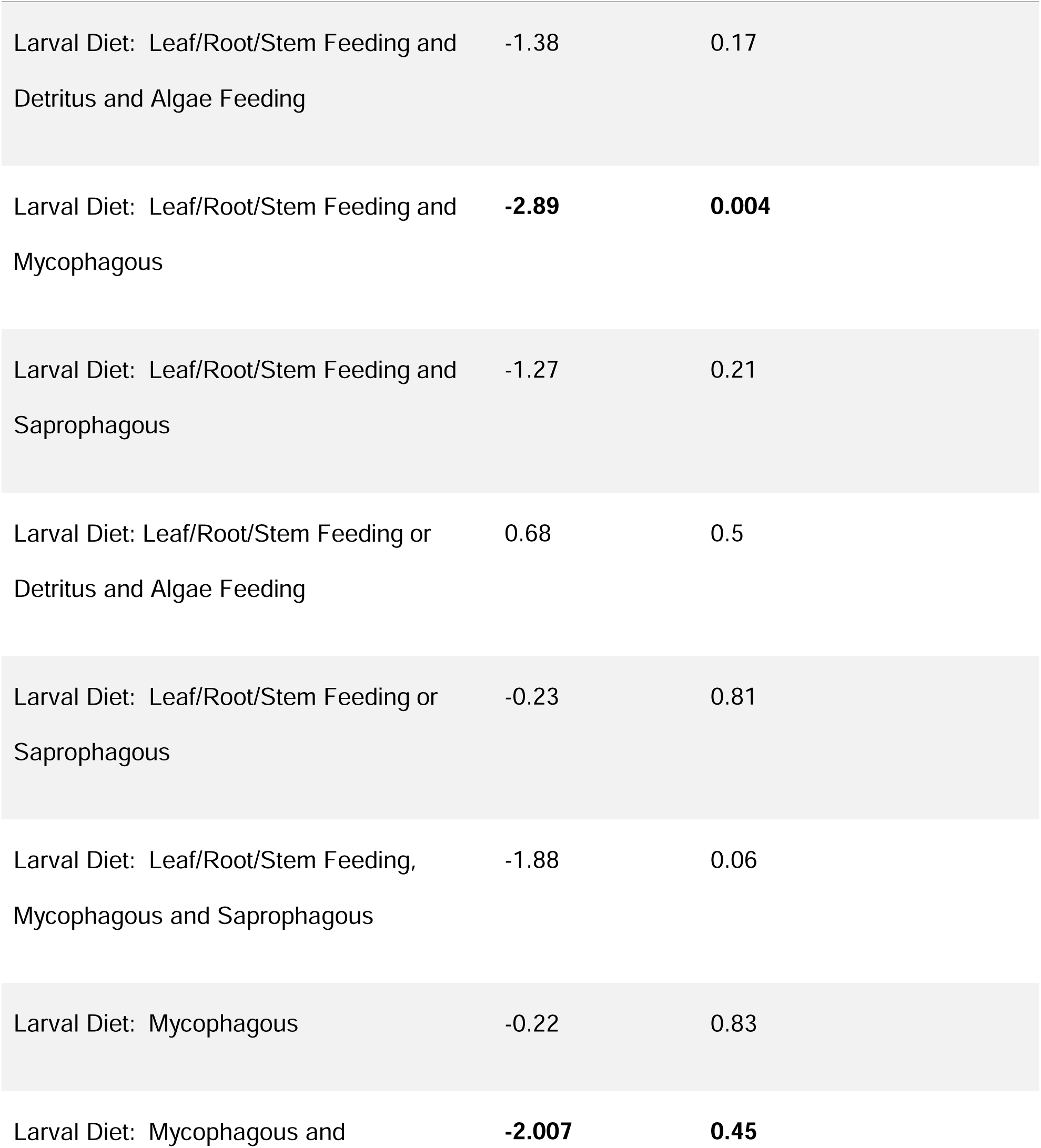
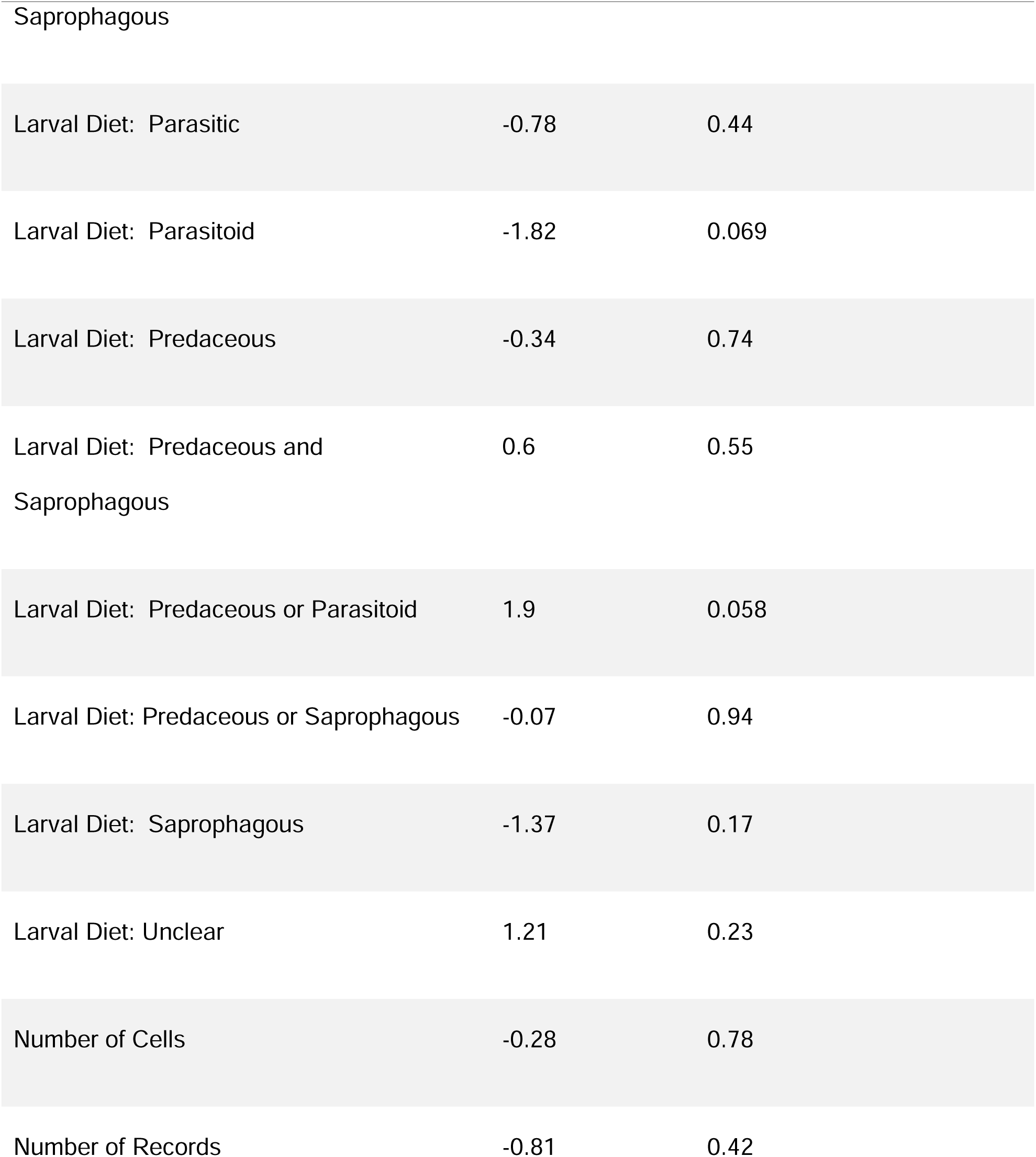
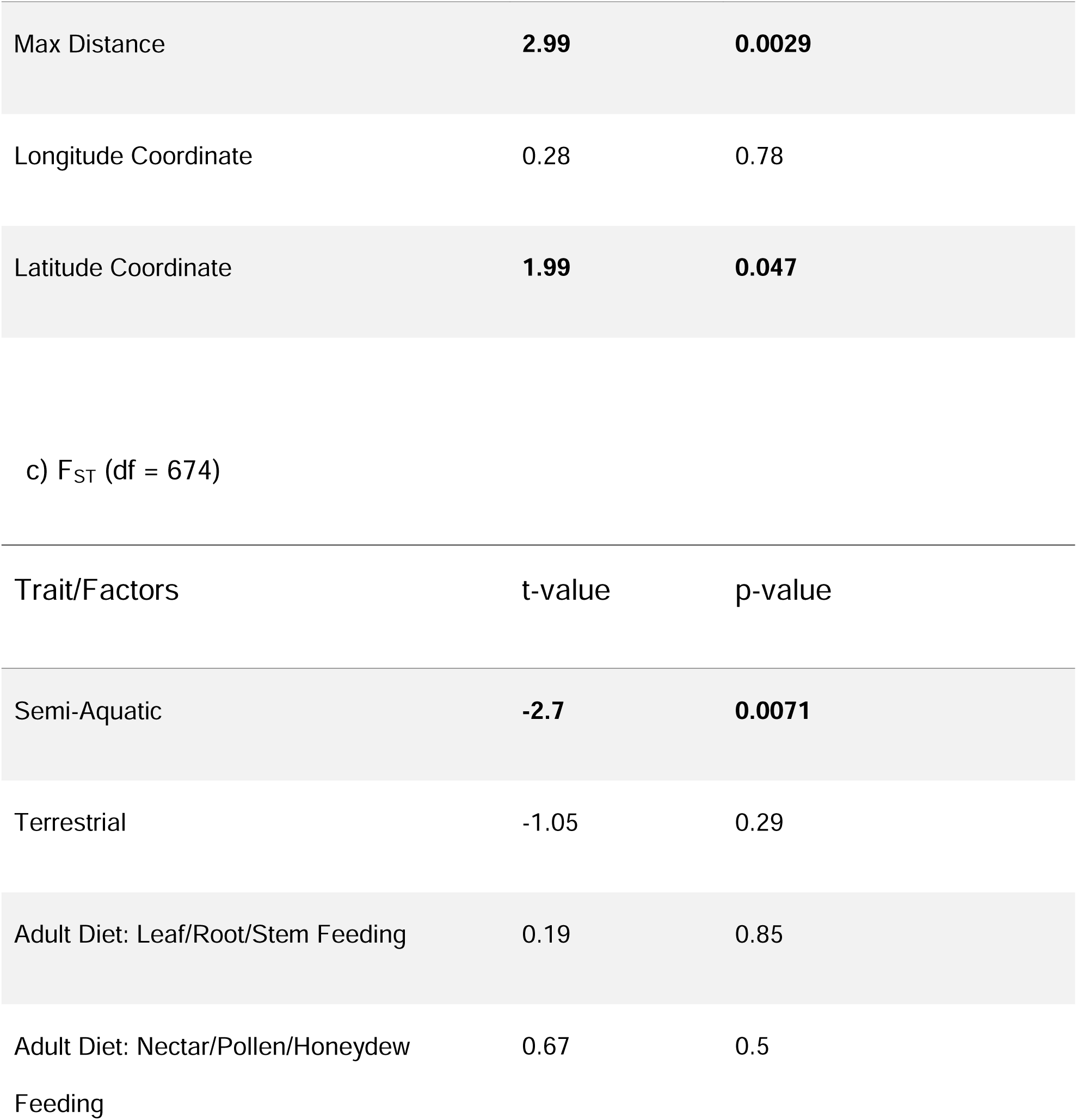
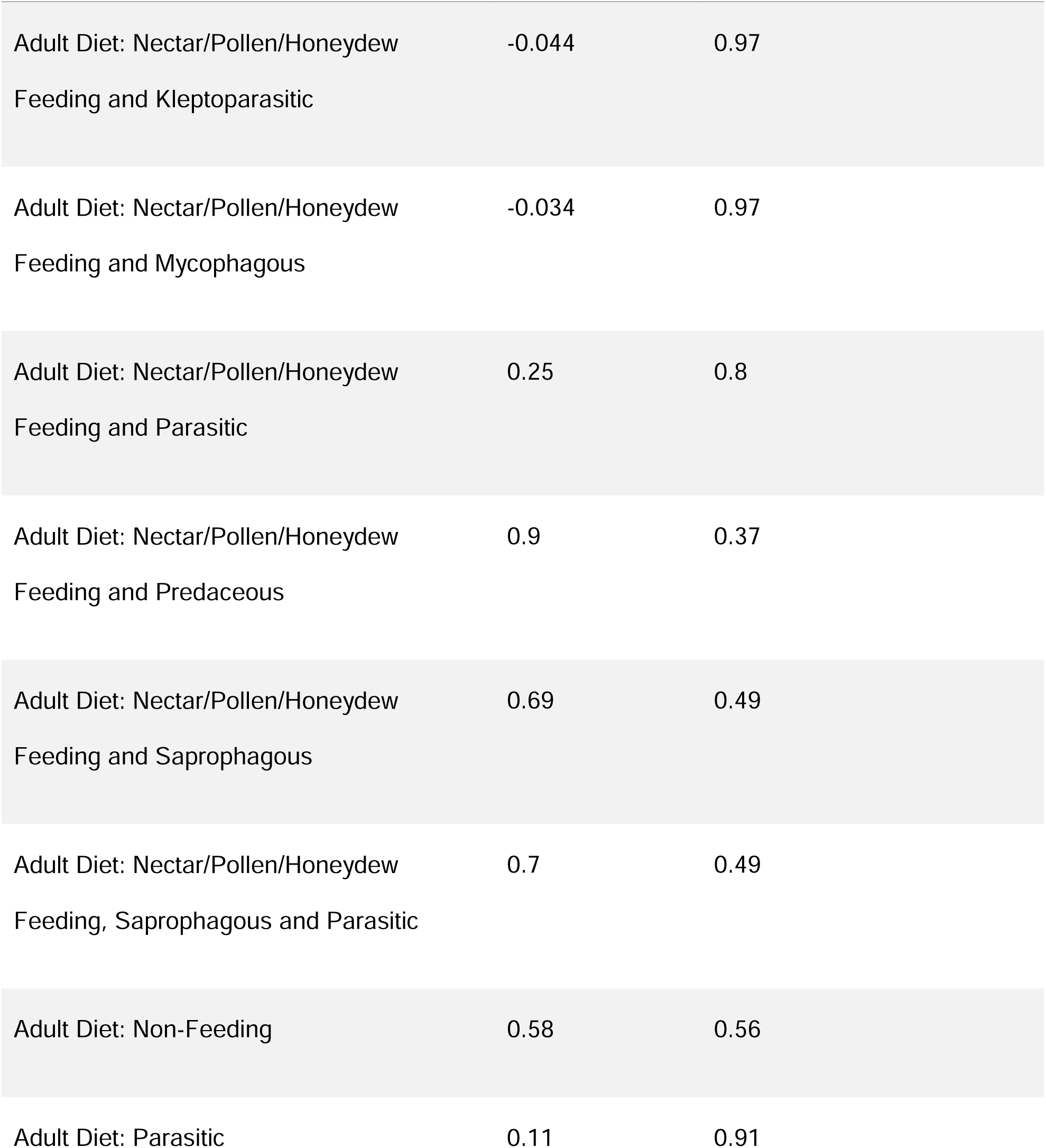
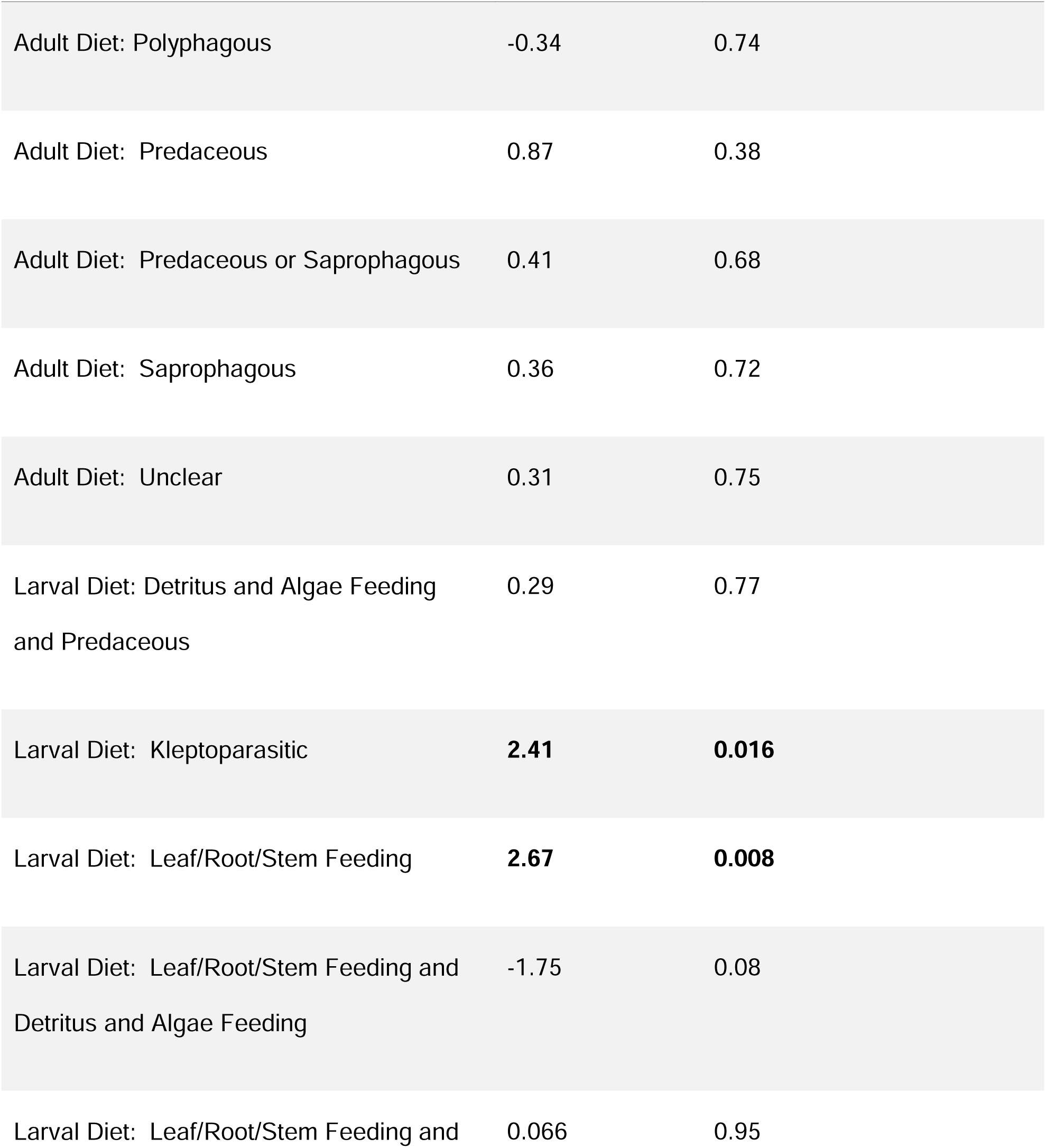
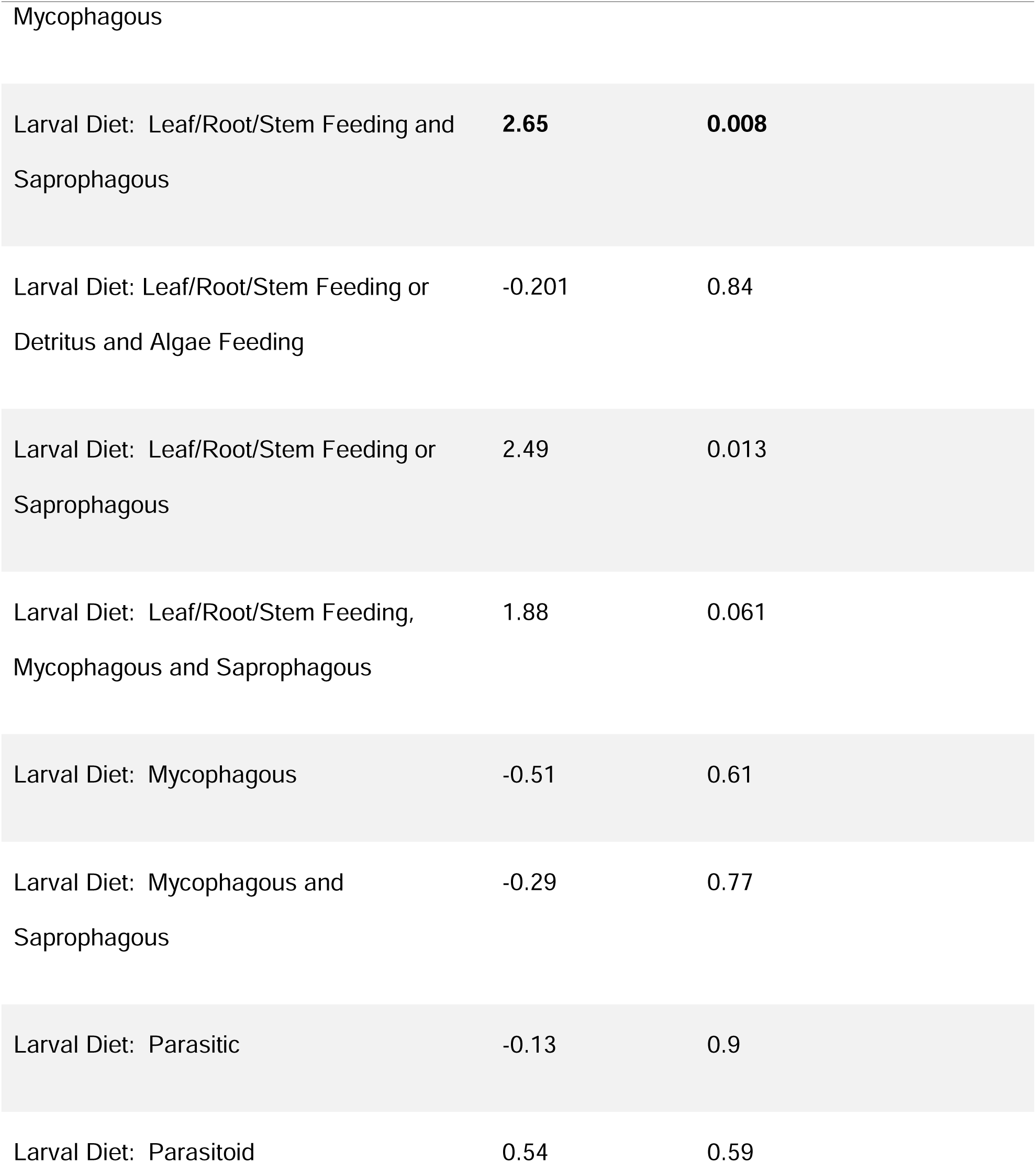
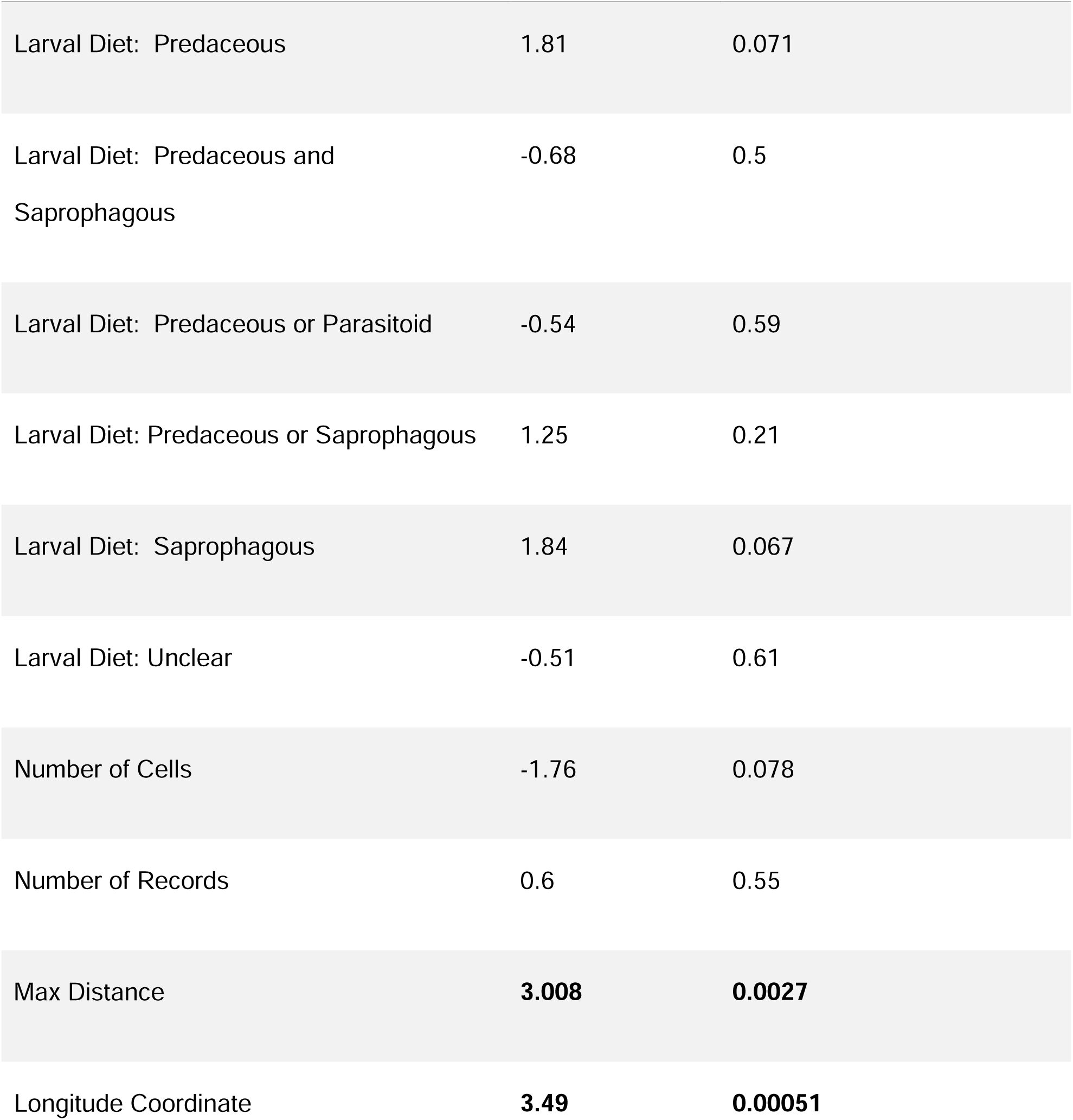
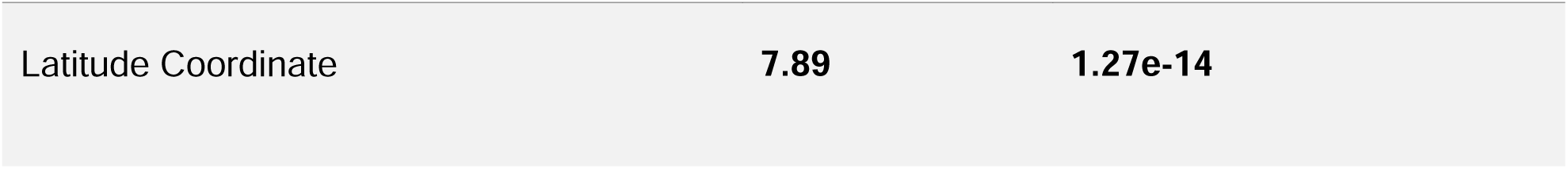
Results from the multiple regression testing whether traits and geographic variables are significant predictors of population genetic structure measures for Canada for a) Jost’s D, b) G_ST_, and c) F_ST_. Significant values are bolded.

When high levels of genetic differentiation were observed, this indicated low realized dispersal rates.

When using the non-linearized measures, a significant relationship between the indicators of population genetic structure and the distance between geographic regions was found for all measures, suggesting that the further the geographic distance, the higher the levels of genetic differentiation (Fig. 12). In F_ST_ and G_ST_, a positive relationship was found between the measures and the latitude coordinates of the center of the BINs’ range. F_ST_ also revealed a positive relationship with longitude. However, the multiple regression using the linearized measures revealed no significant relationship with distance, though the relationship with latitude and longitude and F_ST_ persisted.

**Figure 12:**
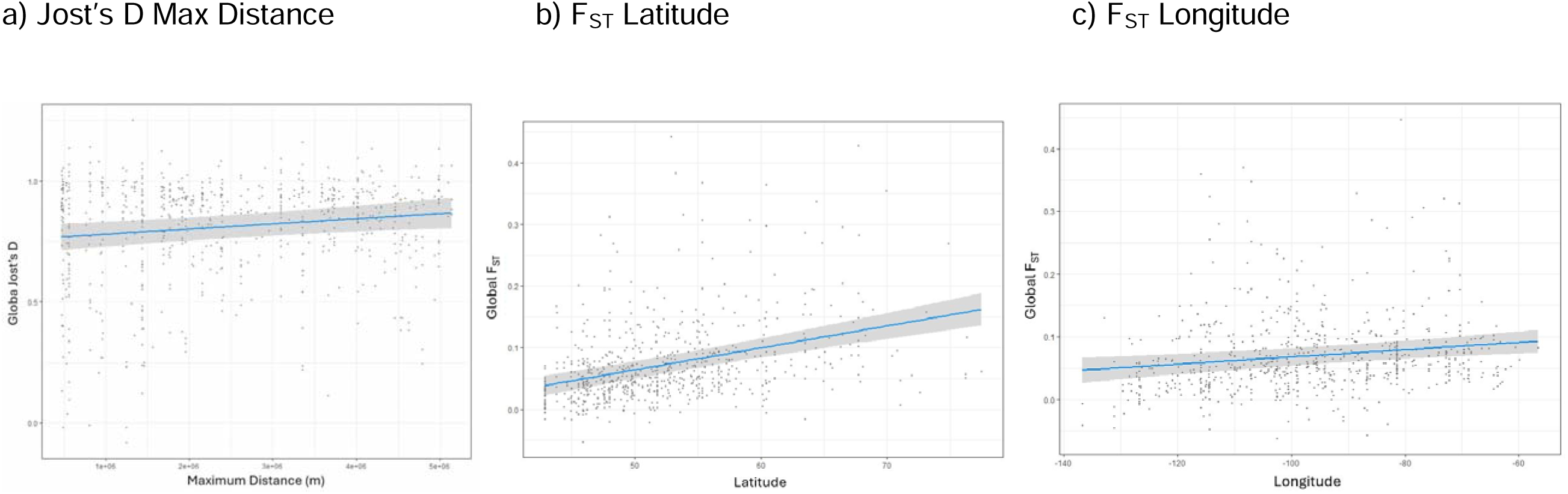
Plots showing the relationships between the global population genetic structure measures and geographic factors in the Canadian dataset of 716 Diptera BINs. Each dot represents a BIN. Only the significant relationships from the multiple regression are shown (Table 3). The relationship between indicators of population genetic structure and a) distance, b) latitude and c) longitude are shown. For all measures, the greater the distance, the greater the genetic differentiation. F_ST_ suggests that as latitude and longitude increase, so does genetic differentiation. The relationship with latitude was also seen in G_ST_.

Multiple regressions have a series of assumptions, so tests were performed to determine whether these assumptions were met. There were no violations of the assumptions of homoscedasticity and linearity. However, the assumptions of multicollinearity and normality were violated. The variance inflation factor was calculated and suggested that adult and larval diet are correlated, and this was confirmed by a chi square test (p-value =2.2x10^-16^). A Shapiro-Wilk test revealed a violation of normality (p-value = 2.2x10^-16^), but a Q-Q plot showed that this was not a large violation.

#### 3.2.3 Sensitivity Analysis

A sensitivity analysis investigating the impact of polygon size on the results was completed for the Canada dataset (Table 6). At a polygon size of 50 km across, there were only 662 BINs available for analysis after filtering, rather than 716 in the 500 km dataset; 105 polygons compared to 37; and 207,151 individuals compared to 225,774.

**Table 6:**
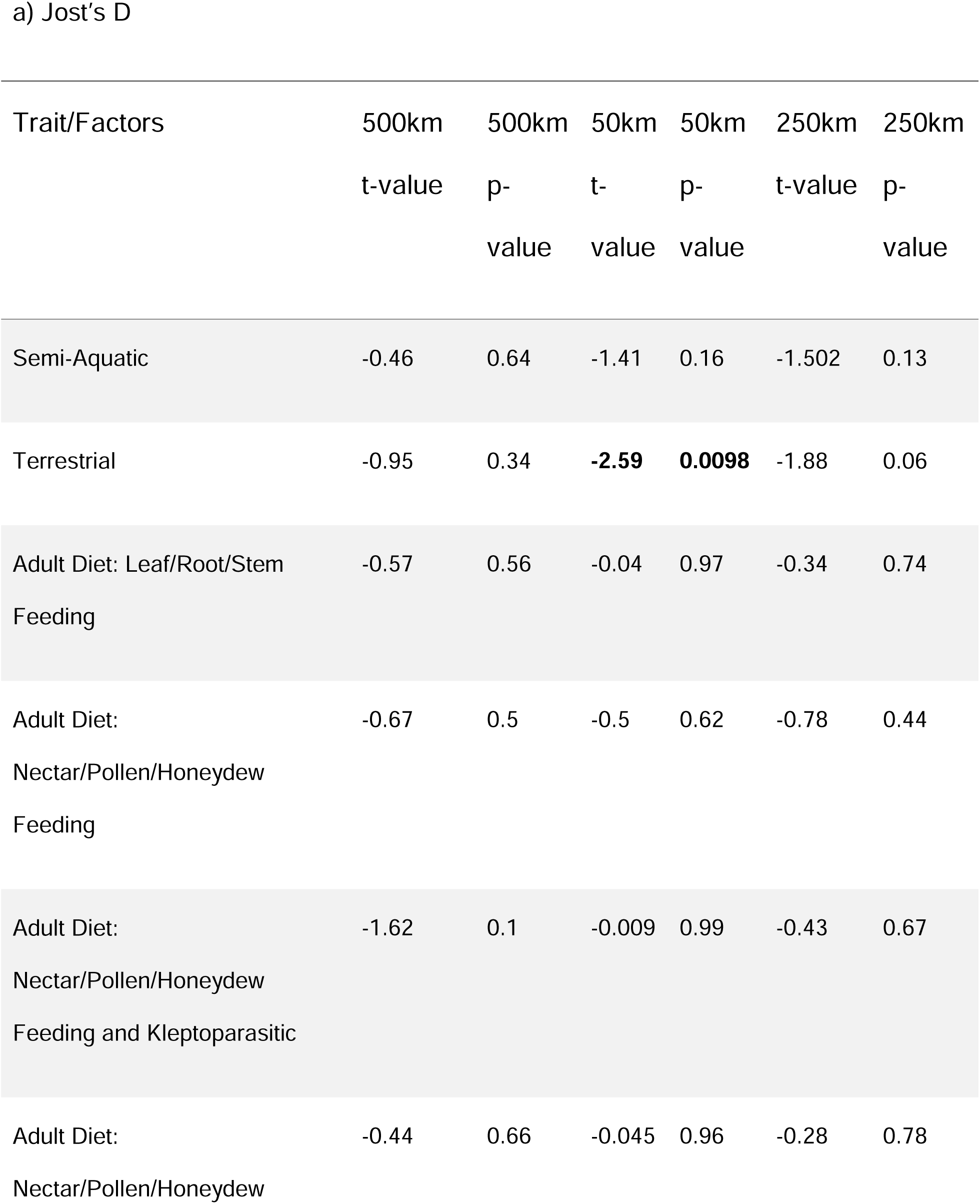
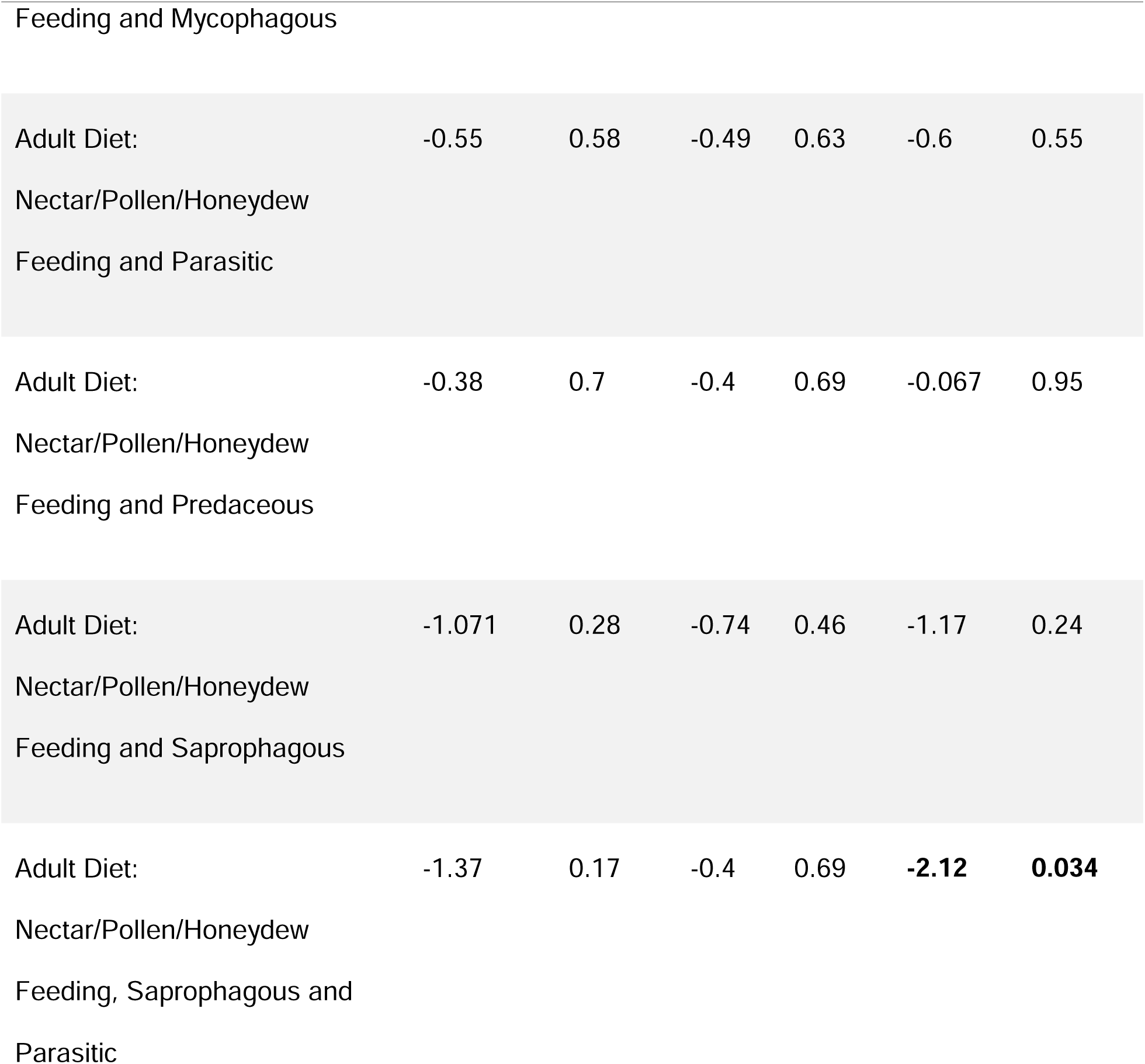
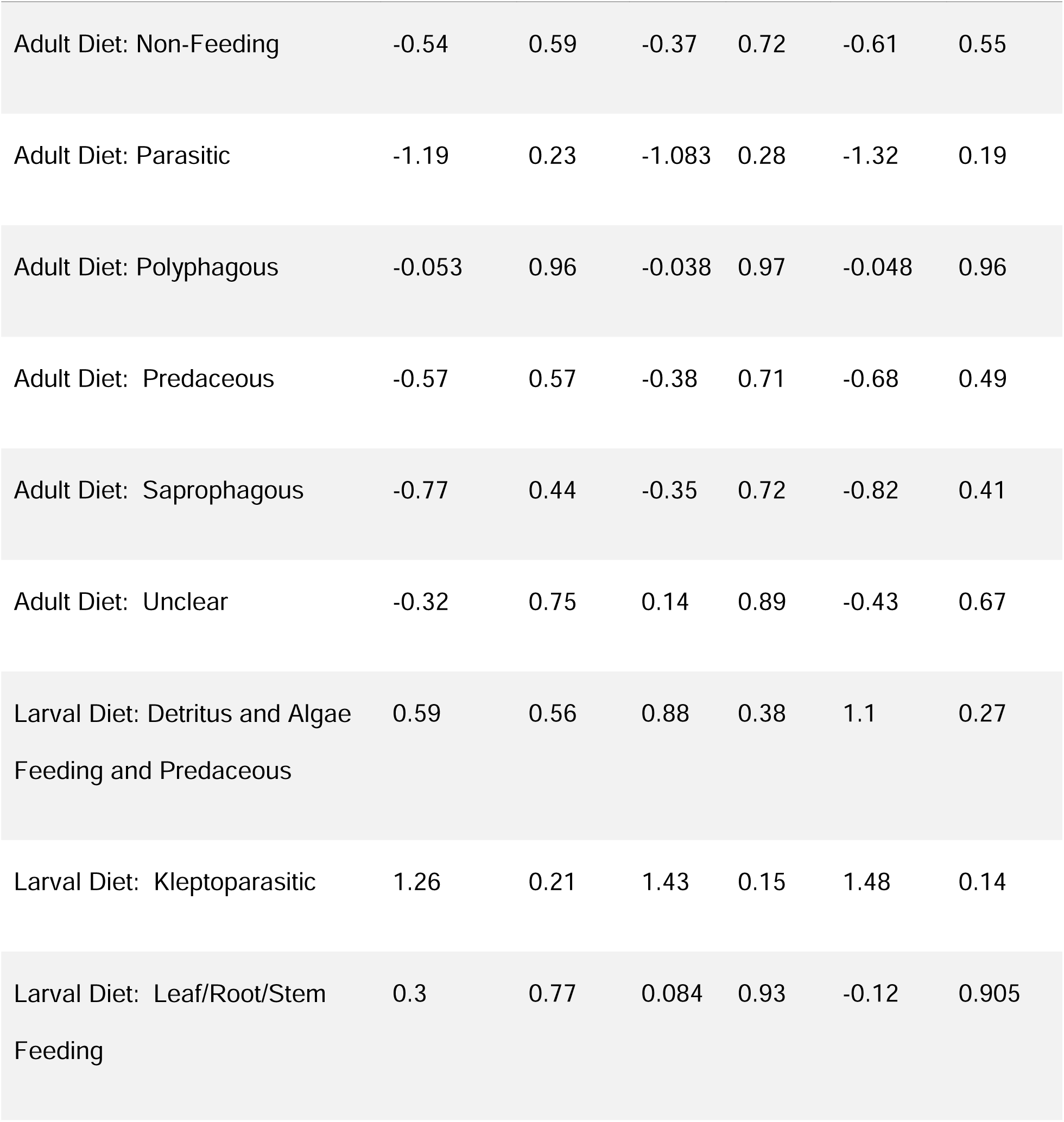
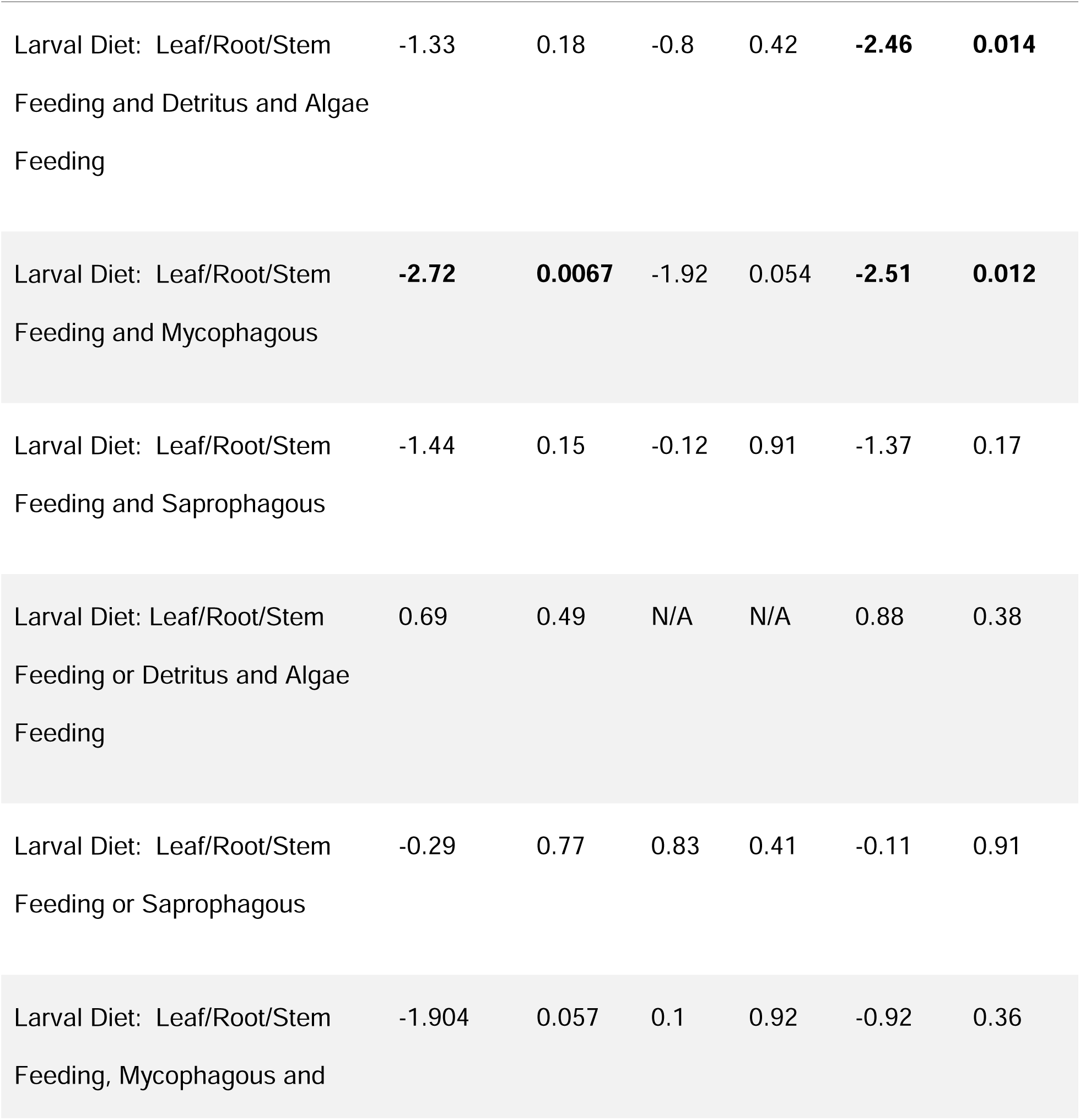
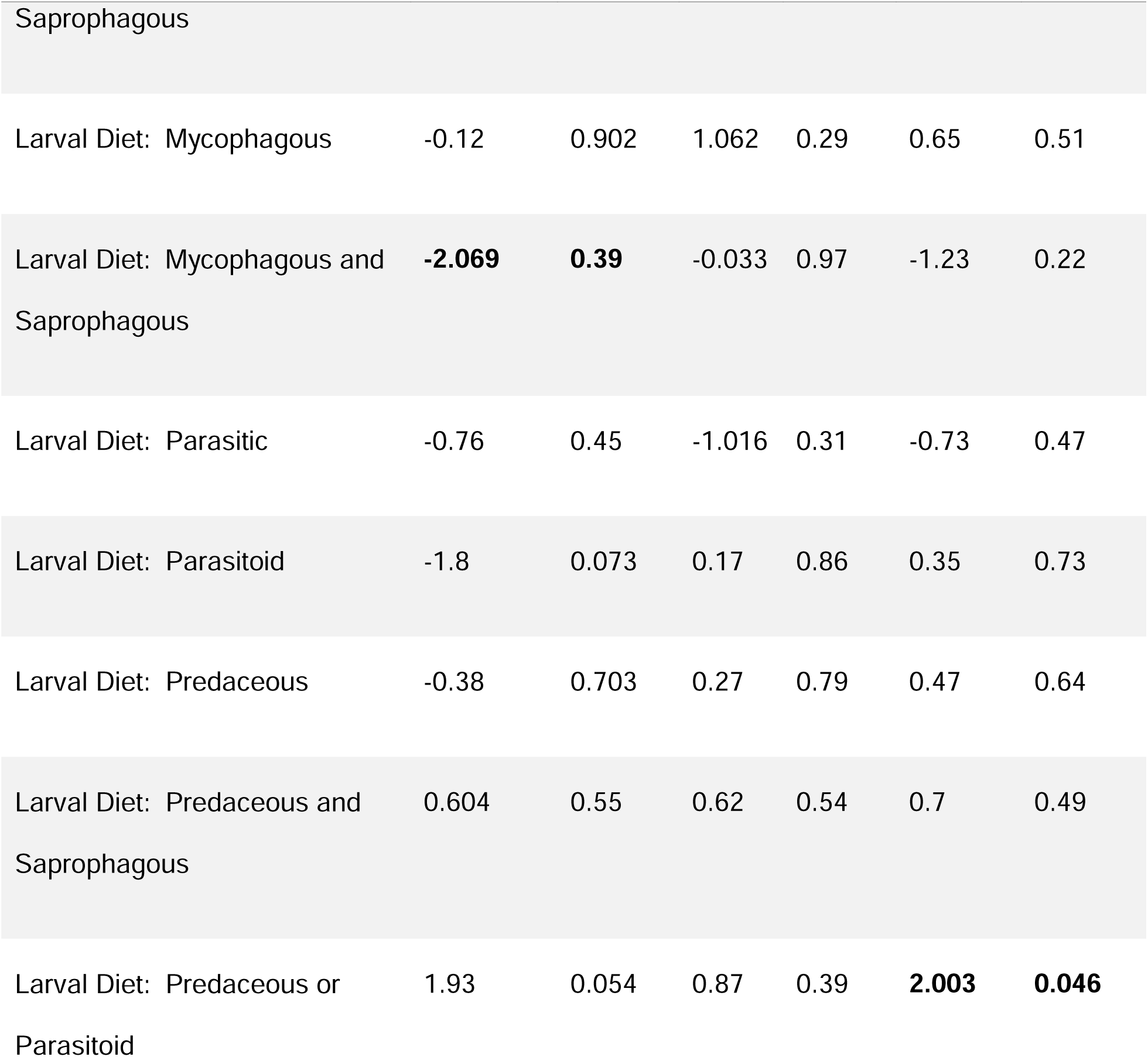
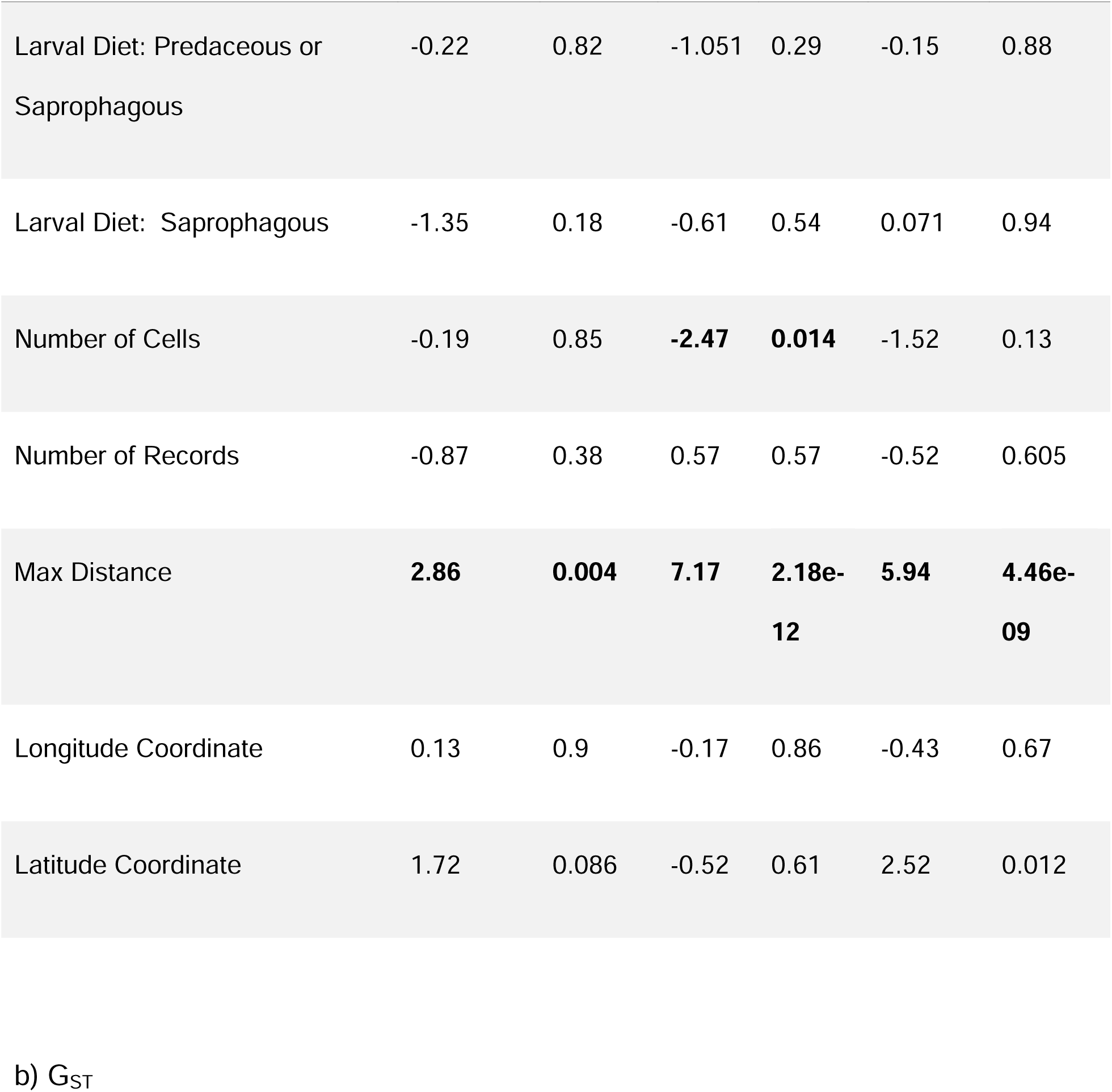
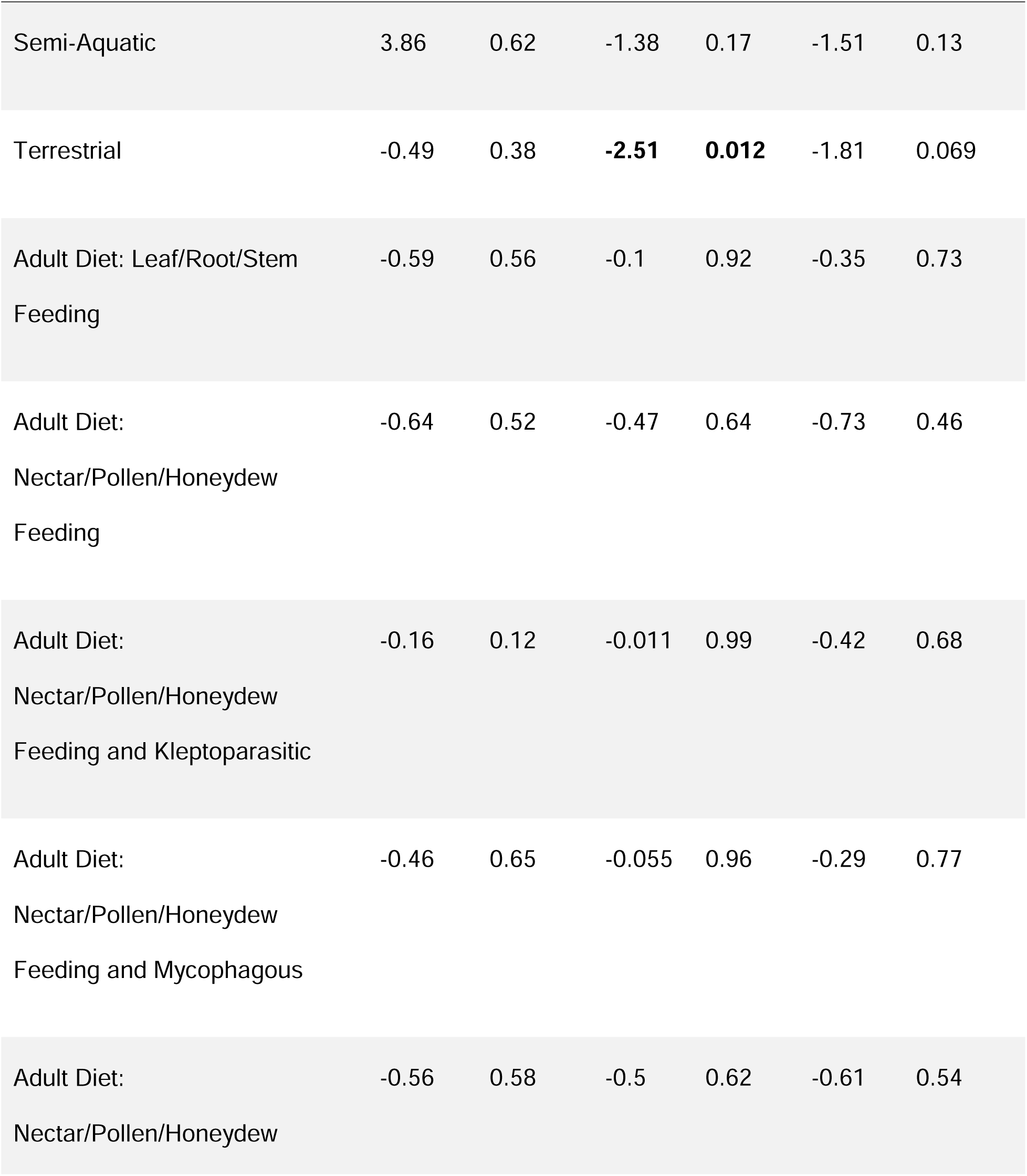
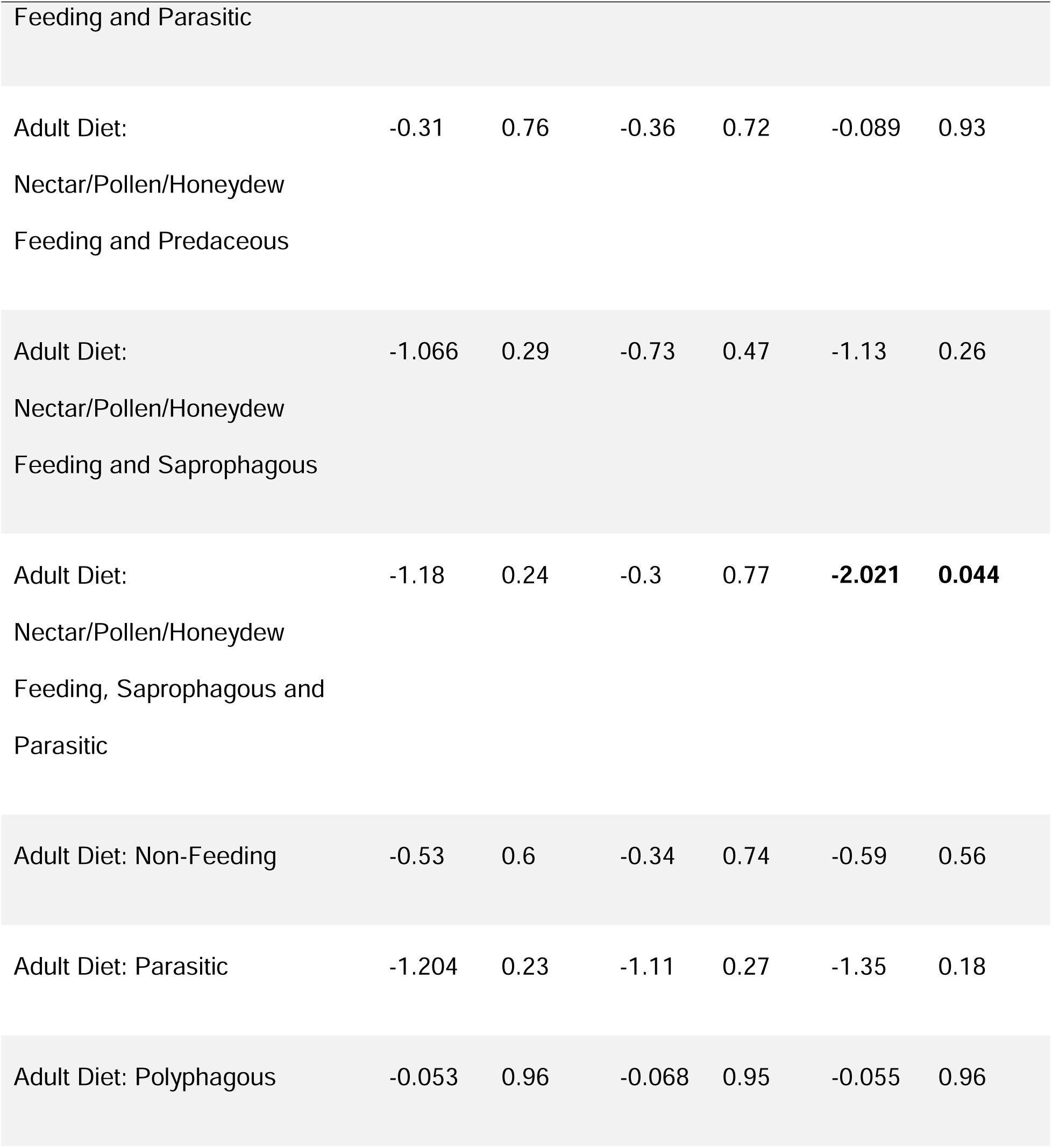
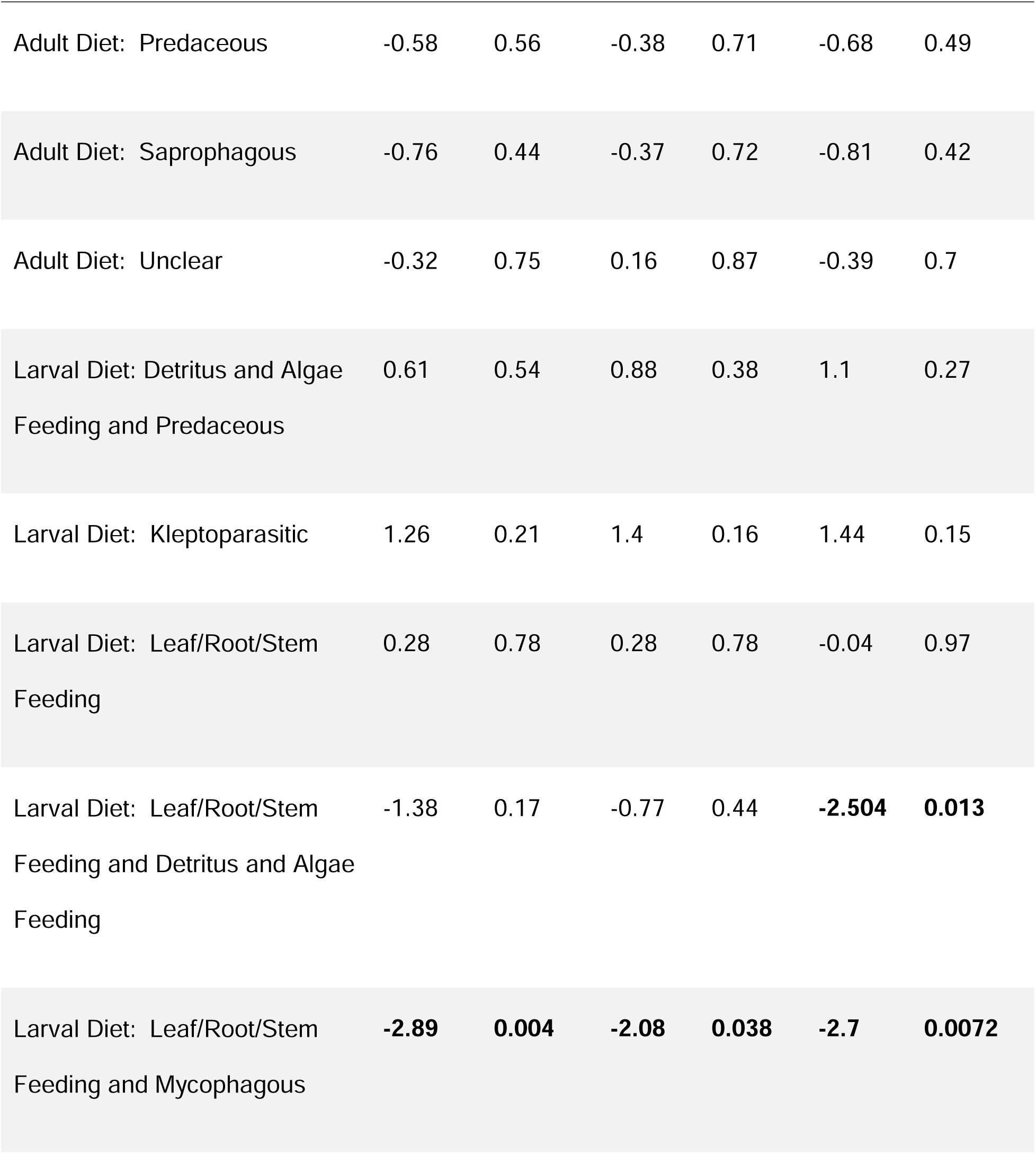
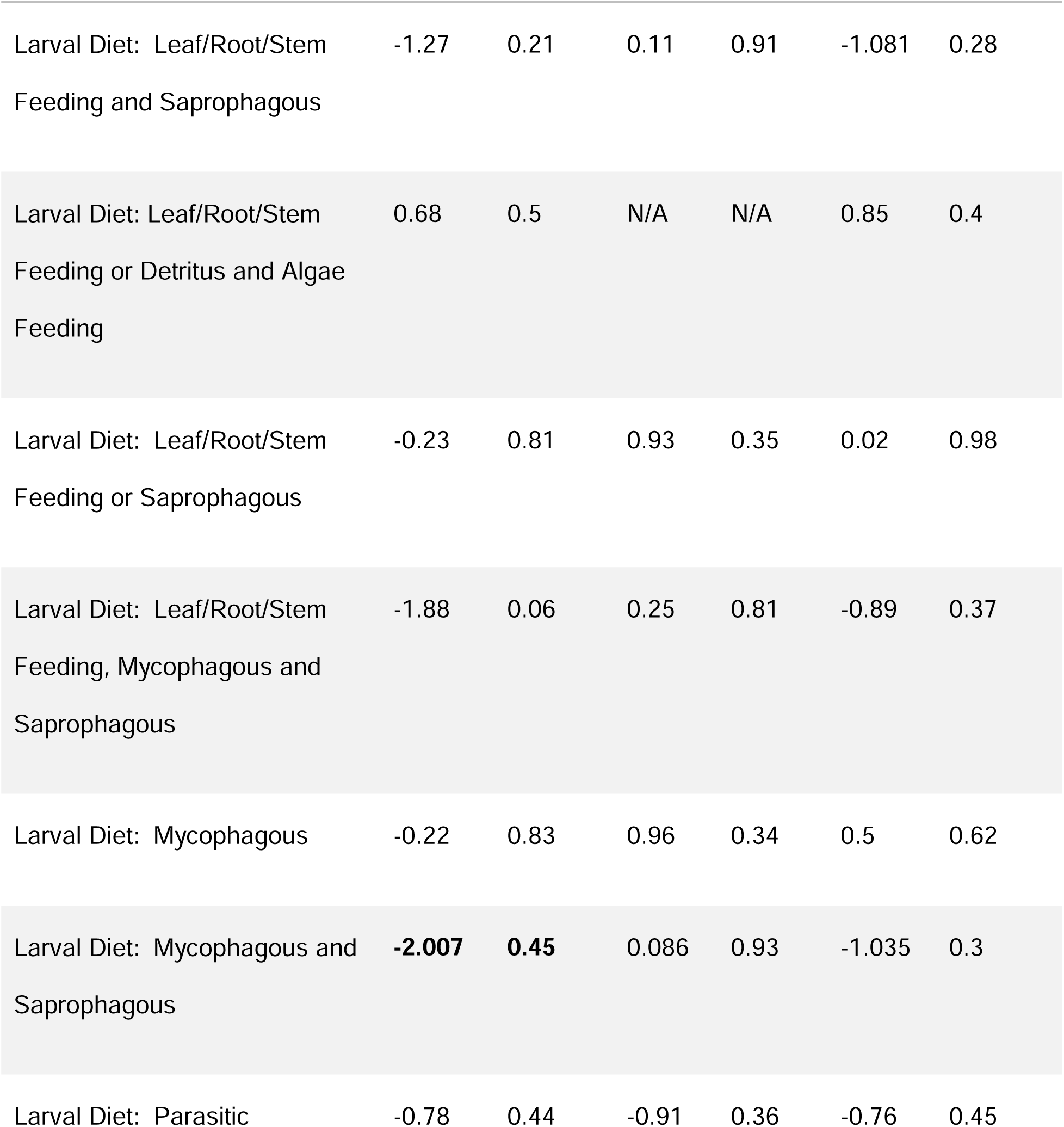
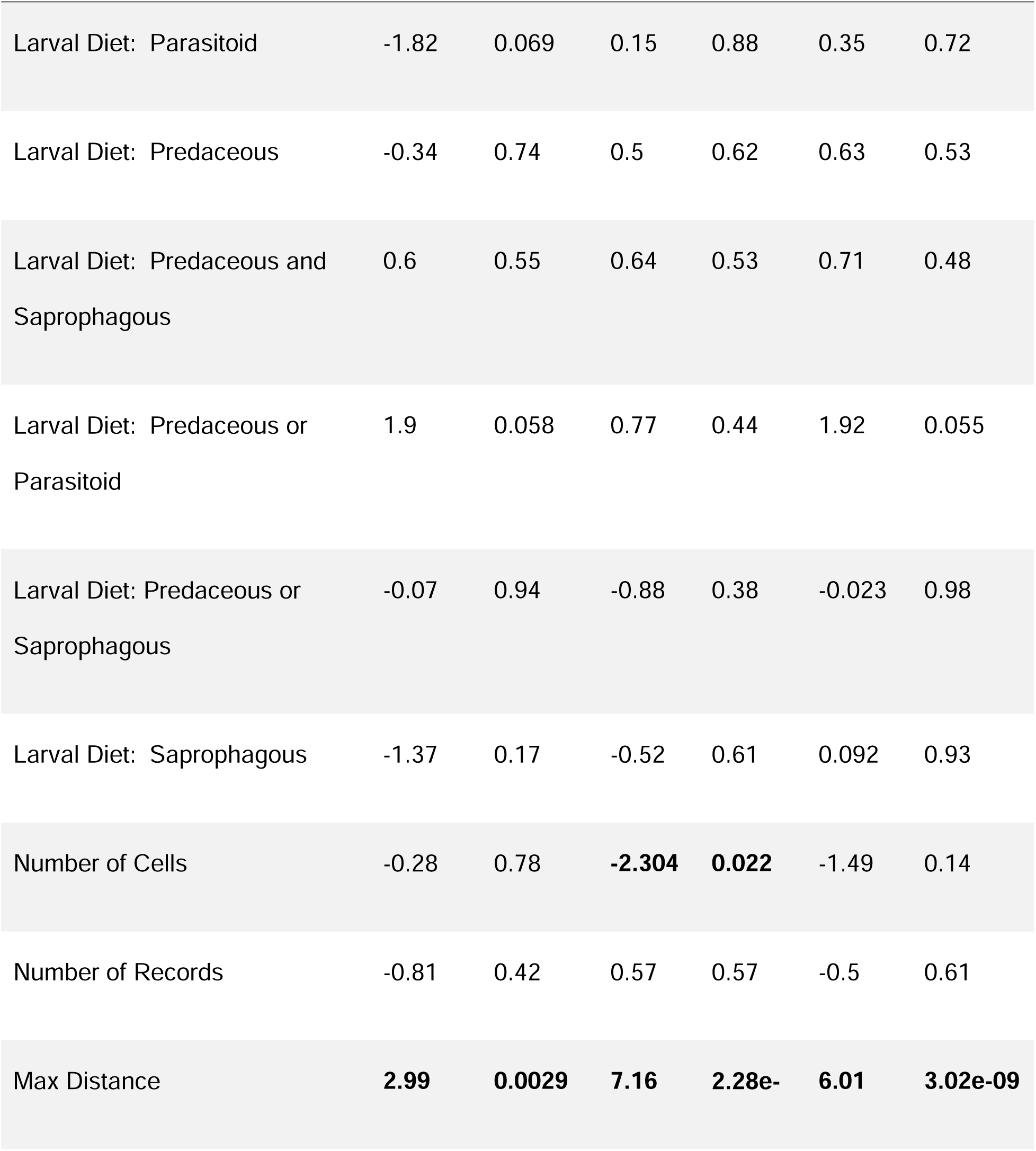
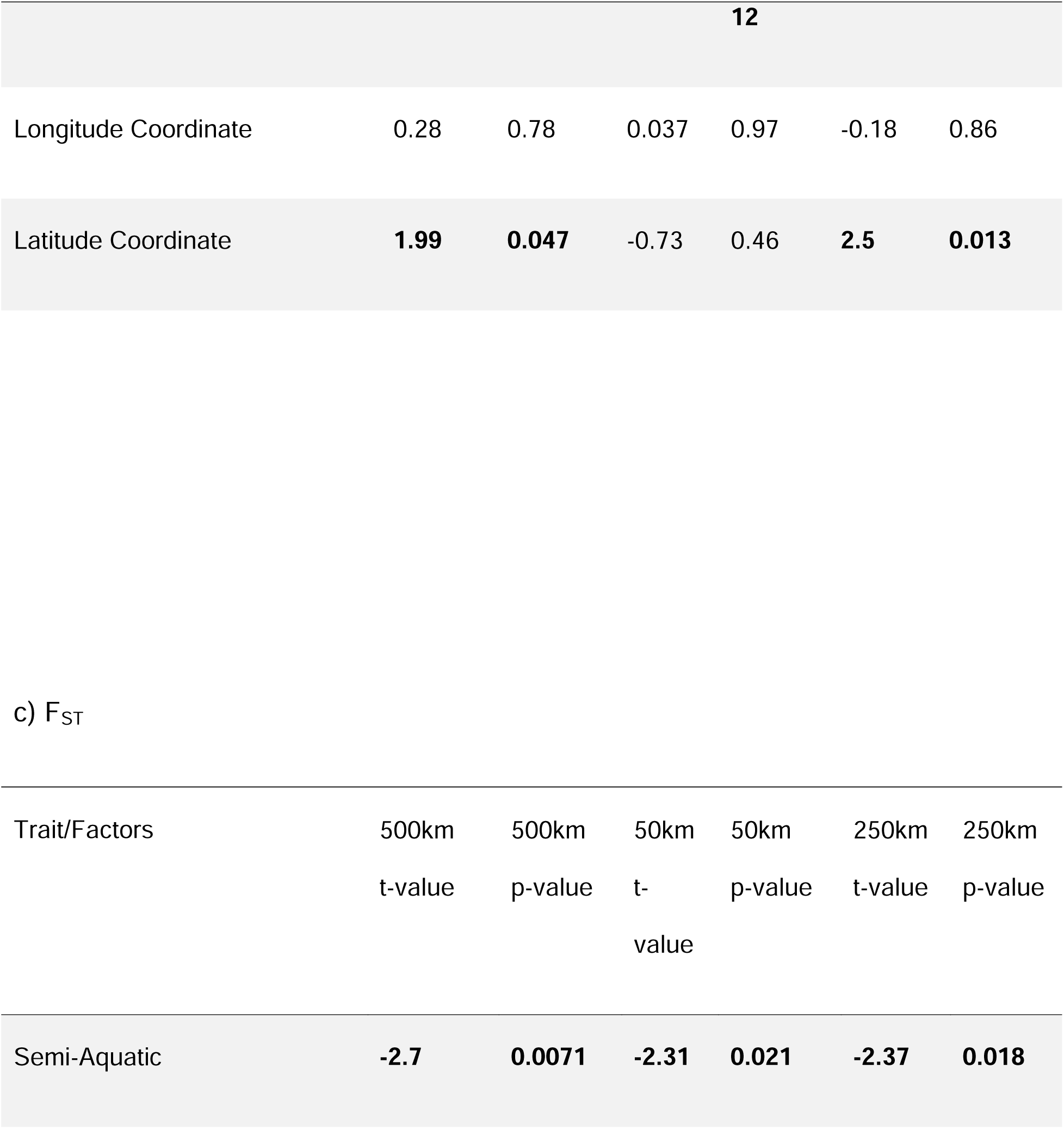
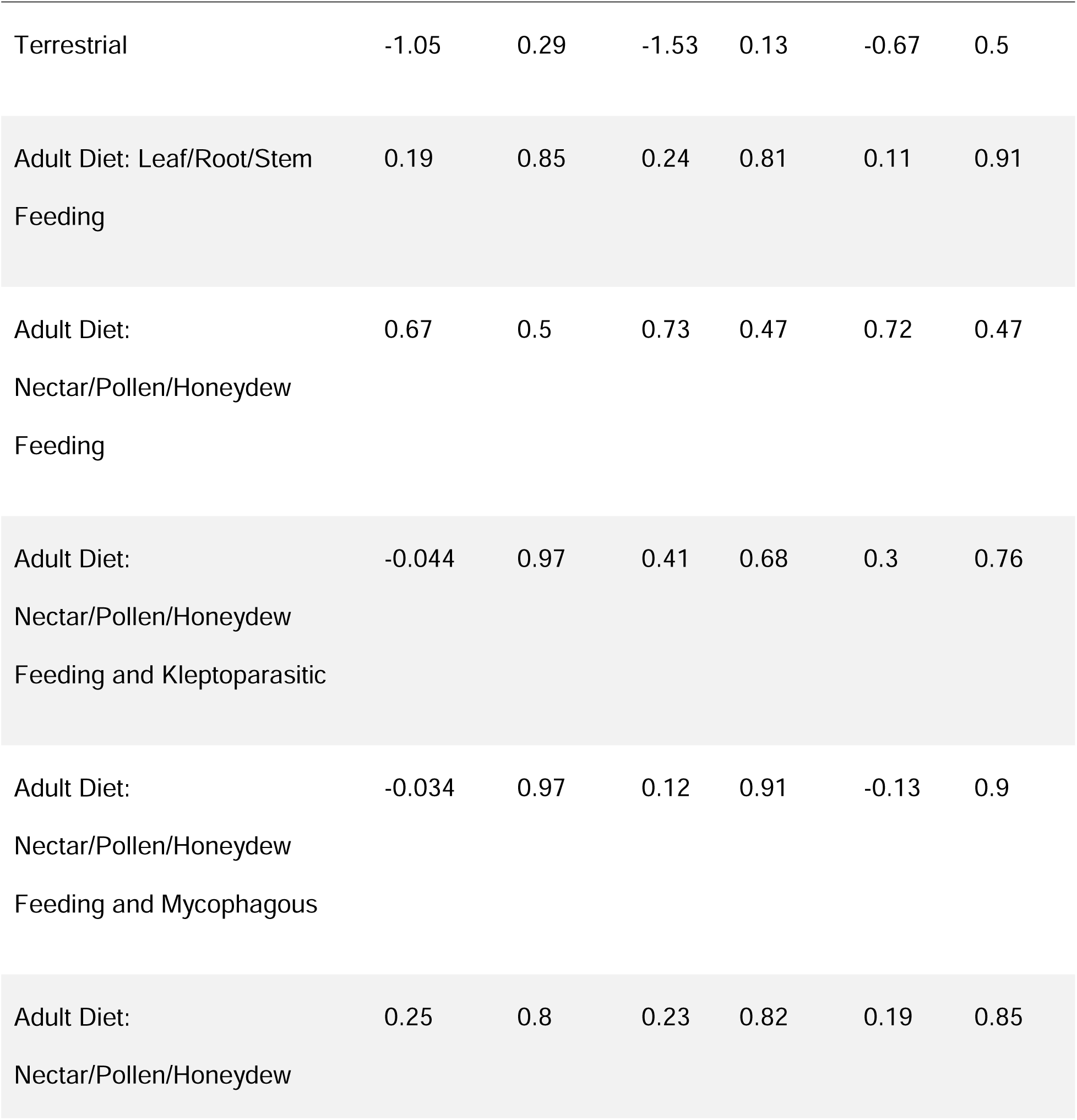
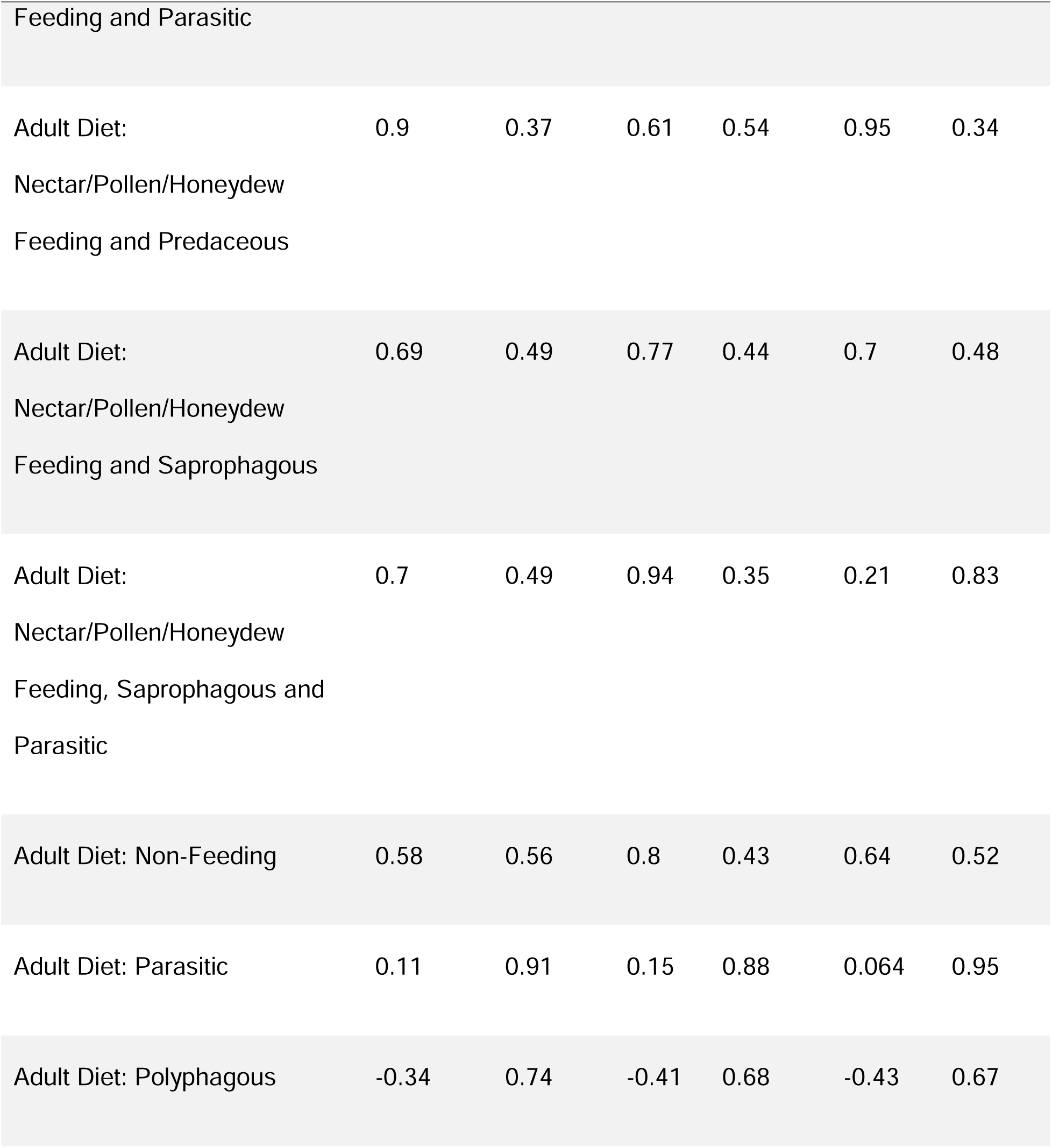
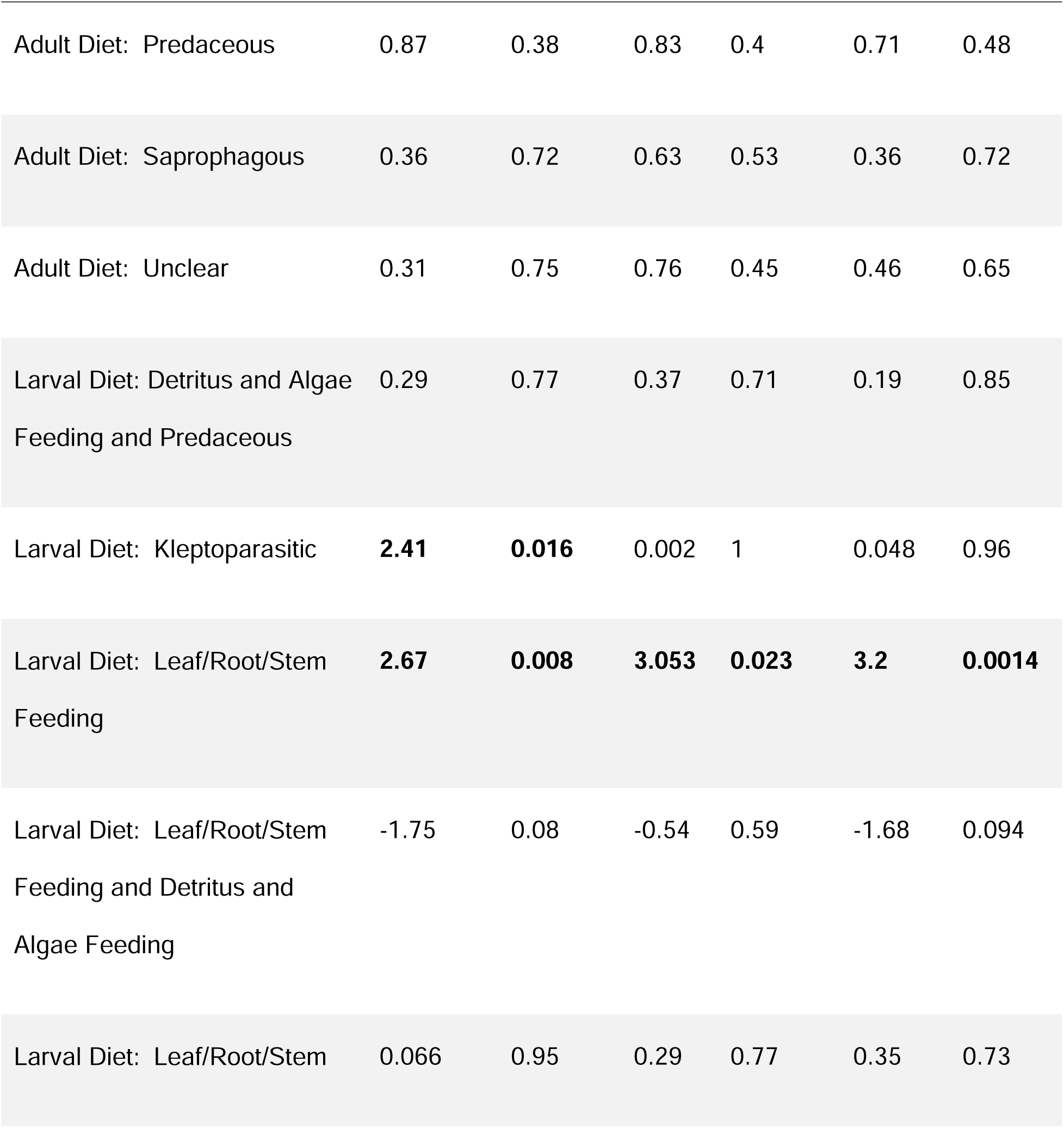
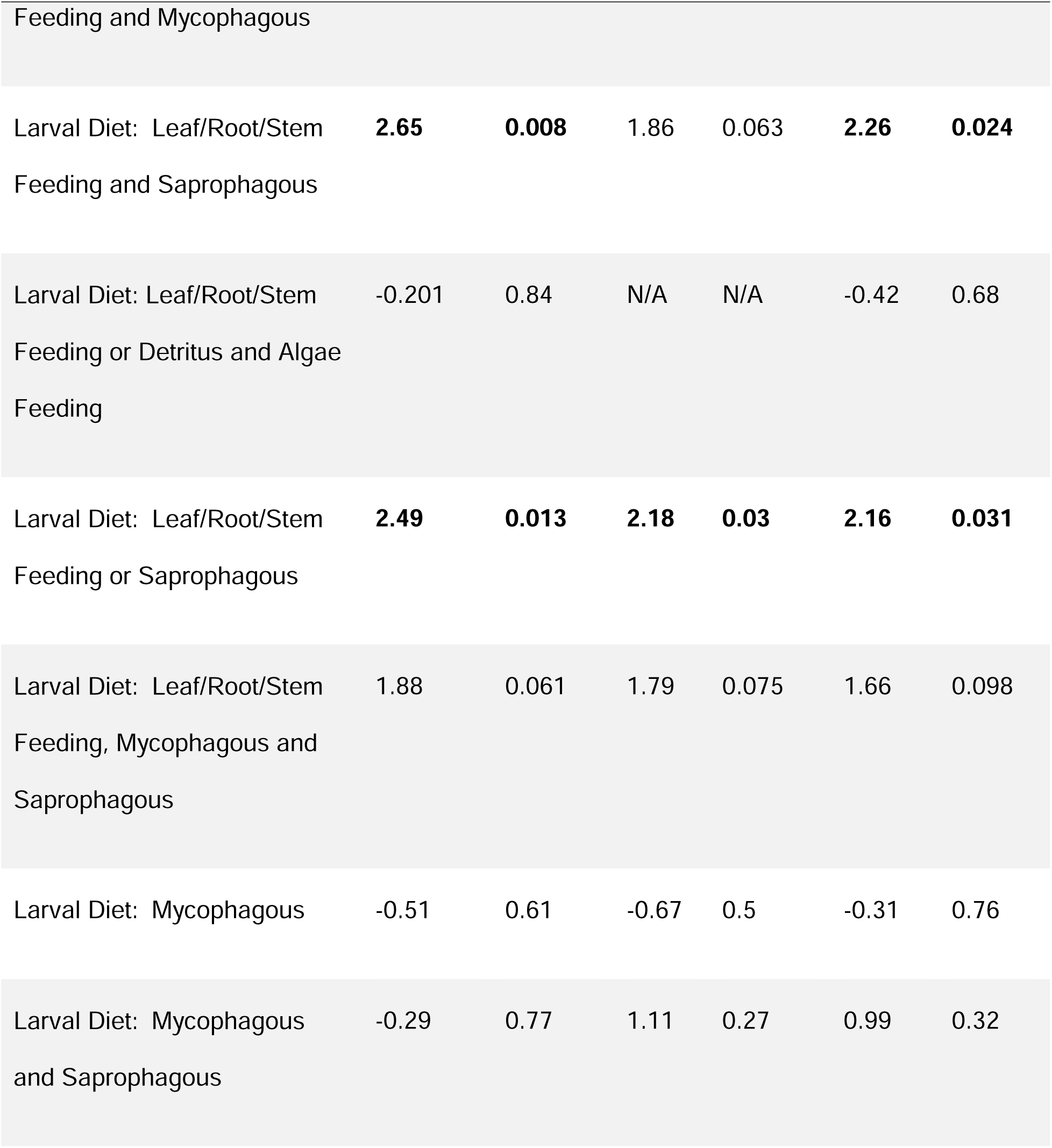
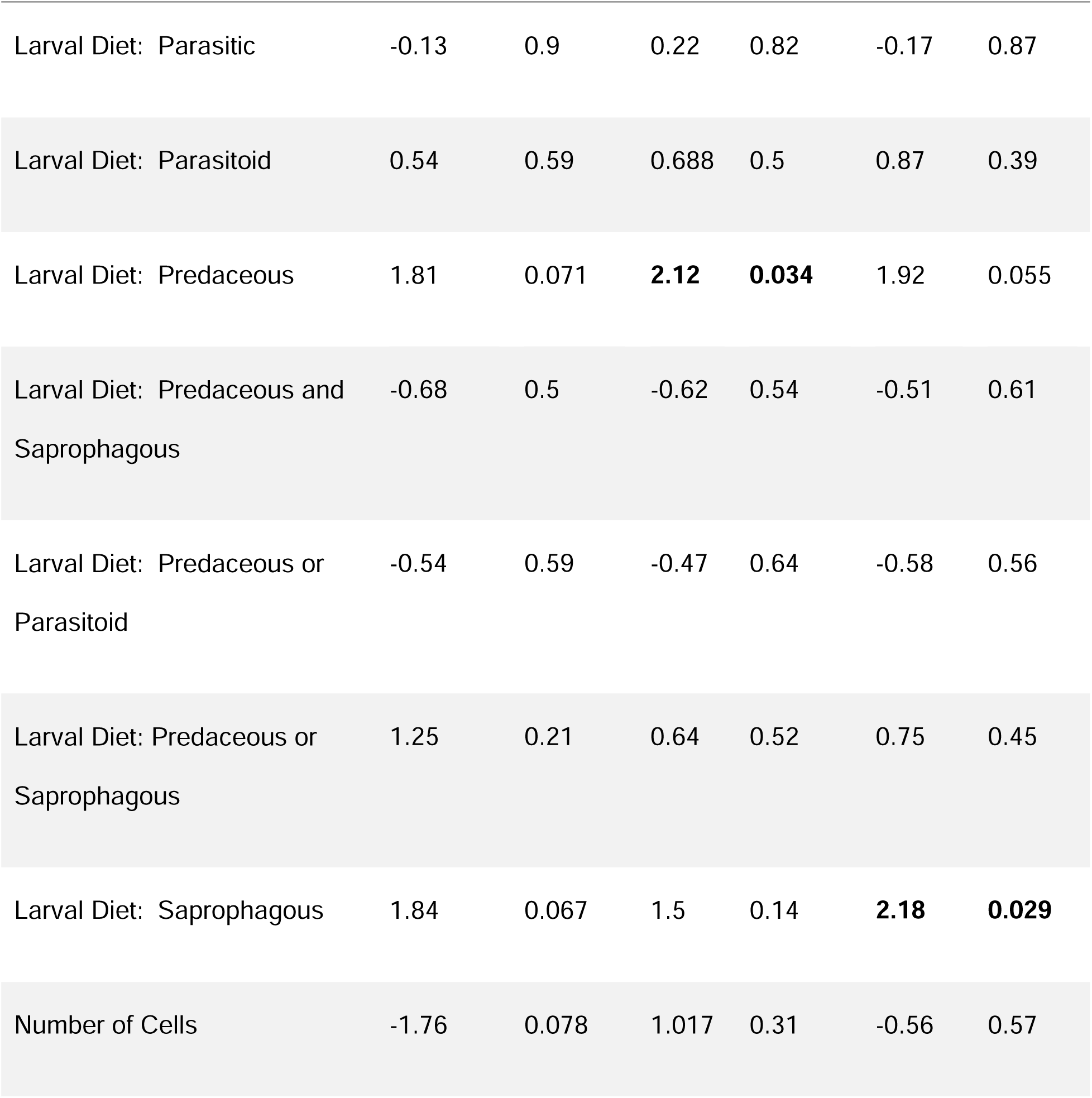
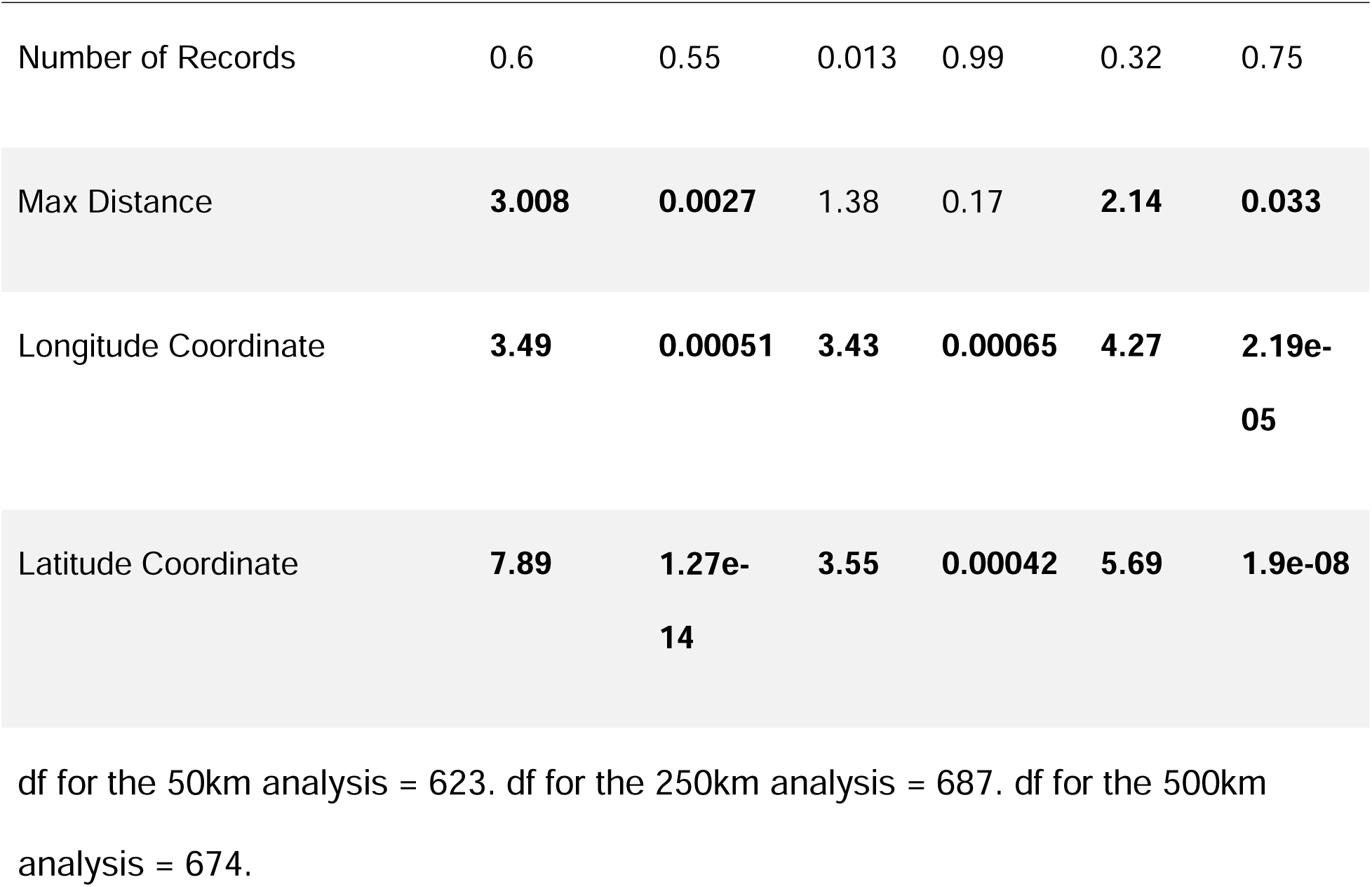
Results from the sensitivity analysis testing the influence of polygon size upon the relationship between population genetic measures and traits for a) Jost’s D, b) G_ST_, and c) F_ST_.

At 250 km across there were 727 BINs, 60 polygons, and 228,962 individuals.

For the 50km analysis, across all population genetic structure measures, results differed significantly from that of the 500km analysis using paired t-tests and including only those BINs present in both datasets (Jost’s D (p-value = 0.0083), G_ST_ (p-value = 0.00045), F_ST_ (p-value = 2.2x10^-16^)). For the trait analysis, results differed between 50km and 500km analysis. In the 50km analysis, a significant relationship was found with habitat in Jost’s D and G_ST_. Species found in terrestrial habitats had significantly lower levels of genetic differentiation. There was no longer a significant relationship with larval diet for Jost’s D, and for G_ST_ only leaf/root/stem feeding and mycophagous were significant. For F_ST_, a leaf/root/stem feeding larval diet and a diet that was a mix of leaf/root/stem feeding and saprophagous still showed significantly higher levels of genetic differentiation, and species with a predaceous larval diet also showed significantly higher levels. A significant relationship was still found between population genetic structure and geographic distance, except in F_ST_. In addition to this, with Jost’s D and G_ST_, a significant relationship was found with the number of cells in which a BIN was found. The more cells in which a BIN was found, the lower the levels of genetic differentiation. The relationship between latitude and longitude and F_ST_ remained the same.

For the 250km analysis, the population genetic structure measures significantly differed from the 500 km analysis only for G_ST_ (p-value = 0.032) and F_ST_ (p-value = 4.701x10^-13^). For G_ST_ and Jost’s D, a larval diet that is both mycophagous and saprophagous no longer showed significantly lower levels of genetic differentiation, but an adult diet that is nectar/pollen/honeydew feeding, saprophagous and parasitic, and a larval diet that is leaf/root/stem feeding and detritus and algae feeding showed significantly lower levels. Also, a larval diet that is predaceous or parasitoid showed significantly higher levels of genetic differentiation in G_ST_ only. For F_ST_, the relationship with diet remained consistent with the 500km analysis for BINs, except there was no longer a significant relationship between the population genetic structure measures and BINs with a kleptoparasitic larval diet. A new positive relationship was found with BINs that had a saprophagous larval stage.

## 4 Discussion

### 4.1 A Novel Pipeline for Large-Scale Comparative Population Genetics

The focus of this study was to create a publicly available pipeline connecting a set of modules to determine indicators of population genetic structure for a variety of taxa and compare these measures across species possessing differing biological traits.

Throughout development, significant effort was made to adhere to best practices and ensure the modules were easily adaptable and useful for other researchers. Fifteen modules were included in this study. These modules can be used together to recreate the analysis we performed or used independently to suit the needs of the researcher. The modules included steps to extract data from various publicly available databases, such as BOLD and GBIF, to filter and group the data into polygons, to determine measures of population genetic structure, and to compare these measures across species with differing habitats and diets using an ANOVA, PGLS, and multiple regression. The first module includes steps to set up and adjust the important variables for all other modules. Other variables are also adjustable throughout the script. The modules provide options to deal with datasets of varying sizes, and also provide options for different analyses based on the dataset and research needs. This novel pipeline provides a way to determine and compare population genetic structure measures for a variety of taxa and geographic regions. The utility of the modules was demonstrated through the case study on Diptera species. This case study shows the power and potential of the pipeline to generate interesting patterns and draw biological conclusions at a large scale.

### 4.2 Case Study

To test the modules, the analysis was run with Diptera species from Greenland and Canada. This analysis was done using BINs as well as a 4% clustering threshold to create Operational Taxonomic Units. Several studies have found strong correlation between BINs and species in Diptera [80–82]. For example, when Bartolini et al. [80] sorted morphological *Anastrepha* (Diptera: Tephritidae) species into BINs, they created 14 BINs for the 16 species represented. Hernandez-Triana et al. [81] created 29 BINs for 28 Simuliidae species, and Yu et al. [82] sorted 5 Chironomidae species into seven BINs. BINs have also been shown to correspond to species in other arthropod groups, such as Coleoptera and Aphididae, where BINs matched with species 90% of the time [83–85], and Lepidoptera, where BINs had a 70% concordance with species [86].

Most species/BINs/clusters showed low levels of genetic differentiation in Greenland, and there was no significant relationship between the biological traits and the population genetic structure measures. Results from Canada were more mixed. There were stronger disagreements between Jost’s D and G_ST_, and F_ST_ when compared to Greenland, with G_ST_ and Jost’s D suggesting high levels of genetic differentiation and F_ST_ suggesting lower. There was a significant relationship between the population genetic structure measures and larval diet, with those with a leaf/root/stem feeding and mycophagous larval stage or a larval stage that was a mix of mycophagous and saprophagous having lower levels of genetic differentiation. There was also a significant relationship between the indicators of population genetic structure and distance between populations. As the geographic distance increased, so did the genetic differentiation.

The strength of the Greenland analysis lies in the fact that Greenland provides an example of the minimum amount of data required to run the modules and analysis. By using this small dataset, we were able to show the script’s functionality. To run the modules, at least two geographic regions/polygons and two BINs (or clusters or species) are required. When running a large analysis, it is beneficial to run the modules first on the minimal dataset size. By beginning with a sampled-down version of a dataset, a researcher can ensure the modules work for the dataset structure, intended research, and computing environment, before scaling up to a dataset size more appropriate for drawing biological conclusions. The Greenland dataset analysis ran without error and yielded reasonable results. Since the two available populations for Greenland were located in the same area and lacked any noticeable physical barrier, there is likely a lot of gene flow between these populations and a large crossover of genetic information. It is also likely that recently deglaciated habitats in Greenland were colonized by a modest number of individuals and then the population spread to adjacent areas. This geography and history were supported by the population genetic structure measures, most of which showed very low levels of genetic differentiation between the populations. This was also supported by the AMOVA and mismatch analysis, both of which suggested the populations are not genetically isolated. Due to the module’s success with analyzing this small dataset size, we were able to scale up to a larger dataset and run our case study analysis on Diptera species from Canada.

The Canada dataset was much larger than that of Greenland. Both G_ST_ and Jost’s D suggested that there is genetic differentiation between regions in Canada. This could be due to the geographical distance and space, as the regions in the Canada dataset were widespread and covered various geographic regions and habitat types across Canada. Genetic differentiation has been shown in Canadian fly species in other studies, such as Maxwell et al. [87], who found significant differentiation among regions in British Columbia and evidence of isolation by distance. They also investigated genetic differentiation using AMOVA and F_ST_ [87]. The results of the AMOVA and of F_ST_ differed between our analyses [87], with lack of genetic differentiation being observed in our study. However, the other population genetic structure measures, G_ST_ and Jost’s D, did detect genetic differentiation and aligned with the findings of Maxwell et al. [87]. Maxwell et al. [87] used microsatellites instead of COI, and the species included in their analyses, *Rhagoletis indifferens,* was not included in our analysis. The genetic differentiation observed by Maxwell et al. [87] in F_ST_ could be due to the higher rate of change in microsatellites compared to COI. Despite these differences, both studies found similar results and evidence of genetic differentiation and isolation by distance [87]. Our modules were not only able to draw conclusions about one species, but general trends across hundreds of Diptera species, making use of public databases and DNA barcoding data. This comparison shows the potential of our modules to determine indicators of population genetic structure and produce results that are supported by the current literature.

F_ST_ suggested lower levels of genetic differentiation. Low levels of genetic differentiation are not uncommon in Diptera. Populations have shown low genetic differentiation and high population connectivity in various regions at large geographic scales, including across Southeast Asia [88] and across several European countries [89].

While we will not describe the results in detail for each species in this study, we will highlight some of the results for the best-sampled species in Canada. The best-sampled BIN from our study was BOLD:AAD1720 and represented the species *Limnophyes asquamatus* with 9086 records in Canada found across 20 geographic regions. The global population genetic structure measures, AMOVA and mismatch distribution, reflect the general conclusions of our study, with Jost’s D and G_ST_ suggesting high levels of genetic differentiation and F_ST_ suggesting lower levels. The population genetic structure measures for each region pair are shown in Figure 13.

**Figure 13:**
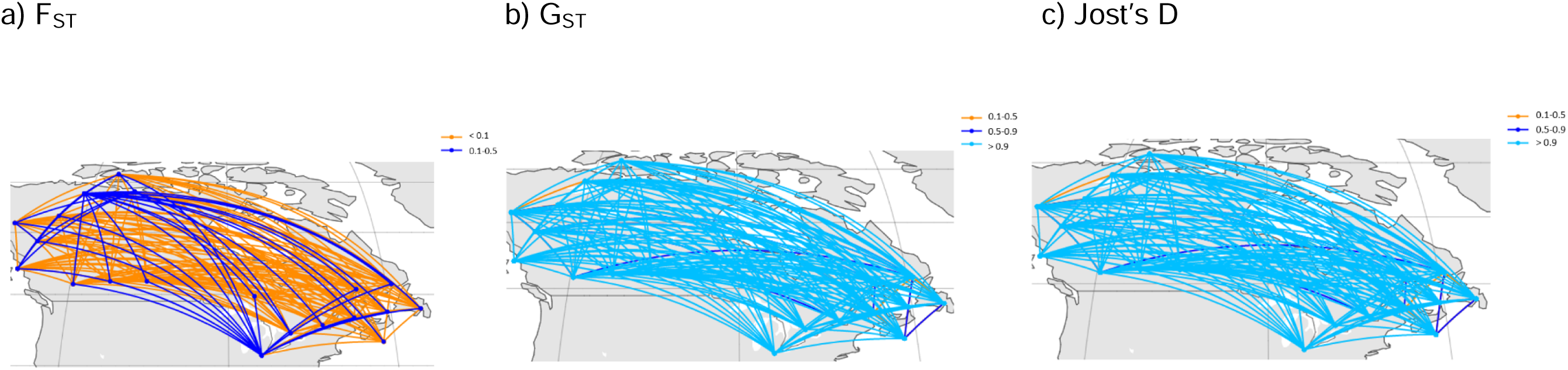
The pairwise a) F_ST_, b) G_ST_, and c) Jost’s D between geographic regions in Canada for BIN number BOLD:AAD1720 representing the species *Limnophyes asquamatus*. Jost’s D and G_ST_ suggest high levels of genetic differentiation between most regions, while F_ST_ suggests lower levels. For G_ST_ and Jost’s D, the lowest population genetic structure values are between regions that are fairly close geographically.

When looking at the measures between regions, it appears that, for G_ST_ and Jost’s D, the lowest population genetic structure values are between regions that are fairly close geographically. The 10 largest BINs and species included in this study (BOLD:AAA3453, BOLD:AAB0377, BOLD:AAC8842, BOLD:AAD0853, BOLD:AAD1720, BOLD:AAH3920, BOLD:AAU8534, BOLD:ABU5525, BOLD:ADA3072, BOLD:ADX9795) have been used to study Diptera diversity across regions, the effectiveness of methods, community composition, phylogenetics, and protein interactions in COI [18, 40, 48–49, 90–97]. Through this case study, we were able to use these data in a new way and investigate the population genetic structure of these BINs.

Not only did we determine levels of genetic differentiation, but we also compared these across Diptera species possessing different biological traits. All known Diptera species that met our sampling requirements were included, and results were found across hundreds of taxa at a Canadian scale. The modules were successful in uncovering interesting relationships between the indicators of population genetic structure and larval diet. Jost’s D and G_ST_ found that species with a leaf/root/stem feeding and mycophagous life stage and those that had a larval stage that was a mix of mycophagous and saprophagous had significantly less genetic differentiation than Diptera with other diets. F_ST_ also displayed several different patterns. F_ST_ suggested higher levels of genetic differentiation in larvae with a kleptoparasitic diet, a leaf/root/stem feeding diet, a diet that is a combination of leaf/root/stem feeding and saprophagous, or a diet that is saprophagous or leaf/root/stem feeding. It is possible that these species rely on a certain species of plants or fungi, and therefore may not be able to disperse and be as connected and widespread as those with a more generalist diet.

The pipeline not only uncovered relationships between indicators of population genetic structure and biological traits, but also showed its ability to detect significant patterns involving other important factors through multivariate analysis. By including these in the modules, we accounted for other variables that may be affecting indicators of population genetic structure and infer whether the patterns discussed earlier were solely due to biological traits. The G_ST_, Jost’s D, and F_ST_ all found that as the geographical distance increases, so does the genetic differentiation. F_ST_ and G_ST_ also exhibited an additional relationship between the latitude coordinates of the centre of each species’ range and the indicators of population genetic structure, and F_ST_ found the same pattern in longitude. The geographic distance and differences in barriers and climate that occur across latitude and longitude within Canada likely influenced the geneflow and connectivity of populations, resulting in differences in genetic differentiation. Historical factors, such as population separation in different glacial refugial areas, likely also affect patterns in Northern regions. These results were not supported, however, when linearized versions of the measures were used.

Several analyses were performed in this study with variations in important factors, such as how species or clusters were defined and the size of the geographic regions being investigated. Even within one analysis, the results between the population genetic structure measures differed. For the Canada dataset, F_ST_ produced much different results than G_ST_ and Jost’s D. This is due to the different way in which F_ST_ and G_ST_ and Jost’s D are calculated. Often for F_ST_, the value does not actually range between 0 and 1, but rather between 0 and a value determined by the heterozygosity in the data [66]. G_ST_ and Jost’s D use a standardized range of 0 – 1 [66]. It is recommended to use G_ST_ and Jost’s D in addition to F_ST_, as all of these measures can be informative [66]. The module is easily adaptable, so if some measures are better suited to a given researcher’s analysis, the option is there to include some measures and exclude others. The parameters of the functions used to calculate these measures could also be adjusted, and a sensitivity analysis could be run to see how these measures and the discrepancies between them are affected. There were also differences between the BIN and cluster analysis. This suggests that it is important to consider which method is best for the analysis and dataset. Results also differed when geographic regions of different sizes were used. It is important to consider what cell size is best for the geographic region and taxa under investigation.

To summarize the main results, in Canada evidence of genetic differentiation was detected, and there were significant relationships between the indicators of population genetic structure and habitat and larval diet. There was also significant evidence of isolation by distance and a relationship between indicators of population genetic structure and latitude and longitude. These results support our hypothesis that biological traits influence indicators of realized dispersal rates and population genetic structure, as they play an important role in the overall dispersal process. This case study provides a good demonstration of the utility of the pipeline as a whole, and of the modules individually. A variety of options are provided to account for different sizes of datasets and preferences and needs of the user. The modules ran efficiently and consistently, allowing results to be generated and biological conclusions to be drawn. Flies are an ecologically important group, having large effects on agriculture, public health, and ecosystem function [98–102]. By applying these modules to taxa such as flies, we can draw biological conclusions that are of interest to diverse groups of researchers and the general public.

### 4.3 Suitable Data for Usage

Though the use of COI has some limitations, such as the limitations in its ability to infer deep phylogenetic relationships [17,103], this marker was suitable for our analysis. COI is frequently concordant with other types of genetic data for detecting genetic variability and population genetic structure. Meriam et al. [33] suggested that COI performs better than 12S and 16S ribosomal RNA (rRNA) when inferring population genetic structure, due to its greater evolutionary rate. Choi et al. [34] and Stark et al.

[30] also found similar results between COI and 16S rRNA when investigating genetic diversity, phylogeographic history, and phylogenetic relationships. COI may be conservative for detecting rates of contemporary gene flow, but is expected to contain information content, due to frequent concordance with microsatellite data in the literature [28, 31]. Talbot et al. [31] used both microsatellites and COI to study genetic diversity within a bat ectoparasite species, and while COI exhibited less genetic variation among individuals with different host associations than microsatellites, they determined this was not due to any lack of ability in COI to detect differentiation, as among other populations, microsatellites showed less variation than COI. Jossart et al. [28] found comparable results between COI and the microsatellites when investigating intraspecific population genetic structure. COI sequences of species having variability in this marker, and which are separate in COI profile from related species, are suitable for use in this pipeline.

This script is also currently suitable for any geographic region(s), and a diverse selection of animal taxa, including terrestrial, marine, and freshwater groups from taxa such as arthropods, fish, and mammals. The pipeline also has the potential to be adapted to plant species, though with further consideration into the molecular markers and biological traits used. We recognize that some cases of hybrids or recently diverged species may be lumped together when using mitochondrial DNA, but we suggest that a large-scale analysis with large sample size is expected generally to reveal robust trends, a proposition that can be tested in future analyses. The other variables in the pipeline and the traits chosen for the analysis can be adjusted in order to suit the goals of the study and the study taxa.

### 4.4 Possible Extensions

In this study we presented an analysis framework to combine large-scale molecular estimates of genetic differentiation based on open-access molecular data of multiple species with species trait data. These modules provide a solid basis upon which other researchers can build and further develop to suit their own needs. The modules can be used together as a whole pipeline, as done in this analysis, or can be used individually and customized for the user’s needs. Modules such as those that involve specific data, such as the data loading and filtering modules, can be replaced or adapted to deal with different data sets, molecular markers, geographic regions, and taxa. The key modules of the pipeline are those that define regions/polygons and calculate and compare population genetic structure measures to biological traits and other important factors. By adapting or replacing other modules, these modules can be applied to a variety of taxa, geographic regions, and traits.

The pipeline can be expanded to include a variety of different data types and answer other kinds of biological questions. The pipeline can be used for de novo phylogenetics; however, a more complex clustering model should be used if users are planning to perform phylogenetics as part of their work. These modules can also be used with phylogeographic and speciation models. The results we obtained could be investigated further by including more statistical analysis. The idea of population stability and expansion could be investigated further, and analyses other than mismatch could be explored. There have been some critiques of mismatch analyses because there are many variables and opportunities to introduce errors [104]. These models can be influenced by the mutation rates, genes being investigated, sample sizes, and population characteristics [104]. Despite these critiques, mismatch distributions have proven useful when studying population genetic structure [36], and were worth including in our study as they provided additional insight into the populations being investigated.

Further research and consideration should be done into the method and how it suits the data and study goals if further research is to be done into population expansion of Diptera species.

Another possible expansion could be to examine whether indicators of population genetic structure have any relationship with the species name. As BINs were used here, and some BINs contain multiple species names, this could have resulted in some grouping together of closely related species.

The flexibility of this pipeline also allows for the inclusion of other types of data, such as climatic data and other types of molecular markers, such as cytochrome b and other mitochondrial markers. Small adjustments would still need to be made, as some functions in the filtering step, like those from coil version 1.2.3 [55], are built for COI only, and this could be replaced by steps to check for stop codons (in case of protein-coding genes) and anomalous sequences in the marker of choice. For markers such as microsatellites, more extensive adaptation would be needed. If a length-variable genetic region, e.g., ITS, is going to be used, then the trimming of the sequence to the reference step will need to be altered. The sequence may need to be anchored at the ends in more conserved regions, or a long sequence would need to be used as the reference. The pipeline can also be expanded to include multi-marker data. For our study, we were limited by the trait data available in the literature for our study taxa, as well as by the geographic regions that were sampled and barcoded. Further sampling and barcoding would allow for a more complete picture of fly species population genetic structure. For future research, the modules can be expanded to include other geographic regions, molecular markers, taxa, and traits to suit the requirements and needs of the researchers. Data from other databases can also be included, such as GBIF.

The modules provide several options to account for datasets of differing sizes, but some analyses can still be computationally demanding. Another possible expansion of the pipeline would be to adapt the script to be run-on high-performance computing platforms. If the dataset is extremely large, future researchers may consider this option rather than running the modules on their individual desktop machines. Future researchers could also make use of parallel processing.

### 4.5 Concluding Remarks

In conclusion, we were able to produce an efficient, reproducible, and publicly available pipeline by selecting and combining modules and functionality for determining indicators of population genetic structure and comparing these measures across species possessing different biological traits. Not only were valuable modules developed and made available for future researchers, but biological conclusions were also drawn regarding the population genetic structure of Diptera species in Canada. The hypothesis that indicators of realized dispersal rates will be influenced by the traits of organisms was supported, and traits such as larval diet were shown to be related to the levels of genetic differentiation between populations in Canada. In addition, evidence was found for isolation by distance, and a positive relationship between the population genetic structure measures and latitude and longitude was found. As our climate and world change, it is important to understand processes such as dispersal and the factors influencing them. By exploring the population genetic structure of species currently inhabiting Arctic and Northern regions, we can better understand how they will be impacted by our changing climate. By developing programming pipelines and workflows and making these accessible, we provide the opportunity for our work to be expanded on and provide tools that can help other scientists answer new questions.

## Supporting information

The full clustering analyses results are outlined in S1 Appendix.

The measures for each individual species are included as a csv file in S2 Appendix.

## Acknowledgements

SJA acknowledges financial support from the Natural Sciences and Engineering Research Council of Canada; the Government of Canada through Genome Canada and Ontario Genomics; the Ontario Ministry of Economic Development, Job Creation and Trade; and Food from Thought: Agricultural Systems for a Healthy Planet Initiative program funded by the Government of Canada through the Canada First Research Excellence Fund. The authors thank Ayesha Ali for her helpful feedback and insight on the project. They thank Tyler Elliott for testing the modules and providing important feedback on the code. They thank Jessica Castellanos-Labarcena for assistance with the geographic analysis in the early stages of the code and feedback on the manuscript. They also thank Matthew Orton, Jacqueline May, and Mehra Balsara, whose scripts were referenced for the pipeline. They are grateful to the researchers who share their data on public databases such as the Barcode of Life Data Systems and the Global Biodiversity Information Facility, and to all the researchers and developers who contribute to R programming packages and documentation.

## Supporting Information Captions

S1 Appendix: Results of 4% Clustering Analysis.

S2 Appendix: Population structure measures for each individual species in the Canada Analysis.

