## Supplementary material for "Novel pipeline for large-scale comparative population genetics Pipeline for population genetics": The full clustering analyses results are outlined in S1 Appendix.

Supporting Information

S1 Appendix: Results of 4% Clustering Analysis

Population Genetic Structure Measures

The full analysis was completed using a 4% clustering threshold and the single-linkage method instead of BINs (Table A1a). Both the clustering threshold and method can be adjusted by the user. 18 clusters across two geographic regions were included in the analysis for Greenland. The results reflected those of the BIN analysis. All of the Nei F_ST_ values for the clusters were low, showing low levels of genetic differentiation. The results for Hedrick’s G_ST_ and Jost’s D were more variable, with five clusters having a higher value (above 0.5) and 13 having a lower value for both measures. The AMOVA analysis revealed low levels of variation between the regions for all clusters. From the figures produced by the mismatch distribution analysis, 11 clusters’ populations appeared to be unimodal, and seven appeared to be bimodal. However, the dip test statistic indicated that there was no significant multimodality.

For the Canada dataset, 602 clusters across 32 regions/polygons were included (Table A1b). The results were similar to those observed in the original BIN-based analysis. Most Nei F_ST_ values were below 0.1, indicating low levels of genetic differentiation. Most Hedrick G_ST_ and Jost’s D values were above 0.8, indicating higher levels of genetic differentiation. The AMOVA for most clusters revealed low levels of variation between regions, with most phi values being below 0.1. The mismatch analysis revealed few significant signs of multimodality, suggesting that the populations are still expanding.

Comparison Across Traits

For the Greenland clustered analysis, eight were terrestrial, six were aquatic, and four were semi-aquatic. For larval diet, five were detritus and algae feeding, two were leaf/root/stem feeding, two were mycophagous and saprophagous, one was parasitic, five were saprophagous, two were predaceous, and one was leaf/root/stem feeding and detritus and algae feeding. For adult diet, one was leaf/root/stem feeding, two were nectar/pollen/honeydew feeding, one was predaceous, 13 were non-feeding, and one was both nectar/pollen/honeydew feeding and saprophagous. Like the BIN analysis, there was no significant relationship or trend in either the ANOVA or PGLS analysis. The traits and measures were also compared to the range size of each clusters. For both the BIN and cluster analysis, there were no significant relationships between range size and population genetic structure measures or traits for either the ANOVA or PGLS analysis.

For the Canada analysis, 142 clusters were aquatic, 47 were semi-aquatic, and 413 were terrestrial. For adult diet, one cluster was detritus and algae feeding, one was kleptoparasitic, 16 were leaf/root/stem feeding, 175 were nectar/pollen/honeydew feeding, three were parasitic, one was polyphagous, 67 were predaceous, 20 were saprophagous, two were both nectar/pollen/honeydew feeding and kleptoparasitic, four were nectar/pollen/honeydew feeding and mycophagous, 39 were nectar/pollen/honeydew feeding and parasitic, eight were nectar/pollen/honeydew feeding and predaceous, 64 were nectar/pollen/honeydew feeding and saprophagous, one was nectar/pollen/honeydew feeding, saprophagous and parasitic, 187 were nonfeeding and 13 had unclear diets. For larval diet, 100 clusters were detritus and algae feeding, one was kleptoparasitic, 63 were leaf/root/stem feeding, 34 were mycophagous, two were parasitic, 38 were parasitoid, 137 were predaceous, 175 were saprophagous, three were detritus and algae feeding and predaceous, 10 were leaf/root/stem feeding and detritus and algae feeding, three were leaf/root/stem feeding and mycophagous, five were leaf/root/stem feeding and saprophagous, one was leaf/root/stem feeding or detritus and algae feeding, seven were leaf/root/stem feeding or saprophagous, 10 were leaf/root/stem feeding, mycophagous and saprophagous, three were mycophagous and saprophagous, one was predaceous and saprophagous, three were predaceous or parasitoids, four were predaceous or saprophagous, and two had unclear diets. The relationship with habitat did not differ from the BIN analysis (Table A2). However, there was no longer a relationship with diet for Jost’s D and G_ST_. The F_ST_ results remained consistent for clusters with a kleptoparasitic larval diet and a leaf/root/stem feeding diet. An additional significant positive relationship was found with clusters that had a saprophagous larval diet, a larval diet that was a mix of leaf/root/stem feeding, mycophagous and saprophagous, and a larval diet that was predaceous or saprophagous.

Unlike the BIN analysis, no relationship was found between indicators of population genetic structure and distance. However, F_ST_ was still positively associated with both latitude and longitude. The multiple regression using the linearized measures only revealed a significant relationship with distance and F_ST_, and the relationship with latitude and longitude also persisted. There was also a relationship between indicators of population genetic structure and the number of cells in which the cluster was found and the number of records of each cluster. As the number of cells increased, so did the genetic differentiation. Oppositely, as the number of records increased, the genetic differentiation decreased.

S1 Table: Table outlining the population genetic structure measures using clustering at a 4% distance threshold from a) Greenland and b) Canada.

a) Greenland

| Cluster | Species Name | Region/Cell | F_ST_ | G_ST_ | Jost’s D | AMOVA Phi Value | AMOVA p-value | Dip Test Value | Dip Test p-value |
| --- | --- | --- | --- | --- | --- | --- | --- | --- | --- |
| 11 | *Spilogona sanctipauli* | 7721, 7478 | 0.011 | 0.18 | 0.16 | -0.0017 | 0.46 | 0.029 | 0.99 |
| 14 | *Phytomyza puccinelliae* | 7721, 7478 | 0.0098 | 0.28 | 0.27 | 0.006 | 0.2 | 0.033 | 1 |
| 20 | *Limnophyes ninae* | 7721, 7478 | 0.0062 | 0.36 | 0.35 | 0.0065 | 0.014 | 0.086 | 0.78 |
| 23 | *Limnophyes pumilio* | 7721, 7478 | 0.025 | 0.89 | 0.89 | 0.038 | 0.001 | 0.05 | 0.99 |
| 24 | *Limnophyes brachytomus* | 7721, 7478 | 0.0015 | 0.1 | 0.099 | 0.00062 | 0.22 | 0.057 | 0.97 |
| 59 | *Lycoriella riparia* | 7721, 7478 | 0.0084 | 0.23 | 0.22 | -0.0051 | 0.58 | 0.1 | 0.45 |
| 65 | *Lycoriella flavipeda* | 7721, 7478 | 0.0027 | 0.16 | 0.15 | 0.0047 | 0.001 | 0.016 | 1 |
| 69 | *Camptochaeta delicata* | 7721, 7478 | 0.0081 | 0.25 | 0.24 | 0.0087 | 0.031 | 0.025 | 1 |
| 83 | *Corynoneura arctica* | 7721, 7478 | 0.0072 | 0.3 | 0.29 | 0.011 | 0.001 | 0.061 | 0.71 |
| 91 | *Nephrotoma lundbecki* | 7721, 7478 | 0.029 | 0.38 | 0.34 | 0.024 | 0.13 | 0.078 | 0.9 |
| 93 | *Cricotopus patens* | 7721, 7478 | 0.05 | 0.75 | 0.72 | 0.054 | 0.058 | 0.12 | 0.24 |
| 103 | *Smittia extrema* | 7721, 7478 | 0.016 | 0.47 | 0.45 | 0.028 | 0.001 | 0.044 | 1 |
| 113 | *Megaselia cirriventris* | 7721, 7478 | 0.023 | 0.69 | 0.68 | 0.041 | 0.001 | 0.033 | 1 |
| 136 | *Paraphaenocladius impensus* | 7721, 7478 | 0.0061 | 0.2 | 0.19 | 0.01 | 0.002 | 0.025 | 1 |
| 147 | *Diamesa arctica* | 7721, 7478 | 0.011 | 0.78 | 0.78 | 0.0083 | 0.018 | 0.098 | 0.41 |
| 149 | *Diamesa chorea* | 7721, 7478 | 0.014 | 0.23 | 0.2 | 0.021 | 0.004 | 0.04 | 1 |
| 180 | *Drymeia groenlandica* | 7721, 7478 | 0.042 | 0.43 | 0.38 | 0.063 | 0.001 | 0.025 | 1 |
| 183 | *Protophormia terraenovae* | 7721, 7478 | 0.037 | 0.84 | 0.83 | 0.053 | 0.008 | 0.038 | 1 |

b) Canada

| Measure | Max | Min |
| --- | --- | --- |
| Global F_ST_ | 0.4 | 0.002 |
| Global G_ST_ | 1 | 0.032 |
| Jost’s D | 1 | 0.019 |
| AMOVA Phi Value | 0.58 | -0.03 |
| AMOVA p-value | 1 | 0.001 |
| Dip Test Value | 0.5 | 0.031 |
| Dip Test p-value | 1 | 0.004 |

S2 Table: Table showing the results from the multiple regression comparing population genetic structure measures to biological traits and other important factors for Canada using a 4% clustering threshold for a) Jost’s D, b) G_ST_ and c) F_ST_. Significant values are bolded.

a) Jost’s D

| Trait/Factors | t-value | p-value |
| --- | --- | --- |
| Semi-Aquatic | -0.93 | 0.35 |
| Terrestrial | -1.72 | 0.086 |
| Adult Diet: Kleptoparasitic | 0.47 | 0.64 |
| Adult Diet: Leaf/Root/Stem Feeding | -0.55 | 0.58 |
| Adult Diet: Nectar/Pollen/Honeydew Feeding | -0.31 | 0.76 |
| Adult Diet: Nectar/Pollen/Honeydew Feeding and Kleptoparasitic | -0.95 | 0.34 |
| Adult Diet: Nectar/Pollen/Honeydew Feeding and Mycophagous | -0.52 | 0.61 |
| Adult Diet: Nectar/Pollen/Honeydew Feeding and Parasitic | -0.3 | 0.77 |
| Adult Diet: Nectar/Pollen/Honeydew Feeding and Predaceous | -0.11 | 0.91 |
| Adult Diet: Nectar/Pollen/Honeydew Feeding and Saprophagous | -0.46 | 0.64 |
| Adult Diet: Nectar/Pollen/Honeydew Feeding, Saprophagous and Parasitic | -1.26 | 0.21 |
| Adult Diet: Non-Feeding | -0.35 | 0.73 |
| Adult Diet: Parasitic | -0.98 | 0.33 |
| Adult Diet: Polyphagous | -0.13 | 0.89 |
| Adult Diet: Predaceous | -0.24 | 0.81 |
| Adult Diet: Saprophagous | -0.32 | 0.75 |
| Adult Diet: Unclear | -0.16 | 0.87 |
| Larval Diet: Detritus and Algae Feeding and Predaceous | 0.46 | 0.64 |
| Larval Diet: Kleptoparasitic | 0.91 | 0.36 |
| Larval Diet: Leaf/Root/Stem Feeding | 1.44 | 0.15 |
| Larval Diet: Leaf/Root/Stem Feeding and Detritus and Algae Feeding | 0.63 | 0.53 |
| Larval Diet: Leaf/Root/Stem Feeding and Mycophagous | -0.88 | 0.38 |
| Larval Diet: Leaf/Root/Stem Feeding and Saprophagous | -1.61 | 0.11 |
| Larval Diet: Leaf/Root/Stem Feeding or Detritus and Algae Feeding | 0.62 | 0.54 |
| Larval Diet: Leaf/Root/Stem Feeding or Saprophagous | -0.82 | 0.41 |
| Larval Diet: Leaf/Root/Stem Feeding, Mycophagous and Saprophagous | -0.45 | 0.65 |
| Larval Diet: Mycophagous | 0.49 | 0.62 |
| Larval Diet: Mycophagous and Saprophagous | -0.82 | 0.41 |
| Larval Diet: Parasitic | 0.31 | 0.76 |
| Larval Diet: Parasitoid | -0.99 | 0.32 |
| Larval Diet: Predaceous | -0.12 | 0.9 |
| Larval Diet: Predaceous and Saprophagous | 0.45 | 0.65 |
| Larval Diet: Predaceous or Parasitoid | 1.53 | 0.13 |
| Larval Diet: Predaceous or Saprophagous | 0.25 | 0.8 |
| Larval Diet: Saprophagous | -0.46 | 0.65 |
| Larval Diet: Unclear | 0.84 | 0.4 |
| Number of Cells | 1.37 | 0.17 |
| Number of Records | -0.47 | 0.64 |
| Max Distance | 1.72 | 0.087 |
| Longitude Coordinate | 0.58 | 0.56 |
| Latitude Coordinate | 1.49 | 0.14 |

b) G_ST_

| Trait/Factors | t-value | p-value |
| --- | --- | --- |
| Semi-Aquatic | -0.92 | 0.36 |
| Terrestrial | -1.7 | 0.09 |
| Adult Diet: Kleptoparasitic | 0.47 | 0.64 |
| Adult Diet: Leaf/Root/Stem Feeding | -0.56 | 0.57 |
| Adult Diet: Nectar/Pollen/Honeydew Feeding | -0.29 | 0.77 |
| Adult Diet: Nectar/Pollen/Honeydew Feeding and Kleptoparasitic | -0.91 | 0.37 |
| Adult Diet: Nectar/Pollen/Honeydew Feeding and Mycophagous | -0.49 | 0.62 |
| Adult Diet: Nectar/Pollen/Honeydew Feeding and Parasitic | -0.3 | 0.77 |
| Adult Diet: Nectar/Pollen/Honeydew Feeding and Predaceous | -0.004 | 1 |
| Adult Diet: Nectar/Pollen/Honeydew Feeding and Saprophagous | -0.45 | 0.65 |
| Adult Diet: Nectar/Pollen/Honeydew Feeding, Saprophagous and Parasitic | -1.052 | 0.29 |
| Adult Diet: Non-Feeding | -0.33 | 0.74 |
| Adult Diet: Parasitic | -0.95 | 0.34 |
| Adult Diet: Polyphagous | -0.14 | 0.9 |
| Adult Diet: Predaceous | -0.24 | 0.81 |
| Adult Diet: Saprophagous | -0.31 | 0.76 |
| Adult Diet: Unclear | -0.14 | 0.89 |
| Larval Diet: Detritus and Algae Feeding and Predaceous | 0.49 | 0.62 |
| Larval Diet: Kleptoparasitic | 0.89 | 0.37 |
| Larval Diet: Leaf/Root/Stem Feeding | 1.49 | 0.14 |
| Larval Diet: Leaf/Root/Stem Feeding and Detritus and Algae Feeding | 0.62 | 0.54 |
| Larval Diet: Leaf/Root/Stem Feeding and Mycophagous | -0.91 | 0.36 |
| Larval Diet: Leaf/Root/Stem Feeding and Saprophagous | -1.6 | 0.11 |
| Larval Diet: Leaf/Root/Stem Feeding or Detritus and Algae Feeding | 0.61 | 0.55 |
| Larval Diet: Leaf/Root/Stem Feeding or Saprophagous | -0.72 | 0.47 |
| Larval Diet: Leaf/Root/Stem Feeding, Mycophagous and Saprophagous | -0.43 | 0.67 |
| Larval Diet: Mycophagous | 0.41 | 0.68 |
| Larval Diet: Mycophagous and Saprophagous | -0.79 | 0.43 |
| Larval Diet: Parasitic | 0.33 | 0.74 |
| Larval Diet: Parasitoid | -0.98 | 0.33 |
| Larval Diet: Predaceous | -0.07 | 0.94 |
| Larval Diet: Predaceous and Saprophagous | 0.46 | 0.65 |
| Larval Diet: Predaceous or Parasitoid | 1.5 | 0.13 |
| Larval Diet: Predaceous or Saprophagous | 0.35 | 0.73 |
| Larval Diet: Saprophagous | -0.46 | 0.65 |
| Larval Diet: Unclear | 0.82 | 0.41 |
| Number of Cells | 1.49 | 0.14 |
| Number of Records | -0.59 | 0.55 |
| Max Distance | 1.7 | 0.089 |
| Longitude Coordinate | 0.62 | 0.53 |
| Latitude Coordinate | 1.54 | 0.12 |

c) F_ST_

|  | Trait/Factors | t-value | p-value |
| --- | --- | --- | --- |
| Semi-Aquatic | | **-2.28** | **0.023** |
| Terrestrial | | -0.99 | 0.32 |
| Adult Diet: Kleptoparasitic | | 0.08 | 0.94 |
| Adult Diet: Leaf/Root/Stem Feeding | | -0.36 | 0.72 |
| Adult Diet: Nectar/Pollen/Honeydew Feeding | | 0.12 | 0.91 |
| Adult Diet: Nectar/Pollen/Honeydew Feeding and Kleptoparasitic | | -0.37 | 0.71 |
| Adult Diet: Nectar/Pollen/Honeydew Feeding and Mycophagous | | -0.47 | 0.64 |
| Adult Diet: Nectar/Pollen/Honeydew Feeding and Parasitic | | -0.11 | 0.91 |
| Adult Diet: Nectar/Pollen/Honeydew Feeding and Predaceous | | 0.93 | 0.35 |
| Adult Diet: Nectar/Pollen/Honeydew Feeding and Saprophagous | | 0.56 | 0.58 |
| Adult Diet: Nectar/Pollen/Honeydew Feeding, Saprophagous and Parasitic | | 0.56 | 0.58 |
| Adult Diet: Non-Feeding | | 0.2 | 0.84 |
| Adult Diet: Parasitic | | -0.069 | 0.94 |
| Adult Diet: Polyphagous | | -0.84 | 0.4 |
| Adult Diet: Predaceous | | 0.44 | 0.66 |
| Adult Diet: Saprophagous | | -0.036 | **0.97** |
| Adult Diet: Unclear | | -0.001 | 1 |
| Larval Diet: Detritus and Algae Feeding and Predaceous | | 0.041 | 0.97 |
| Larval Diet: Kleptoparasitic | | **2.75** | **0.0062** |
| Larval Diet: Leaf/Root/Stem Feeding | | **3.69** | **0.00024** |
| Larval Diet: Leaf/Root/Stem Feeding and Detritus and Algae Feeding | | -0.93 | 0.35 |
| Larval Diet: Leaf/Root/Stem Feeding and Mycophagous | | 0.23 | 0.82 |
| Larval Diet: Leaf/Root/Stem Feeding and Saprophagous | | 1.61 | 0.11 |
| Larval Diet: Leaf/Root/Stem Feeding or Detritus and Algae Feeding | | -0.22 | 0.83 |
| Larval Diet: Leaf/Root/Stem Feeding or Saprophagous | | 1.91 | 0.056 |
| Larval Diet: Leaf/Root/Stem Feeding, Mycophagous and Saprophagous | | **2.36** | **0.019** |
| Larval Diet: Mycophagous | | -0.39 | 0.7 |
| Larval Diet: Mycophagous and Saprophagous | | 0.66 | 0.51 |
| Larval Diet: Parasitic | | 0.16 | 0.87 |
| Larval Diet: Parasitoid | | 1.22 | 0.22 |
| Larval Diet: Predaceous | | 1.15 | 0.25 |
| Larval Diet: Predaceous and Saprophagous | | -0.56 | 0.58 |
| Larval Diet: Predaceous or Parasitoid | | -0.98 | 0.33 |
| Larval Diet: Predaceous or Saprophagous | | **1.98** | **0.049** |
| Larval Diet: Saprophagous | | **2.008** | **0.045** |
| Larval Diet: Unclear | | -0.76 | 0.45 |
| Number of Cells | | **3.41** | **0.00071** |
| Number of Records | | **-3.33** | **0.00092** |
| Max Distance | | 0.68 | 0.49 |
| Longitude Coordinate | | **2.83** | **0.0048** |
| Latitude Coordinate | | **6.21** | **1.03e-09** |
